# Mapping cells through time and space with moscot

**DOI:** 10.1101/2023.05.11.540374

**Authors:** Dominik Klein, Giovanni Palla, Marius Lange, Michal Klein, Zoe Piran, Manuel Gander, Laetitia Meng-Papaxanthos, Michael Sterr, Aimée Bastidas-Ponce, Marta Tarquis-Medina, Heiko Lickert, Mostafa Bakhti, Mor Nitzan, Marco Cuturi, Fabian J. Theis

**Affiliations:** Institute of Computational Biology, Helmholtz Center Munich, Germany; Department of Mathematics, Technical University of Munich, Germany; TUM School of Life Sciences Weihenstephan, Technical University of Munich, Germany; Department of Biosystems Science and Engineering, ETH Zürich, Basel, Switzerland; School of Computer Science and Engineering, The Hebrew University of Jerusalem, Israel; Google Research; Apple; Institute of Diabetes and Regeneration Research, Helmholtz Center Munich; German Center for Diabetes Research, Neuherberg, Germany; School of Medicine, Technical University of Munich, Germany; Racah Institute of Physics, The Hebrew University of Jerusalem, Israel; Faculty of Medicine, The Hebrew University of Jerusalem, Israel

## Abstract

Single-cell genomics technologies enable multimodal profiling of millions of cells across temporal and spatial dimensions. Experimental limitations prevent the measurement of all-encompassing cellular states in their native temporal dynamics or spatial tissue niche. Optimal transport theory has emerged as a powerful tool to overcome such constraints, enabling the recovery of the original cellular context. However, most algorithmic implementations currently available have not kept up the pace with increasing dataset complexity, so that current methods are unable to incorporate multimodal information or scale to single-cell atlases. Here, we introduce multi-omics single-cell optimal transport (moscot), a general and scalable framework for optimal transport applications in single-cell genomics, supporting multimodality across all applications. We demonstrate moscot’s ability to efficiently reconstruct developmental trajectories of 1.7 million cells of mouse embryos across 20 time points and identify driver genes for first heart field formation. The moscot formulation can be used to transport cells across spatial dimensions as well: To demonstrate this, we enrich spatial transcriptomics datasets by mapping multimodal information from single-cell profiles in a mouse liver sample, and align multiple coronal sections of the mouse brain. We then present moscot.spatiotemporal, a new approach that leverages gene expression across spatial and temporal dimensions to uncover the spatiotemporal dynamics of mouse embryogenesis. Finally, we disentangle lineage relationships in a novel murine, time-resolved pancreas development dataset using paired measurements of gene expression and chromatin accessibility, finding evidence for a shared ancestry between delta and epsilon cells. Moscot is available as an easy-to-use, open-source python package with extensive documentation at https://moscot-tools.org.

## Introduction

Single-cell genomics technologies have revolutionized our understanding of the dynamics of cellular differentiation and tissue organization. Single-cell assays like scRNA-seq or scATAC-seq^1^ profile the molecular state of individual cells at unprecedented resolution, and spatial assays like MERFISH^2^ or Stereo-seq^3^ recover the spatial organization of single-cell profiles. However, these experiments are cell-destructive, and capture only a subset of molecular information, often yielding an incomplete picture of the cell state variation in the biological system of interest. As a result, reconstructing cellular trajectories with time-series scRNA-seq requires matching unaligned snapshot datasets captured at different time points^4, 5^. Similarly, in the analysis of spatial transcriptomics datasets, where measurements often contain sections from different samples or areas of the tissue, it is desirable to align them to a common reference template, and map additional molecular information from external datasets such as reference atlases^6, 7^.

Previous work addressed such mapping and alignment problems using optimal transport (OT) theory, an area of applied mathematics concerned with comparing and aligning probability distributions^8, 9^. For example, OT has been instrumental in delineating cellular reprogramming processes towards induced pluripotent stem cells (iPSCs) and mouse gastrulation^4, 10, 11^, reconstructing tissue architecture by enhancing spatial data with single-cell references^12^ and building common coordinate frameworks (CCFs) of a biological system by aligning spatial transcriptomics data^13^.

Despite the potential of OT-based methods to address mapping problems in single-cell genomics, their use faces three key challenges. First, current implementations of OT-based tools to map cells across time and space are geared towards uni-modal data and do not incorporate multimodal information. Second, OT methods currently used in single-cell genomics are computationally expensive; the original linear programming formulation of OT scales cubically in cell number. More recent approaches, grounded on entropic regularization, scale quadratically^14^ (or cubically for Gromov-Wasserstein extensions^8, 15^), similarly memory scales quadratically in cell number, which altogether prevents using GPUs and the application to emerging atlas-scale datasets^5, 16^. Third, existing OT-based tools are based on various OT backends and heterogeneous implementations^4, 12, 13, 17^, making it difficult to adapt or combine approaches towards new problems such as spatiotemporal mapping. In contrast, user-friendly and extensible application programming interfaces (APIs) accelerate and facilitate research, as exemplified by scVI-tools^18^ with its many extensions^19–21^.

Here, we present Multi-Omics Single-Cell Optimal Transport (moscot), a computational framework to solve mapping and alignment problems that arise in single-cell genomics and demonstrate its capabilities for temporal, spatial, and spatiotemporal applications. Moscot implements three fundamental design principles to overcome previous limitations. First, moscot supports multimodal data throughout the framework by exploiting joint cellular representations. Second, moscot improves scalability by adapting and demonstrating the applicability of recent methodological innovations^22–24^ to atlas-scale datasets. Third, moscot unifies previous single-cell applications of OT in the temporal and spatial domain and introduces a novel spatiotemporal application. All of this is achieved with a robust and intuitive API that interacts with the broader scverse^25, 26^ ecosystem.

Moscot’s improved scalability enables us to map 1.7 million cells across an atlas of mouse embryogenesis. Furthermore, we illustrate that moscot can be used to map multimodal CITE-seq^27^ information to high-resolution spatial readouts in the mouse liver as well as aligning large spatial transcriptomics sections of mouse brain samples. We introduce the concept of spatiotemporal mapping, which involves the integration of spatial coordinates and gene expression data. Leveraging a spatiotemporal atlas of mouse embryogenesis^3^, we demonstrate that incorporating spatial information enhances mapping accuracy, facilitates the transfer of high resolution cell type annotation, and enables the study of differentiation trajectories. Finally, we jointly profile gene expression and chromatin accessibility during mouse pancreatic development and apply moscot to formulate a novel hypothesis about cell ancestry of delta and epsilon cells. Moscot unlocks optimal transport for multi-view atlas-scale single-cell applications; we make it available as an easy-to-use, open-source python package with extensive documentation at https://moscot-tools.org.

## Results

### Moscot is a general and scalable framework to map cellular distributions across time and space

The core idea of the moscot framework is to translate biological mapping and alignment tasks into OT problems and solve them using a consistent set of algorithms. Moscot takes as input unpaired datasets, which can represent different time points, modalities, or spatial representations (Methods and Fig. 1a). Additionally, moscot accepts prior biological knowledge, such as cellular growth and death rates, to guide the mapping.

**Fig. 1.**
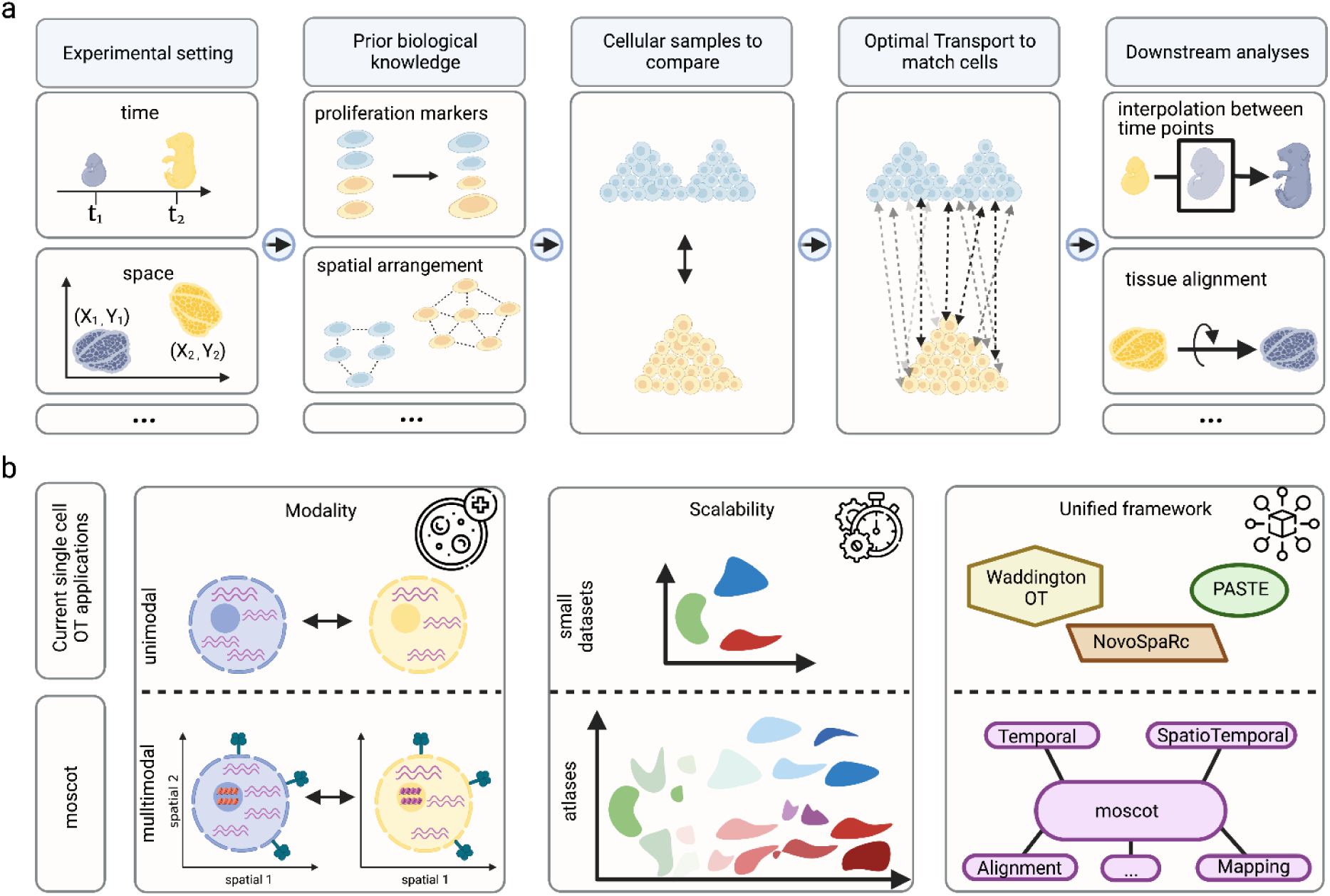
moscot enables efficient multimodal optimal transport across single-cell applications. **a.** Sketch of a generic optimal transport (OT) pipeline in single-cell genomics: experimental shifts (e.g., time points, spatial vs. dissociated) lead to disparate cell populations that must be mapped (**i**). Prior biological knowledge (e.g., proliferation rates, spatial arrangement) is often available and should be used to guide the mapping (**ii**). The mapping problem mathematically compares probability distributions over sampled cellular states (**iii**). OT provides a standardized way to solve the mapping problem and many of its variants (**iv**). Solving the mapping problem creates various downstream analysis opportunities; moscot supports many of these (**v**). **b.** moscot introduces three key innovations that unlock the full power of OT for problems: first, it supports multimodal data across all models. Second, overcomes previous scalability limitations to enable atlas-scale applications. Third, moscot is a unified framework with a consistent application programming interface (API) across biological problems; this enables consistent application of methods across different problems and fast generalization to new ones.

Importantly, using a notion of distance between the unpaired datasets, moscot solves an OT problem and returns a coupling matrix that probabilistically relates samples in each of the datasets. Equipped with that coupling matrix, moscot offers various application-specific downstream analysis functions, such as time point interpolation and tissue alignment (Methods and Fig. 1a). Chaining of coupling matrices allows modeling sequential time points or multiple spatial transcriptomics slides.

Moscot builds on three notions of OT to accommodate various biological problems. These OT-notions differ in how samples are related across datasets: Wasserstein-type (W-type)^8^ OT compares individual samples, i.e. one cell per dataset per comparison. Gromov-Wasserstein-type (GW-type)^15^ OT compares pairwise samples, i.e. two cells per dataset per comparison. Finally, Fused Gromov-Wasserstein-type (FGW-type)^28^ OT compares both individual and pairwise samples (Methods and Supplementary Note 1). Accordingly, W- and (F)GW-type OT can be used to map samples within and across molecular representations, including gene expression at different time points^4^ and spatial coordinate systems^13^, respectively.

To support multimodality throughout the framework, we define the cost of transporting cells using shared latent representations (Fig. 1b and Methods). Depending on the model, we obtain these representations by concatenating uni-modal latent representations or using joint latent space learning techniques^20, 29–32^. This approach generalizes to more than two modalities and additional temporal or spatial information.

We make moscot applicable to atlas-scale datasets by reducing compute time and memory consumption of W-, GW-, and FGW-type notions by orders of magnitude compared to prior OT-based tools (Fig. 1b, Methods and Supplementary Note 2). Specifically, we base moscot on Optimal Transport Tools^22^ (OTT), a scalable JAX^33^ implementation of OT algorithms that supports just-in-time compilation, on-the-fly evaluation of the cost function, and GPU acceleration (Methods). Where required by dataset size, we use recent methodological innovations^23, 24^ that constrain the coupling matrix to be low-rank, enabling linear time and memory complexity for W, GW, and FGW-type notions (Supplementary Note 3).

A unified API makes moscot easy to use and extend (Supplementary Fig. 1). In particular, a modular implementation enables using similar infrastructure for solving different biological problems (Supplementary Fig. 1 and Methods). Moscot models share interfaces with core packages like SCANPY^25, 26^ or AnnData^34^, use a small set of OT algorithms in the backend to compute coupling matrices, and reuse downstream analysis functions across applications.

### Recovering trajectories of mouse embryogenesis on a developmental atlas using moscot

Modeling cell-state trajectories for biological systems that are not in steady state requires time-course single-cell studies combined with computational analysis to infer cellular differentiation across time points. The popular Waddington OT^4^ (WOT) method solves the problem using W-type OT; however, it remains limited to uni-modal gene expression data and does not scale to hundreds of thousands of cells per time point, needed for numerous recent datasets. Thus, we introduce moscot.time. Our model inherits WOT’s popular cell growth- and death rate modeling, is applicable to multimodal data, scales to millions of cells and, like all trajectory inference methods in moscot, can be combined with downstream analyses such as CellRank^35^ (Methods).

We asked whether moscot.time’s improved scalability translates into a more faithful description of real biological systems. Thus, we applied our model to an atlas of early mouse development, containing almost 1.7 million cells across 20 time points spanning embryonic days (E) 3.5-13.5 (ref.^16^) (Fig. 2a and Methods). We first assessed whether we could use the previous state-of-the-art, WOT^4^, to analyze this dataset. We selected the E11.5 to E12.5 time point pair, containing over half a million cells, and benchmarked memory and compute time on subsets of increasing cell number (Fig. 2b, Methods and Supplementary Table 1). While moscot.time computed a coupling for all 275,000 cells at both time points using less than 10 GiB memory in approximately 3 hours, WOT ran out of memory as soon as we exceeded 100,000 cells, even though we provided 700 GiB server memory. In principle, moscot.time’s linear memory complexity allows it to process developmental atlases on a laptop, where WOT fails on a server (Methods and Fig. 2b).

**Fig. 2.**
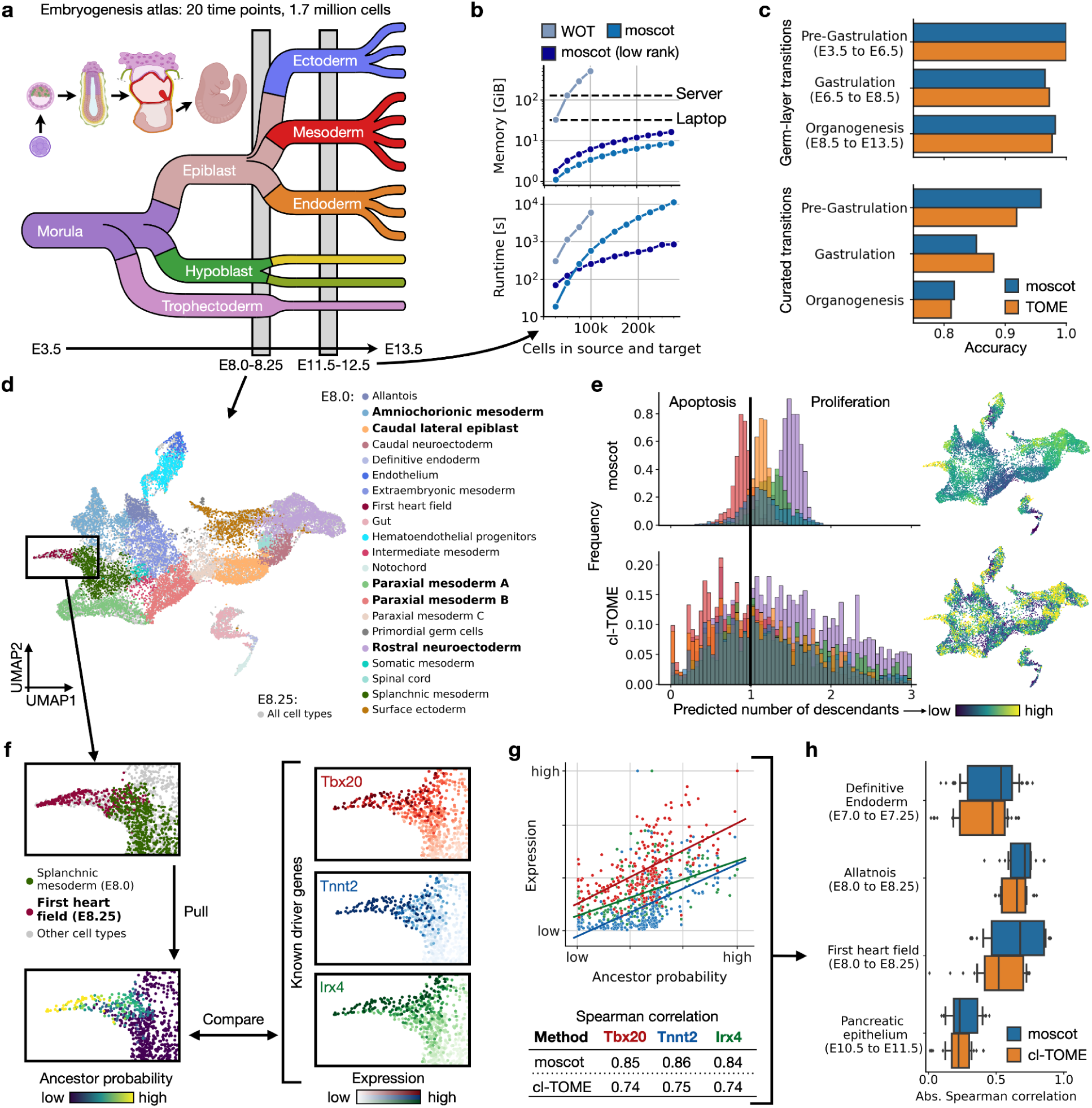
moscot faithfully reconstructs atlas-scale developmental trajectories. **a.** Schematic of the mouse embryogenesis atlas example^16^. **b.** Benchmark of peak memory consumption (top, on CPU) and compute time (bottom, on GPU) on the embryonic day (E) 11.5 and E12.5 time point pair (Methods and Supplementary Table 1). Waddington OT^4^ (WOT) compared with default moscot.time and low-rank^23^ moscot.time (rank 200) on 11 subsampled versions of the full dataset (Supplementary Note 3; WOT was run on CPU as it does not support GPU acceleration). **c.** Accuracy comparison between TOME^16^ and moscot.time in terms of germ layer and cell type transition scores by developmental stage (Methods and Supplementary Table 2). **d.** UMAP^96, 97^ projection of the E8.0/8.25 time point pair, colored by original cluster annotations. **e.** Growth rate estimates of moscot.time (top) and cl-TOME (bottom) for the bold E8.0 cell types in (d), as histograms (left) and UMAP projections (right). The black vertical bar denotes growth rate one. **f.** moscot.time’s ancestor probability for E8.25 first heart field cells (left) versus gene expression levels of known driver genes *Tbx20*^50^, *Tnnt2*^98^ and *Irx4*^99^ (right) (Methods). **g.** Quantification of comparison in (f) via Spearman correlation. Genes are colored as in (f), each dot denotes a cell, and lines indicate a linear data fit. **h.** Spearman correlation between ancestor probabilities and known driver-gene expression for moscot.time and cl-TOME (Methods and Supplementary Table 4).

As WOT does not scale to a dataset of this size, the authors of the developmental atlas^16^ devised a novel deterministic approach based on k-nearest neighbor (k-NN) matching, Trajectories Of Mammalian Embryogenesis (TOME). To investigate how moscot.time compares with TOME in terms of accuracy, we formulated two metrics that operate on the level of germ layers (i) and cell types (ii). These metrics encourage within-germ layer transitions (i), and transitions among cell types that have been reported to transition into one-another (ii) (Methods and Supplementary Table 2). In both metrics, moscot.time achieved comparable performance to TOME across all time points and developmental stages, even though TOME was designed specifically for this dataset (Fig. 2c).

To compare TOME and moscot.time at finer resolution, we focused on cellular growth- and death rates. As TOME only provides a cluster-level mapping, we extended the original approach to yield cell-level output with cell-level Trajectories Of Mammalian Embryogenesis (cl-TOME)

(Methods). Using the E8.0/8.25 pair of time points as an example, we mapped cells using moscot.time and cl-TOME (Fig. 2d). Across cell types, cl-TOME frequently assigned growth rates much smaller than one, predicting that over 19% of the population at this stage is apoptotic (Fig. 2e and Supplementary Table 3). Such a high death-rate represents an unrealistic scenario for embryonic development, where, beyond E7.0, the fraction of cells going through apoptosis is typically much below 10% (ref.^36, 37^). In contrast, for moscot.time, we were able to tune the apoptosis rate to lie in a biologically plausible range (Methods). Further, the growth rates predicted by moscot.time were more cell-type specific (Fig. 2e), and these results generalized to all other time points that contained sufficient cell numbers (Supplementary Figures 2-4).

Next, we were interested in comparing the linkages between early- and late cells predicted by either method. As moscot’s growth and death rates were biologically more plausible, we expected the learnt cell couplings to be more accurate, too. We first assessed this using the example of E8.25 first heart field cells, a population that emerges from the splanchnic mesoderm^38^. We used moscot.time and cl-TOME to compute ancestor probabilities; these quantify the likelihood of E8.0 cells to evolve toward E8.25 first heart field cells. To score the accuracy of these predictions, we compared ancestor probabilities with the expression of known driver genes for first heart field formation at E8.0 (Fig. 2f, Methods and Supplementary Table 3). We quantified the comparison using Spearman correlation (Fig. 2g). Indeed, we found that moscot.time consistently achieved higher absolute correlations, a result that generalized to three other cell types we investigated across early development (Fig. 2h and Supplementary Table 4).

Moscot’s scalability enabled us to apply OT to a single-cell atlas of mouse gastrulation^16^, obtain realistic growth- and death rates, as well as biologically more plausible couplings compared with k-NN based approaches^16^.

### Mapping and aligning spatial samples with moscot

Spatial omics technologies enable profiling thousands of cells in their tissue context. The analysis of such data requires methods that are able to integrate datasets across molecular layers and spatial coordinate systems. OT has proven useful to tackle these problems, most notably novoSpaRc^12^ for gene expression mapping and PASTE^13^ for alignment and registration of spatial transcriptomics datasets. Moscot implements both applications of OT, in the moscot.space *Mapping* and *Alignment* problems, leveraging scalable implementations and more performant algorithms (Methods).

Image-based spatial transcriptomics data is often limited in the number of measured genes (hundreds to a few thousands^7^). The *Mapping* problem learns a map between dissociated single-cell profiles and their spatial organization. Once the map is learned, it can be used to transfer gene expression or additional multimodal profiles to spatial coordinates (Fig. 3.a). We implemented the mapping application in moscot.space using an FGW-type problem; such a formulation enables us to incorporate cellular similarities across molecular representations as well as physical cell-distances into the optimization problem (Methods).

**Fig. 3.**
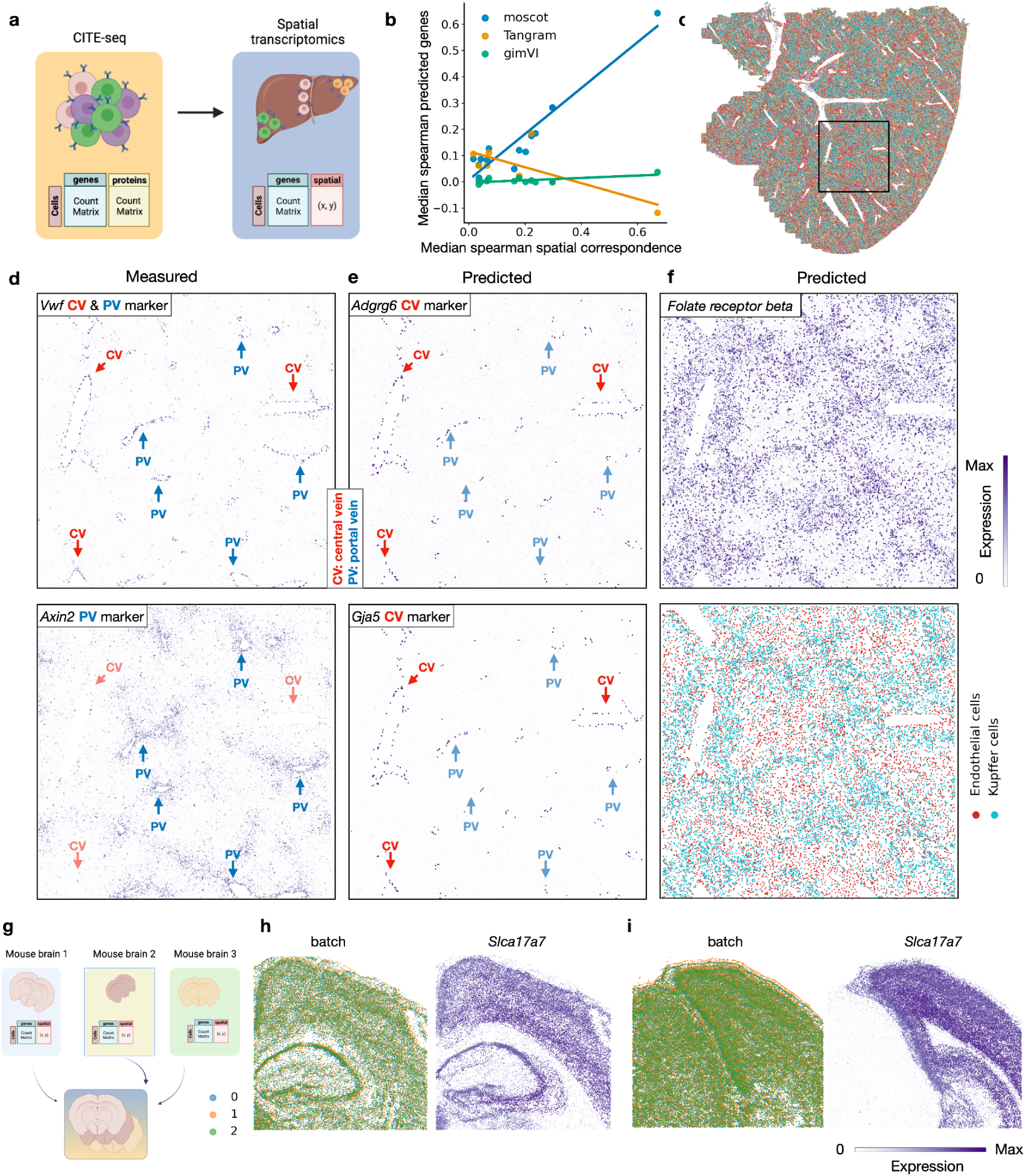
moscot enables multimodal mapping and alignment of spatial transcriptomics data. **a.** Schematic of the moscot application of multimodal mapping. A multimodal single-cell reference dataset can be mapped onto a spatial dataset in order to project multimodal features to spatial coordinates. **b.** Spatial correspondence is associated with prediction accuracy in moscot. The median Spearman correlation (spatial correspondence) is plotted on the x-axis against the median Spearman correlation across predicted gene expression for the spatial gene expression mapping task (y-axis). Each point corresponds to a different dataset (same datasets as the ones used in the benchmark, although only those 11 yielding a valid result across all seeds for all methods are considered) and the color represents the method. The overlaid line segment is the result of a linear regression for each method. **c.** Spatial plot of the liver MERSCOPE section with annotations mapped from the CITE-seq dataset, the legend for the cell type annotation can be found in Supplementary Fig. 6. The region surrounded by the solid line corresponds to the cropped tiles in the subsequent panels. **d**. Spatial visualization of measured gene expression values for *Vwf* (endothelial cell marker) and *Axin2* (hepatocytes and endothelial marker). *Vwf* is used to identify all epithelial cells that define the boundaries of CV and PV. *Axin2* is used as a positive marker for PV. **e.** Predicted gene expression for *Adgrg6* and *Gja5*, which are known endothelial cells markers for CV. **f.** The moscot mapping can also be used to impute protein expression and transfer cell type annotations. The top panel shows the predicted expression of *Folate receptor beta*, a marker for Kupffer cells. The bottom plot visualizes the imputed cell types for Kupffer cells and endothelial cells. **g.** Schematic of the moscot application of spatial alignment. Sections from multiple samples can be aligned to a common reference sample. **h.** Visualization of a tile of the spatial sections of Vizgen MERSCOPE mouse brain for sections 1 colored by batch (left) and by expression of *Slc17a7* (right). **i.** Visualization of a tile of the spatial sections of Vizgen MERSCOPE mouse brain for sections 2 colored by batch (left) and by expression of *Slc17a7* (right).

To assess the performance of moscot.space to map dissected single-cells, we evaluated the methods moscot.space *Mapping* tool, and two state-of-the-art methods, Tangram^39^ and gimVI^40^, according to a recent benchmark^41^, on datasets provided by Li et al^41^. We assessed the quality of the mapping by computing correlation of held out genes in spatial coordinates (Methods). Moscot consistently outperformed the other methods across 14 datasets of various technologies used in the benchmark (Supplementary Fig. 5).

Furthermore, for each dataset we quantified “spatial correspondence”, a measure of correlation between gene expression similarity and distances in physical coordinates, as originally proposed by Nitzan et al.^12^ (Methods). A spatial transcriptomics dataset has high spatial correspondence if nearby cells have similar gene expression (Supplementary Fig. 5). Moscot showed a positive correlation between spatial correspondence and accuracy (Fig. 3b), indicating that it is able to leverage spatial associations between distances in gene expression and physical space. Nevertheless, even when spatial correspondence is low, moscot still outperformed the competing methods.

Having demonstrated the power of moscot to map gene expression to space, we used moscot to map multimodal single-cell profiles to their spatial context. This is of particular interest as spatial transcriptomics technologies are mostly limited to gene expression measurements^6, 7^. We considered two datasets, collected in different labs: a CITE-seq dataset of ∼91k cells of the mouse liver^42^ and a spatial transcriptomics section consisting of ∼367k cells measured with Vizgen MERSCOPE^43^ (Fig. 3c). We map the annotation from the CITE-seq dataset as no cell type annotation was provided in the original data (Supplementary Fig. 6). We then solved a FGW-type problem with moscot which allowed us to incorporate gene expression, protein and spatial information to recover the spatial organization of the proteins (Methods). Using any of the competing methods is not feasible due to prohibitive time or memory complexity.

A central problem in liver physiology is the identification of central veins (CV) and portal veins (PV) to characterize liver zonation^44^. This problem can be solved by considering expression patterns of marker genes, cell type localization and protein abundance. CVs can be identified by the central vein-associated endothelial cell marker *Axin2*^45^ (Fig. 3d). Similarly, *Vwf,* a known marker for endothelial cells in blood vessels, indicates the presence of both CVs and PVs^46^. Yet, due to the limited number of measured genes in the spatial transcriptomics data, it proved challenging to identify PV based on marker gene expression. Leveraging moscot, we alleviated this constraint by mapping the expression of the PV-specific markers *Adgrg6* and *Gja5*^42^ (Fig. 3e and Supplementary Fig. 7). Another limitation to characterize cellular niches of liver zonation was the lack of detailed cell type annotation and protein expression. Hence, we transferred the cell type annotation provided by the single-cell dataset using moscot (Fig. 3f). Focusing on resident liver macrophages called Kupffer cells, we could confirm their enriched presence in areas around CV where liver sinusoids are more prevalent^42^. We corroborated our findings by mapping the *folate receptor beta* protein to its spatial organization using moscot (Fig. 3f)^42^. By integrating results from cell type annotation, measured and imputed marker genes as well as transferred protein expression, we could characterize in-depth the tissue niche of liver zonation in a mouse liver sample.

In addition to mapping cells to spatial coordinates, analysts often face the challenge of building a consensus view of the tissue of interest, which requires aligning several spatial measurements from contiguous sections or from the same section from different biological replicates. With the *Alignment* problem, moscot allows to both align several sections as well as build such a consensus view from multiple spatial transcriptomics slides (Fig. 3g). This is an important step towards building a common coordinate framework (CCF)^47^ of biological systems. First, we evaluated moscot’s capability to spatially align synthetic datasets in a benchmark study adapted from Jones et al.^48^ and Zeira et al.^13^. The benchmark results showed that moscot performs on par or better than the previously published method PASTE^13^ (Supplementary Fig. 8 and Methods).

Having demonstrated moscot’s capabilities in a benchmark against state-of-the-art models, we set out to investigate the methods’s scalability to larger datasets. As an example, we considered the Vizgen MERSCOPE brain coronal sections (Methods)^43^. Such a dataset is prohibitively large for competing methods like PASTE, as only moscot is able to leverage low-rank factorization of the coupling matrix (∼250k cells for coronal brain section 1 and ∼300k cells for coronal brain section 2; Methods). Moscot accurately aligned two samples to the reference slide (Fig. 3h-i and Supplementary Fig. 9) for both coronal sections of the mouse brain. We evaluated the alignment by investigating gene correspondence of aligned sections. Because of the lack of cell type annotation, we set out to evaluate the alignment by comparing gene densities in cellular neighborhoods across sections (Methods). We observed that for most genes, there is a strong correspondence of gene expression densities across cellular neighborhoods both quantitatively (Supplementary Fig. 10) and visually (Supplementary Fig. 11). By visualizing one particular gene (*Slc17a7*) we could observe its expected expression in regions such as cortical layers and hippocampus (Fig. 3h-i). We showed that moscot enables the application of OT to large-scale spatial data analysis problems. Moscot allowed us to align large spatial transcriptomics datasets and enhance them trough imputation of gene expression, protein expression and cell type annotations.

### Charting the spatiotemporal dynamics of mouse embryogenesis with moscot

The advent of spatially-resolved single-cell datasets of developmental systems enables the characterization of cellular differentiation in space and time. Such data presents the challenge of developing methods that are able to delineate cellular trajectories, leveraging both intrinsic and extrinsic effects on cellular phenotypes. To this end, we introduce a novel trajectory inference method that incorporates similarities at gene expression and physical distances to infer a more accurate matching between cellular states within spatiotemporal datasets. It consists of a FGW-type problem that merges the algorithmic capabilities of moscot.time and moscot.space in a novel spatiotemporal method (Methods).

We assessed moscot’s capabilities to perform trajectory inference in the mouse embryogenesis spatiotemporal transcriptomic atlas (MOSTA)^3^, consisting of eight time points, from E9.5 to E16.5. Using spatially enhanced resolution omics-sequencing (Stereo-seq), this dataset captures the developing mouse at full embryo scale with spatial transcriptomics. Here, we analyzed a single slide for each time point with a total of ∼500,000 cells (Fig. 4a and Methods). To assess the performance of our new method, we evaluated the computed trajectories using the cell type transition score (similar to the analysis in moscot.time; Supplementary Table 5 and Methods). We compared the results with a W-type problem that incorporates only gene expression information (Fig. 2). Accounting for spatial similarity in the trajectory inference resulted in an improved prediction of cell transitions scores (Fig. 4b and Methods). We also found that the performance is robust with respect to hyperparameters (Supplementary Fig. 12).

**Fig. 4.**
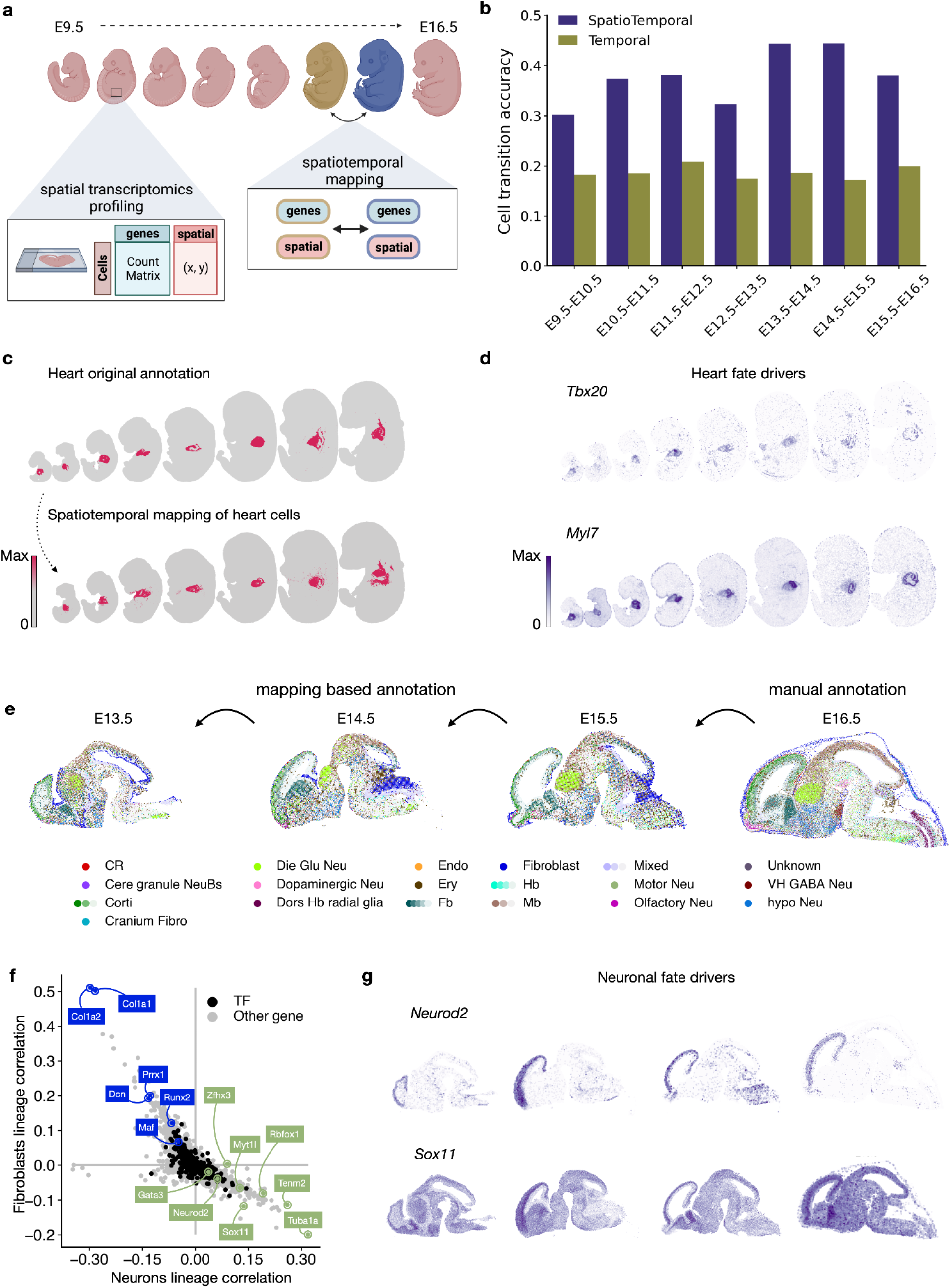
Inference of spatiotemporal dynamics with moscot. **a.** Schematic of applying moscot to spatiotemporal trajectory inference. Moscot allows for the integrative analysis of a time-resolved spatial transcriptomics dataset of mouse embryogenesis. **b.** Accuracy on curated transitions across developmental stages (Methods and Supplementary Table 5) with and without incorporation of spatial information**. c.** Mapping heart cells across time points (bottom) and the ground truth annotation of the heart lineage (top). **d.** Heart lineage driver genes found by interfacing moscot with CellRank^35^. (top) *Tbx20*^50^, a transcription factor known to play a variety of fundamental roles in cardiovascular development and (bottom) *Myl7*^53^ a gene related to metabolism and heart regeneration. **e.** Transferring high-resolution cell type annotations only provided in the latest time point (E16.5) to earlier time points. **f.** Pearson correlations of gene expression with Neuronal (x-axis) and Fibroblast (y-axis) fate probabilities. Annotated genes are among the 20 top driver genes and were previously associated with Fibroblasts and Neuronal lineage (Supplementary Tables 7-8). **g.** Spatial visualization of sample neuronal driver genes, *Neurod2*^57^ (top), and *Sox11*^55, 56^ (bottom).

Having established moscot’s performance in recapitulating expected cell transitions, we focused our analysis on the cell fates of the heart and the brain of the developing mouse embryo. For each pair of consecutive time points, we visualized heart cells at the earlier time point and where these cells are mapped to in the later time point (Fig. 4c). Noticeably, the predicted location of heart cells matches the annotation very well. To further characterize the cellular dynamics, we interfaced moscot with CellRank^35^, a state-of-the-art method for cell fate mapping (Methods). The convenient interface enables the identification of cell fates by CellRank, leveraging the spatiotemporal couplings learnt by moscot. Importantly, cell fates identified by CellRank correspond to the known differentiation lineages of the mouse embryo^3^ (Supplementary Fig. 13). We identified known driver genes of heart development, such as the transcription factors *Gata4*^49^ and *Tbx20*^50^ and genes related to metabolism and heart regeneration, such as *Myl7* and *Myh6* (Fig. 4d and Supplementary Table 6)^51–53^.

Understanding the evolution of cell types across time is crucial when studying developmental trajectories. Chen et al.^3^ provided a cell type annotation of the brain tissue at E16.5, but not for earlier time points. To investigate developmental trajectories in the brain, we utilized moscot to transfer cell type annotation from the E16.5 data to the three preceding time points, down to E13.5. Qualitatively, the predicted annotations retained the spatial distribution of the manual annotation (Fig. 4e), and quantitatively, they showed strong correspondence with reported marker genes (Supplementary Fig. 14 and Methods).

At last, the interplay between moscot and CellRank allowed us to identify terminal cell states of brain development in the mouse embryo, with fate probabilities which are in accordance with the predicted annotation (Supplementary Fig. 15). As done for the heart, we used CellRank to predict neuron and fibroblast driver genes (Fig. 4f-g, and Methods). For the neuronal fate, the top identified transcription factors (TFs) have been previously reported as relevant for neuronal development, indicating the reliability of the analysis (Supplementary Table 7). Among the TFs are *Tcf7l2*^54^, *Sox11*^55, 56^, *Myt1l*^55, 56^, and *Zfhx3*^57^. We further obtained key genes which were previously reported such as *Tuba1a*^58^, *Tenm2*^59^ and *Rbfox1*^60^. Importantly, our results included known spatially localized drivers, such as *Neurod2*, associated with forebrain glutamatergic neurons^61^, as well as non-regional drivers, such as *Sox11* (Fig. 4g). For fibroblasts, we identified the transcription factors *Prrx1*, *Runx2*, and *Msx1*^56, 62, 63^, and known key genes such as *Dcn*, *Col1a2* and *Col1a1*^64^ (Supplementary Table 8).

In summary, we demonstrated that moscot provides a novel analysis method to investigate the spatiotemporal dynamics of cellular differentiation. By combining moscot with CellRank, we characterized the spatiotemporal dynamics of mouse embryogenesis, recovering known markers of differentiation of the developing heart and brain.

### Disentangling lineage relationships of delta and epsilon cells in pancreatic development leveraging multiple modalities with moscot

To highlight moscot’s potential for disentangling complex lineage relationships, we focused on the poorly understood process of delta and epsilon cell formation during mouse pancreatic development^35, 65, 66^ (Supplementary Note 4). Hypotheses of lineage specification range from delta cells splitting simultaneously with alpha and beta cells after going through a Fev+ cell state^67^ to delta cells being derived from the same progenitor population as beta cells^68^. In previous work, we had hypothesized delta cells to evolve from a *Fev*+ population^35, 65^, but we could not resolve their precise lineage hierarchy. Similarly, while our previous analysis suggested that epsilon cells can be generated from alpha and *Ngn*3+ cells, Yu et al.^68, 69^ found that epsilon cells derive exclusively from *Ngn3*+ cells, and can in turn give rise to alpha cells. Previous studies demonstrated that combining the measurements of gene expression and open chromatin regions allows for a more holistic view of developmental cellular systems^70–72^. Thus, we generated a single nucleus RNA and assay for transposase-accessible chromatin (ATAC) multi-omics dataset of E 14.5 (9,811 nuclei) and 15.5 (7,107 nuclei) of the pancreatic epithelium enriched for endocrine progenitors (EPs) using the Ngn3-Venus fusion (NVF) mouse line^65^ (Fig. 5a and Methods). *Ngn3* is a TF expressed in endocrine precursors. Hence, enrichment of *Ngn3*+ cells allows for a detailed study of endocrine lineage formation. Compared to previous scRNA/ATAC-seq studies that relied on bulk ATAC measurements^66^ or a low number of cells for scRNA-seq^67^, our novel dataset allows for a detailed and comprehensive multimodal analysis of endocrine cell differentiation.

**Fig. 5.**
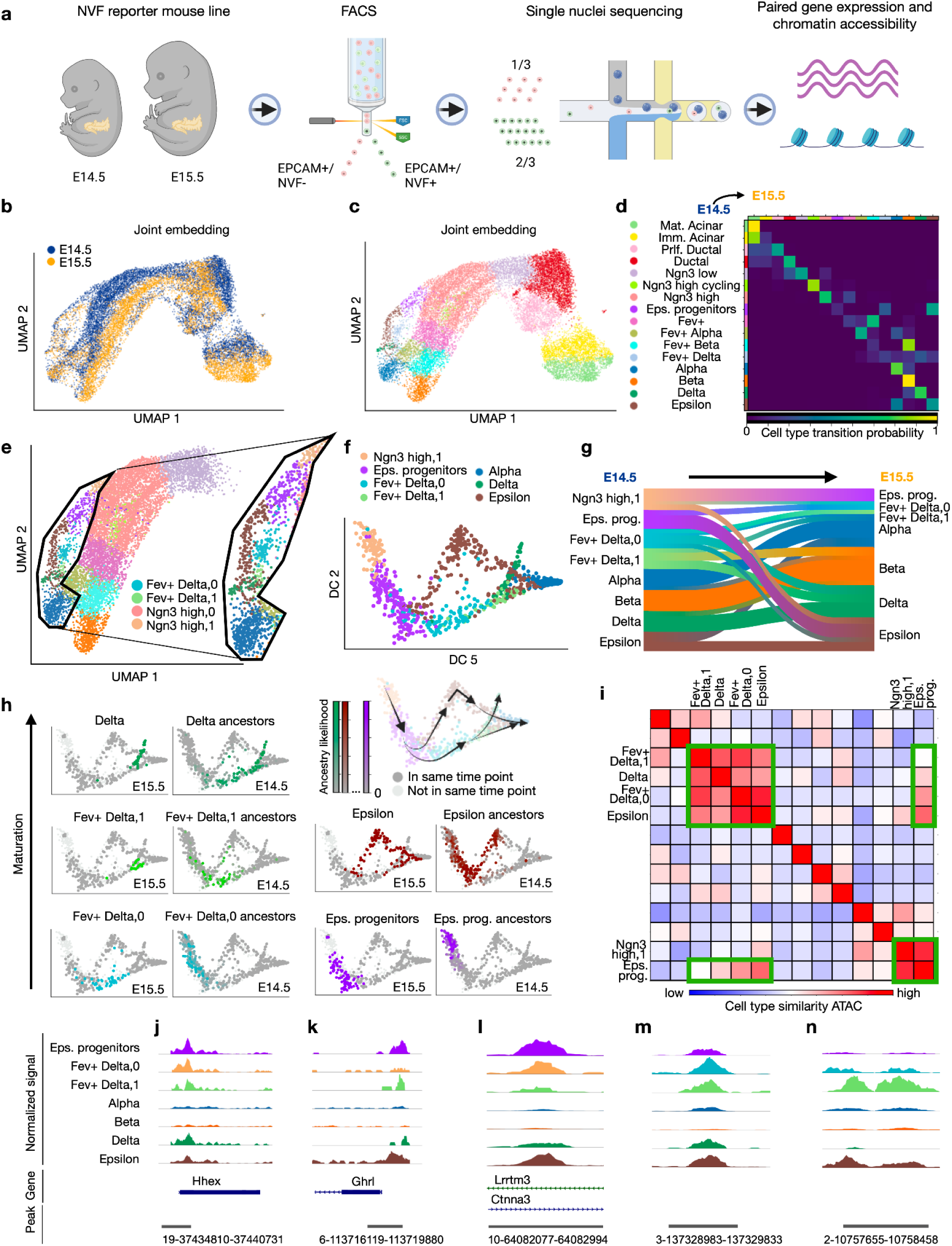
moscot disentangles lineage ancestries of delta and epsilon cells. **a.** Schematic of the experimental protocol to generate paired gene expression and ATAC data capturing the development of the mouse pancreas. **b.-c.** Multimodal UMAP^96, 97^ embedding, colored by time (b) and cell type annotation (c) (Methods). **d.** Heatmap visualizing descendancy probabilities of cell types in E14.5 as obtained by moscot.time**. e.** UMAP embedding, restricted to endocrine cells and their progenitors, colored as in (c), highlighting sub-clustered Ngn3^high^ and Fev+ delta populations. The inset highlights alpha cells as well as delta and epsilon cells, and their putative progenitors. **f.** Multimodal diffusion map of the cells which are inset in panel (e). **g.** Sankey diagram of the E14.5-15.5 cell type transitions. **h.** Putative developmental trajectory per cell type in E15.5 and its corresponding ancestor population in E14.5. **i.** Similarity in ATAC profile between different cell types (Methods). The green boxes highlight the cell types whose ancestry we focus on. **j.-k.** Chromatin accessibility around the promoter region of *Hhex* (j), a key transcription factor for delta cells^80^, and *Ghrl* (k), the gene corresponding to the hormone released by epsilon cells. **l-n.** Chromatin accessibility of the most accessible peak in the epsilon progenitors population (l), the Fev+ delta-0 population (m), and the Fev+ delta-1 population (n) (Methods).

As expected, we observed a distribution shift between the two time points (Fig. 5b and Supplementary Fig. 16). Clustering based on both modalities revealed the expected cell type heterogeneity in the endocrine branch, ranging from Ngn3^low^ to heterogeneous progenitors of terminal endocrine cell states (Fig. 5c, Supplementary Fig. 17, Supplementary Table 9 and Methods). We linked the cells of the two time points using moscot.time by leveraging information from both gene expression and ATAC (Supplementary Note 5). To validate the coupling, we aggregated the coupling matrix to the cell type level and found the majority of recovered transitions to be supported by literature^35, 65, 67, 73–75^ (Fig. 5d, Supplementary Fig. 18 and Methods).

Subsequently, we explored the lineage segregation of delta and epsilon cells with moscot.time. Therefore, we restricted our analysis to the endocrine branch and subclustered the Ngn3^high^ EP and Fev+ delta populations (Fig. 5e). To emphasize the developmental axes of variation, we computed a diffusion map (Fig. 5f). We used moscot.time to compute putative descendants of cells in E14.5 and found that mature endocrine cells mostly evolve into themselves, as expected (Fig. 5g and Methods). Moreover, epsilon cells partly evolve into alpha cells (Fig. 5e and Supplementary Fig. 19), which has been reported in recent literature^68, 69^. We found the epsilon population to be mainly derived from epsilon progenitors, which themselves originate from the Ngn3^high^-1 subcluster (Fig. 5g-h and Supplementary Fig. 20). Contrary to our recent hypothesis that precursors of all endocrine cells express *Fev* during development^65^, our model implies that epsilon cells mature without necessarily going through a state of high expression of *Fev* (Supplementary Fig. 21). Yet, we found a smaller proportion of the epsilon cells to be derived from the Fev+ delta-0 and Fev+ delta-1 populations.

Next, we investigated whether the Fev+ delta-0 and Fev+ delta-1 subclusters could give rise to a population other than epsilon cells. We hypothesize Fev+ delta-0 cells to be direct ancestors of Fev+ delta-1 cells. In fact, we found Fev+ delta-1 cells to subsequently give rise to delta cells, in line with our previous findings^35, 65^ (Fig. 5g-h and Supplementary Fig. 22, Supplementary Fig. 23). At the same time, moscot.time predicted cells in Fev+ delta-1 to be able to evolve into alpha and, to a larger extent, beta cells. Thus, we found that the Fev+ delta population retains high plasticity, carrying the potential of developing into either of the four endocrine cell states alpha, beta, delta, and epsilon.

To fully delineate the lineage relationships of delta and epsilon cells, we focused on the ancestry of Fev+ delta cells. Based on the results of moscot.time, we conjectured Fev+ delta cells to also be derived from epsilon progenitors (Fig. 5h and Supplementary Fig. 24). Hence, our analysis indicated a common lineage origin of epsilon and delta cells, as both are predicted to evolve from epsilon progenitors. Having delineated the cell trajectories of endocrine cells and their progenitors, we used moscot.time to find marker genes (Supplementary Fig. 25 and Methods). The recovery of known marker genes like *Arx* and *Mafb*^76^ of the well-studied alpha and beta cells, respectively, validated this method (Supplementary Fig. 26). For example, we find *Nefm,* which is known to play a role in endocrine maturation^77, 78^, to be involved in the fate specification of epsilon progenitors. Similarly, *Cdkn1a,* a known cell cycle inhibitor^79^, turned out to be among the top four marker genes for the Fev+ delta population.

To corroborate the hypothesis of a similar ancestry between epsilon and delta cells, we investigated the similarity of chromatin regions for cell types in the endocrine branch. The similarity between the ATAC profiles of Ngn3^high^-1, epsilon progenitors, Fev+ delta-0, Fev+delta-1, epsilon, and delta is striking (Fig. 5i and Supplementary Fig. 27). We continued the analysis of co-accessibility by considering specific chromatin regions. We observed noticeable similarities in chromatin accessibility in the promoter regions of both *Ghrl* (epsilon) and *Hhex,* the key TF of delta cells^80^ (Fig. 5j-k and Supplementary Fig. 28). To obtain further relevant chromatin regions, we performed differentially accessible peak analysis of the epsilon progenitor (Fig. 5l), Fev+ Delta-0 (Fig. 5m), and Fev+ Delta-1 (Fig. 5n) populations and found that the peaks are co-accessible among the hypothesized ancestors of delta and epsilon cells (Supplementary Fig. 29 and Supplementary Note 6).

Another way to understand regulatory mechanisms at single-cell resolution is motif analysis. Similarity in motif profiles is an indication of a similar cell state as related TFs govern the developmental trajectories. Due to a temporal shift between gene expression of a TF and the activity of a TF, profiling of motif activity and gene expression within the same sample might fail to recover regulatory mechanisms^81^. Moscot allows alleviating this constraint by linking gene expression in an earlier time point with motif activity in cells corresponding to the later time point (Supplementary Fig. 30, Supplementary Fig. 31, Methods and Supplementary Note 7). We found *Isl1* to have a high motif activity in delta cells, complemented by a high gene expression in their progenitors. Similarly, we report a motif associated with *Tead1* to be highly active in epsilon cells, and the gene to be highly expressed in their progenitors. The similarity in motif activity among predicted ancestors of delta and epsilon cells corroborates the hypothesis of shared ancestry of delta and epsilon cells.

Thus, we showed how moscot can help to delineate complex lineage relationships. Multimodality helped to both recover meaningful trajectories and bridge the time gap between the expression of a transcription factor and its activity.

## Discussion

We presented moscot, a computational framework for mapping cellular states across time and space using optimal transport (OT). Unlike previous applications of OT, moscot incorporates multimodal information, scales to atlas-sized datasets, and provides an intuitive and consistent interface. We demonstrated the added value of our scalable implementation across various biological problems. In particular, we accurately recovered murine differentiation trajectories during embryogenesis across time and space^3, 16^, enriched spatial liver samples with multimodal information^42^, and aligned brain tissue slides^43^ in datasets that were previously inaccessible with state-of-the-art techniques. We also showcased the benefits of multimodal information in a new developmental pancreas dataset where we learned differentiation trajectories in a joint space of epigenetic and transcriptomic information. Notably, our multimodal trajectory reconstruction predicted a shared lineage origin of epsilon and delta cells in the pancreas, which is supported by computational analysis based on similar epigenetic regulation. While this hypothesis requires further experimental validation, our results demonstrate the potential of moscot to generate novel insights into complex biological systems.

Cellular differentiation is a tightly regulated process that is influenced by a range of internal and external factors. In particular, the spatial organization of tissues plays a critical role in guiding differentiation, and understanding this process requires temporal sampling of molecular states within their spatial context. Although spatiotemporal data is beginning to emerge^3, 82, 83^, current computational methods for analyzing such data are limited. With moscot, we present one of the first analysis approaches for spatiotemporal data. Combined with CellRank^35^, our framework predicted spatiotemporal differentiation trajectories in mouse embryogenesis and identified their putative regulators. Parallel to moscot, Qiu et al. developed SPATEO^84^ which also uses OT to map cells across spatial time courses. However, SPATEO is less scalable: on the same MOSTA Stereo-Seq application^3^, SPATEO required downsampling to 2,000 cells per embryo^84^ while moscot mapped entire embryos (500,000 cells). Further work and community contributions can focus on incorporating the effect of cell-cell communication on observed differentiation trajectories. Such an extension would enable studying the complex interplay between intrinsic and extrinsic effects on cellular differentiation in even greater detail.

Moscot will greatly simplify future OT applications by the single-cell analysis community in Python. With our unified API and extensive documentation, incorporating cross-modality data integration^17, 85^, patient manifold learning^86, 87^, and cell-cell communication inference^88, 89^ becomes substantially easier. Importantly, new applications of OT to problems in single-cell biology will benefit from the implementation advancements presented in this work, including GPU acceleration, linear time and memory complexity through low-rank approximations, and multimodal support. While low-rank approximations allow applications to atlas-scale datasets, they have not yet been combined with unbalancedness, where marginal constraints do not have to be satisfied. This concept is important when cell type proportions vary substantially across time points or spatial slides, and can be incorporated by future methodological advances.

Moscot is a powerful approach to map observed cell state distributions across biological representations. The current approach of using discrete OT is well-suited for the applications described in this paper and for the extensions outlined above. However, these models are not applicable to out-of-sample data points. Recently, Neural OT^90^ has been suggested to alleviate this limitation. Neural OT has been successfully applied to study perturbation effects^91–93^ and developmental trajectories at single-cell level^92, 94^. In the latter case, the use of Neural OT is particularly compelling for building developmental trajectories of cellular atlases as computed OT maps can be applied to new datasets without refitting the model.

The ability to map and align single-cell genomic data is becoming increasingly important as datasets grow larger and more complex, with new modalities being measured at varying temporal and spatial resolutions^95^. Moscot allows scientists to extract biological insights across time- and spatially resolved complex datasets, and is a powerful tool for investigating cellular differentiation, spatial organization, and other biological processes.

## Acknowledgments

The following figures have been created with BioRender: Fig. 1, Fig. 2a, Fig. 3a,g Fig. 4a, Fig. 5a. We thank Meshal Ansari for the design of the moscot logo. We are grateful for the work of the Genomics Core Facility of Helmholtz Munich who performed the sequencing of the pancreas data. Moreover, we would like to thank Ignacio Ibarra for valuable input on motif databases and helpful comments on the manuscript. We thank Sara Jimenez for improvements on the manuscript, Meyer Scetbon for insights into the low-rank optimal transport approximations, Alex Tong for advice on geodesic cost functions and Arina Danilina for useful comments on early implementations of the moscot alignment problem. This work was supported by the BMBF-funded de.NBI Cloud within the German Network for Bioinformatics Infrastructure (de.NBI) (031A532B, 031A533A, 031A533B, 031A534A, 031A535A, 031A537A, 031A537B, 031A537C, 031A537D, 031A538A). G.P. is supported by the Helmholtz Association under the joint research school Munich School for Data Science and by the Joachim Herz Foundation. M.L. acknowledges financial support by the Joachim Herz Foundation. Z.P. is supported by a scholarship for outstanding doctoral students in data-science by the Israeli Council for Higher Education and the Clore Scholarship for Ph.D students. M.N. is supported by an Azrieli Foundation Early Career Faculty Fellowship, the Center for Interdisciplinary Data Science Research at the Hebrew University of Jerusalem, the Israel Science Foundation (Grant no. 1079/21), and the European Union (ERC, DecodeSC, 101040660). F.J.T. acknowledges support by the Helmholtz Association’s Initiative and Networking Fund through Helmholtz AI (grant # ZT-I-PF-5-01) and by Wellcome Leap as part of the ΔTissue Program. For all support coming via EU funding, the views and opinions expressed are those of the author(s) only and do not necessarily reflect those of the European Union or the European Research Council. Neither the European Union nor the granting authority can be held responsible for them.

## Author contributions

D.K., G.P., M.L., M.K., and Z.P. contributed equally. M.L. conceived the initial idea of the project, guided by F.J.T. D.K., G.P., M.L., M.K., and Z.P. conceptualized the project. M.K., D.K., and G.P. designed and implemented the moscot software package with contributions from Z.P and L.M.P. M.G. benchmarked and conducted analyses for moscot.time together with M.L, with contributions from D.K. G.P. benchmarked and conducted analyses for moscot.space with contributions from D.K.. Z.P. conducted analyses for moscot.spatiotemporal with contributions from G.P. and D.K. A.B.P. and M.T.M. generated the pancreas data under supervision of H.L.. D.K. processed and analyzed the pancreas dataset with contributions from M.S., M.B., and M.L.. M.L., D.K., and G.P. wrote the manuscript with contributions from all authors. M.L. and

D.K. wrote the Supplementary Notes. F.J.T., M.C., and M.N. supervised the project and contributed to the conception of the project. All authors read and approved the final manuscript.

## Data availability

The mouse embryogenesis atlas by Qiu et al.^16^ is available at http://tome.gs.washington.edu. The mouse liver CITE-seq by Guilliams et al. ^42^ is available at https://www.livercellatlas.org/. The Vizgen MERSCOPE Liver and Brain coronal sections dataset is available at the Vizgen public dataset release website https://vizgen.com/data-release-program/. The datasets for benchmarking the spatial mapping problems were taken from the original publication of Li et al.^41^. The spatiotemporal atlas of mouse embryogenesis (MOSTA) by Chen et al.^3^ is available at https://db.cngb.org/stomics/mosta/. The pancreas data will be made available soon.

## Code availability

The moscot software package is available at https://moscot-tools.org including documentation, tutorials and examples. Code to reproduce our analysis is available at https://github.com/theislab/moscot-framework_reproducibility and code to reproduce the benchmarks can be found at https://github.com/theislab/moscot_benchmarks.

## Ethics statement

Animal studies were conducted with adherence to relevant ethical guidelines for the use of animals in research in agreement with German animal welfare legislation with the approved guidelines of the Society of Laboratory Animals (GV-SOLAS) and the Federation of Laboratory Animal Science Associations (FELASA). The study was approved by the Helmholtz Munich Animal Welfare Body and by the Government of Upper Bavaria. Ngn3-Venus fusion (NVF) mice were kept at the central facilities at Helmholtz Munich under SPF conditions in animal rooms with light cycles of 12/12 h, temperature of 20–24°C, and humidity of 45–65%. The mice received sterile filtered water and a standard diet for rodents ad libitum.

## Supplementary Materials

Supplementary Figures: Supplementary Figures 1-31

Supplementary Tables: Supplementary Tables 1-9

Supplementary Notes: Supplementary Notes 1-7

## Moscot methods section

## 1 The moscot algorithm

### 1.1 Model overview and introduction

`moscot` (**m**ulti-**o**mic **s**ingle **c**ell **o**ptimal **t**ransport) is a software framework for scalable, multi-modal applications of optimal transport (OT) to single-cell genomics with consistent user experience. With `moscot`, we couple cells across experimental time points (`moscot.time`), map dissociated single-cell datasets into space (`moscot.space.mapping`), align tissue slides (`moscot.space.alignment`), and reconstruct spatiotemporal trajectories (`moscot.spatiotemporal`). Thus, moscot represents a unified and extensible framework to map cells across time and space.

#### 1.1.1 Optimal Transport (OT) for single-cell genomics

OT is an area of mathematics that is concerned with comparing probability distributions in a geometry-aware manner^1^. OT-based tools have been successfully applied to various problems that arise in single-cell genomics, including mapping cells across time points^2–7^, mapping cells from molecular to physical space^8, 9^, aligning spatial transcriptomics samples ^10^, integrating data across molecular modalities^11, 12^, learning patient manifolds^13, 14^ or mapping cells across different experimental perturbations ^15, 16^. Despite this notable success, the widespread adaptation of OT-based tools in single-cell genomics faces three key challenges.

First, the majority of current OT-based tools is geared towards a single modality and cannot use the added information provided by multi-modal assays. Second, compute time and memory consumption scale quadratically in cell number for vanilla OT and cubically for Gromov-Wasserstein extensions ^8–10, 17^. Such poor scalability limits the application of these tools to current datasets containing millions of cells. Third, the landscape of OT-based tools is split across programming languages and software providing OT algorithms, resulting in a fractured landscape of incompatible application programming interfaces (APIs). This makes it difficult for users to adapt and for developers to create new tools. In contrast, user-friendly and extensible APIs accelerate and facilitate research, as powerfully demonstrated through the scVI-tools framework ^18^.

### 1.1.2 moscot unlocks the full power of OT for spatiotemporal applications

Our method is built on three key design principles to overcome previous limitations and unlock the full potential of OT for single-cell applications:

1. Multi-modality: all moscot models extend to multi-modal data, including CITE-seq^19^ and multiome ^20–22^ (RNA and chromatin accessibility) data.
2. Scalability: we use both engineering and methodological innovations to overcome previous scalability limitations; in particular, we reduce compute time and memory consumption to be linear in the number of cells.
3. Consistency: our implementation unifies temporal, spatial, and spatiotemporal problems through one consistent API that interacts with the wider scverse/SCANPY ^23, 24^ ecosystem and is easy to use. Solving any of these problems in moscot follows a common pattern that translates the biological problem into an OT problem that is solved by the scalable optimal transport tools (OTT)^25^ backend.

We describe in the following sections how we realize these principles for temporal, spatial, and spatiotemporal applications. Our modular interface easily extends to new biological problems; we anticipate and encourage community contributions to solve a growing number of single-cell mapping and integration problems using `moscot`. Our framework is implemented as user-friendly, open-source software with extensive documentation, examples, and tutorials, available at https://moscot-tools.org.

### 1.2 moscot.time for mapping cells across time

#### 1.2.1 Model rationale, inputs, and outputs

Biologists frequently employ time series experiments to study biological processes like development or regeneration that are not in a steady state. As current single-cell assays are cell-destructive, such experiments result in disparate molecular profiles measured at different time points. As previously suggested ^2^, OT can be used to probabilistically link cells from early to late time points. Previous methods had limited scalability and were only applied to gene expression; we outline in the following how `moscot.time` overcomes these limitations.

Let thus *X* ∈ ℝ^N×D^ and *Y* ∈ ℝ^M×D^ represent pairs of state matrices for *N* and *M* cells observed at early (*t*_1_) and late (*t*_2_) time points, respectively. State matrices *X* and *Y* may represent e.g. gene expression (scRNA-seq) or chromatin accessibility (scATAC-seq ^26^) across *D* features (e.g. genes or peaks). Optionally, the user may provide marginal distributions ***a*** ∈ *Δ_N_* and ***b*** ∈ *Δ_M_* over cells at *t*_1_ and *t*_2_ for probability simplex 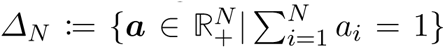. Any cell-level prior information may be represented through the marginals, including cellular growth- and death rates.

`moscot.time’s` key output is a coupling matrix *P* ∈ *U* (***a***, ***b***) where *U* (***a***, ***b***) is the set of feasible coupling matrices, defined by

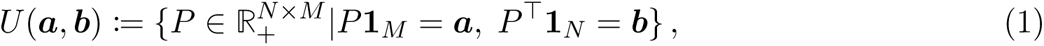

for constant one vector **1***_N_* = [1*, …,* 1]*^⊤^* ∈ ℝ*^N^* . We link *t*_1_-cells to *t*_2_-cells through the coupling matrix *P* ; the *i*-th row *P_i,_*: represents the amount of probability mass transported from cell *i* at *t*_1_ towards any *t*_2_-cell. The set *U* (***a***, ***b***) contains those coupling matrices *P* that are compatible with the user-provided marginal distributions ***a*** and ***b*** at *t*_1_ and *t*_2_, respectively.

These definitions allow us to formalize the aim of `moscot.time`: we seek to find a coupling matrix *P* ∈ *U* (***a***, ***b***) which couples *t*_1_-cells to *t*_2_-cells such that their overall traveled distance in phenotypic space is minimized.

#### 1.2.2 Model description

To quantify the distance cells travel in phenotypic space between time points, let *c*(***x****_i_,* ***y****_j_*) be a cost function for early (***x****_i_*) and late (***y****_j_*) molecular profiles, representing e.g. gene expression or chromatin accessibility state. moscot allows for the use of various cost functions (Supplementary Note 5). We use the cost function *c* to measure cellular distances in a modality-specific, shared latent space, e.g. PCA for gene expression data, latent semantic indexing^27^ (LSI) for ATAC data or corresponding scVI-tools models ^18^.

We evaluate the cost function *c* for all pairs of cells (*i, j*) ∈ {1*, …, N* } × {1*, …, M* } to form the cost matrix 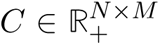 . Given the cost matrix *C* quantifying distances along the phenotypic manifold, we solve the optimization problem

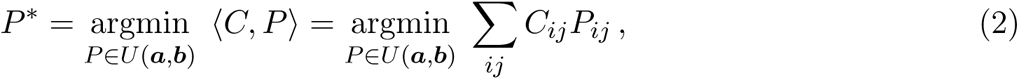

known as the Kantorovich relaxation of OT ^1^ where *P^∗^* is the optimal coupling matrix. When using *P^∗^* to transport *t*_1_-cells to *t*_2_-cells, we accumulate the lowest cost according to *C*. In the following, we refer to this type of OT problem as a *Wasserstein* (W)-type OT problem.

##### Introducing entropic reguluarization

In practice, the OT problem of Equation (2) is usually not solved directly because it is computationally expensive, and the solution has statistically unfavorable properties ^1, 28^. Instead, it is much more common to consider a regularized version of the problem,

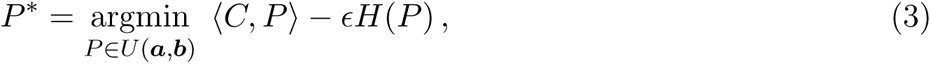

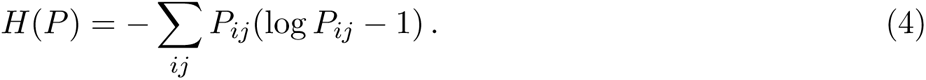

The parameter *ɛ >* 0 controls the regularization strength. Intuitively, entropic regularization introduces uncertainty to the solution; it has a “blurring” effect on *P^∗^*. Mathematically, it renders the problem *ɛ*-strongly convex, differentiable and less prone to the curse of dimensionality ^1, 28, 29^.

##### The Sinkhorn algorithm for optimization

It can be shown that the solution to the regularized W-problem of Equation (3) has the form *P_ij_* = *u_i_K_ij_v_j_* for Gibbs kernel

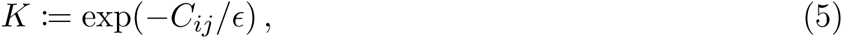

and unknown scaling variables 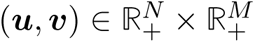. Using this formulation, we may rewrite the constraints *P* **1***_M_* = ***a*** and *P* **1***_N_* = ***b*** of Equation (1) to yield

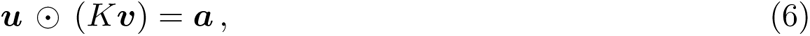

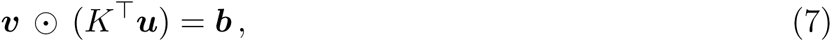

where ⊙ denotes element-wise multiplication. Iteratively solving these equations gives rise to *Sinkhorn’s algorithm* ^29–31^,

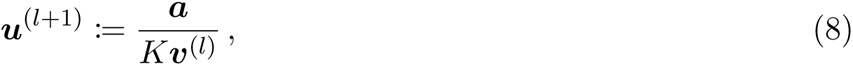

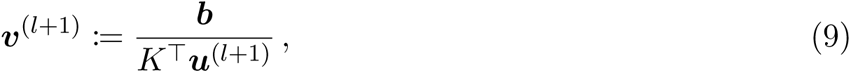

where the division is applied element-wise, and *l* is the iteration counter. Using this algorithm, the (unique) solution to the regularized W-problem of Equation (3), corresponding to the optimal coupling of *t*_1_-cells to *t*_2_-cells, can be computed in time and memory quadratic in cell number ^1^.

##### Adjusting the marginals for growth and death

Cells differentiate, proliferate, and die as the biological process unfolds between time points *t*_1_ and *t*_2_. The coupling matrix *P^∗^*, computed by solving Equation (3), reflects a mixture of these effects. To disentangle proliferation and apoptosis from differentiation, we adjust the left marginal ***a*** for cellular growth and death. Specifically we follow Schiebinger et al. ^2^ in defining

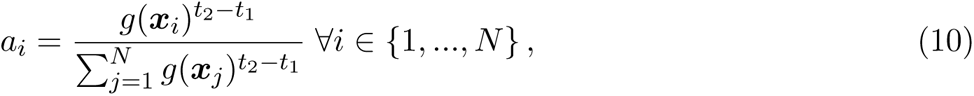

where *g* : ℝ*^D^* → ℝ_+_ corresponds to the expected value of a birth-death process *g*(***x***) = *e^β^*^(^***^x^***^)^*^−δ^*^(^***^x^***^)^ with proliferation at rate *β*(***x***) and death at rate *δ*(***x***). We estimate growth- and death rates from curated marker gene sets; moscot comes with pre-defined gene sets for mice and humans. For the right marginal ***b***, we assign uniform weights *b_j_* = 1*/M*, ∀*j* ∈ {1*, …, M* }. Intuitively, our adjustment allows *t*_1_-cells cells likely to proliferate (die) to distribute more (less) probability mass towards *t*_2_-cells. Such an adjustment encourages the optimal coupling matrix *P^∗^* to reflect differentiation rather than proliferation and apoptosis.

As it is difficult to adjust the hyperparameters of the proliferation and apoptosis rate, we also implement a more intuitive and more easily adjustable estimation of the growth rates via

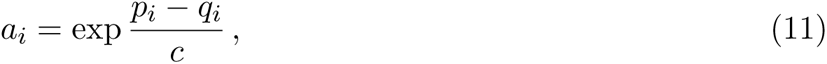

where *p_i_* denotes a proliferation score and *q_i_* an apoptosis score, obtained by `scanpy.tl.score_genes`. *c* denotes a scaling parameter.

##### Unbalancedness for stochastic cell sampling and uncertain marginals

Our formulation of Equation (3) enforces the pre-specified marginals ***a*** and ***b*** to be exactly met by the solution *P^∗^*. This is problematic from two perspectives:

• The cells profiled at each time point usually correspond to a sample from the overall population - small variations in cell type frequencies across time points do not necessarily reflect underlying differentiation but might result from stochastic cell sampling. Exactly enforcing the marginals thus implies that we encode the sampling effect in the coupling, which confounds the actual differentiation signal.
• Our growth/death-adjusted marginals of Equation (10) are unlikely to reflect ground-truth proliferation/apoptosis rates as they are estimated based on noisy gene expression and do not include any post-transcriptional effects. Thus, exactly enforcing these marginals propagates noise into the coupling matrix *P^∗^*.

To avoid both pitfalls, we follow Schiebinger et al. ^2^ to allow small deviations from the exact marginals in an unbalanced OT framework^1, 32^. Specifically, we replace the hard constraint *P* ∈ *U* (***a***, ***b***) with soft KL-divergence penalties,

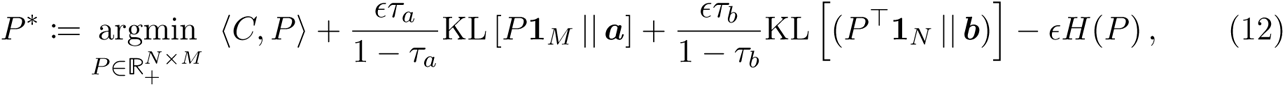

which may be solved at the same computational complexity level using a generalization of Sinkhorn’s algorithm ^1, 33, 34^. The parameters *τ_a_, τ_b_* ∈ (0, 1) are hyperparameters that determine the weight we give to complying with the left and right marginals ***a*** and ***b***, respectively. Values near one or zero correspond to strict or weak marginal penalties, respectively.

#### 1.2.3 Multi-modal data and scalability

The model we presented in the previous section is similar to the Waddington OT (WOT) model of Schiebinger et al. ^2^ . However, WOT was only applied to uni-modal data and has quadratic time- and memory complexity in the number of cells, largely preventing its application to atlas-scale temporal datasets containing multiple modalities. This section extends the `moscot.time` model to overcome both limitations.

##### Application to multi-modal data

We incorporate multi-modal data in `moscot.time` via an adjusted definition of the cost function. Intuitively, we use a joint representation to render the computed distances more faithful to the phenotypic manifold. Specifically, given bi-modal representations (*X*^(1)^*, X*^(2)^) and (*Y* ^(1)^*, Y* ^(2)^) at *t*_1_ and *t*_2_, respectively, we scale these to have the same variance, and measure distances in a concatenated space. In this example, (1) and (2) can represent any pair of modalities, for example, gene expression and ATAC data. This strategy naturally extends beyond two modalities towards any number of jointly measured modalities, making `moscot.time` truly *multi* -modal. Alternatively, `moscot.time` may be applied to representations computed using shared latent space learning techniques, including TotalVI^35^ for CITE-seq ^19^ data, and MultiVI^36^ or Multigrate ^37^ for shared ATAC and RNA ^20–22, 38^ readout.

##### Scalability through engineering-type innovations

To solve the W-problem of Equation (3), `moscot.time` builds on OTT^25^ in the backend which offers three key engineering-type improvements:

1. Online evaluation of the cost function.
2. GPU execution.
3. Just-in-time compilation (jitting).

While memory complexity is quadratic in vanilla Sinkhorn, it can be reduced to linear through online-cost matrix evaluation. The key observation is that the Sinkhorn algorithm only accesses the cost matrix *C* through the matrix-vector products *K****v*** and *K^⊤^****u*** (Equation (8)) which are evaluated row by row. Thus, the cost function *c* can be queried on the fly for those cell-cell distances which are required to evaluate the current row of the matrix-vector product. Online evaluation reduces the memory complexity to be linear in cell number ^25^ (first improvement).

Second, while the Sinkhorn algorithm can, in principle, be run on GPUs to greatly accelerate optimization^1^, the quadratic memory complexity prevents this in practice. While CPUs can handle very large memory consumption, GPUs are usually much more limited (typically around 40 GB). Online memory evaluation (first improvement) renders GPU acceleration possible, and OTT^25^ implements it in practice. Performing computations on GPU greatly accelerates the computation of cell-cell couplings in `moscot.time` (second improvement).

Third, jitting compiles python code before it is executed for the first time, further reducing compute time ^39^ (third improvement).

Combining these three engineering-type innovations allows `moscot.time` to run datasets containing a few hundred thousand cells per time point with linear memory and quadratic time complexity on modern GPUs. Once we go beyond that towards millions of cells per time point, the quadratic time complexity becomes prohibitive.

##### Scalability through methodological innovations

To enable the application of `moscot.time` to future datasets containing millions of cells per time point, we must overcome the quadratic time complexity in the number of cells. Following previous work ^40, 41^, we achieve this imposing low-rank constraints on the set of feasible couplings, i.e. requiring *P* ∈ *U* (***a***, ***b****, r*) for nonnegative coupling matrix rank *r* (Supplementary Note 3). Such a regularization leads to linear time and memory complexity in the number of cells. Low-rank Sinkhorn is implemented in OTT^25^ and available through `moscot.time`, enabling the application to future atlas-scale developmental studies.

#### 1.2.4 Downstream applications

The coupling matrix *P^∗^* optimally links *t*_1_-cells to *t*_2_-cells for the cost function *c*. `moscot.time` uses the coupling to relate cellular states and derive insights about putative driver genes; consider thus a *t*_1_ cell state *𝒫* of interest, where *𝒫* is the set of corresponding cell indices. This state may represent, for example, a rare or transient cell population. Define the corresponding normalized indicator vector ***p*** ∈ {0, 1}*^N^* via

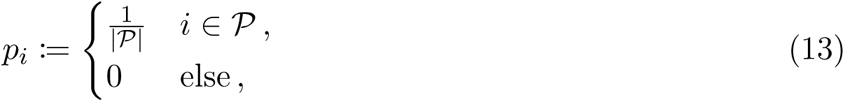

for *t*_1_-cell *i* and |𝒫|, the number of cells in state *𝒫*. Following the original suggestion by Schiebinger et al. ^2^, we compute *t*_2_-descendants of cell state *𝒫* by a push-forward operation of ***𝒫***,

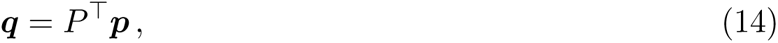

where 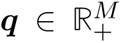 describes the probability mass that cell state *𝒫* distributes to *t*_2_-cells. Using *P* rather than its transpose, we analogously compute ancestors of a cell state *𝓠* at *t*_2_. For a global view of cell-state transitions, we aggregate pull- and push operations over all states into transition matrices which we visualize via heatmaps or Sankey diagrams. In addition, we correlate pull- and push distributions with gene expression to uncover putative driver genes.

##### Coupling more than two time points

Following the Waddington OT model ^2^, we couple several time points by assuming that the state of *t_r_*_+1_ cells depends only on the state of *t_r_* cells and not on any other earlier or later states. The index *r* runs over time points, *r* ∈ {1*, …, R*}, for *R* time points. This Markov assumption allows us to chain together time points by matrix multiplication; for time points {*t*_1_*, …, t_R_*} and corresponding sequential coupling matrices {*P* ^(1)^*, …, P* ^(*R*−1)^}, we link *t*_1_-cells to *t_R_* cells by matrix multiplication, *P* ^(1)^*P* ^(2)^*… P* ^(*R*−1)^.

### 1.3 moscot.space.mapping for spatial reference mapping

#### 1.3.1 Model rationale, inputs, and outputs

Techniques to simultaneously measure a cell’s spatial context and its transcriptional state have matured in recent years; in particular, spatial resolution, the field of view, and the number of profiled transcripts have increased ^42, 43^. However, current approaches still fall short of measuring the full transcriptome at true single-cell resolution. This experimental difficulty has fueled the development of a range of computational tools ^8, 44–48^ that map dissociated single-cell reference datasets onto spatial coordinates, a problem known as *spatial mapping*. Solving a spatial mapping problem can provide two types of information:

1. Annotation-centric perspective: spatial mapping annotates cell types in single cell resolved spatial transcriptomics technologies (e.g. MERFISH ^49^ and Seqfish ^50^).
2. Feature-centric perspective: spatial mapping imputes unmeasured gene expression in the spatial domain for techniques that do not achieve full transcriptome coverage (e.g. MERFISH ^51^, seqFISH+^52^).

As previously suggested in the NovoSpaRc method^8^, a variant of OT^17^ can be used to probabilistically map reference cells into the spatial domain. However, previous approaches faced several limitations, including scalability, applicability beyond gene expression reference data, and incorporation of spatial information in the mapping problem. With `moscot.space.mapping`, we introduce a model that applies to both the sample- and feature-centric perspectives, scales to large datasets, and incorporates multi-modal information. Moreover, `moscot.space.mapping` explicitly makes use of spatial information when solving the mapping problem.

Let thus *X* ∈ ℝ*^N^*^×*D*^*^x^* and *Y* ∈ ℝ*^M×Dy^* represent a pair of state matrices for *N* cells and *M* samples (cells, spots, etc.) observed in the dissociated reference and the spatial dataset, respectively. We assume state matrices to represent gene expression for different numbers of genes, *D_x_* for the dissociated reference and *D_y_* for the spatial dataset. We allow further multi-modal information in *X*, e.g. from joint RNA/ATAC reaodut ^20–22, 38^. In addition, let 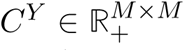 encode spatial similarity among the *M* samples in *Y* (we define *C^X^* below). Depending on the spatial technology, *C^Y^* contains either euclidean distance among spatial locations or similarities in spatial graphs^42, 53^. Optionally, as in `moscot.time`, the user may provide marginal distributions ***a*** ∈ *Δ_N_* and ***b*** ∈ *Δ_M_* over cells in the dissociated reference and samples in the spatial dataset. In the context of `moscot.space.mapping`, these may represent sample-level uncertainties or estimated cell numbers per spot in the spatial dataset for barcoding-based spatial technologies^54–56^.

`moscot.space.mapping`’s key output is a coupling matrix *P* ∈ *U* (***a***, ***b***), linking cells in the dissociated reference with samples in the spatial dataset. In particular, the *i*-th row *P_i,_*: represents the amount of probability mass transported from cell *i* in the reference towards any spatial sample *j*.

These definitions allow us to formalize the aim of `moscot.space.mapping`: we seek to find a coupling matrix *P* ∈ *U* (***a***, ***b***) which relates reference cells with spatial samples such that their distance in the shared transcriptome space is minimized while the correspondence between molecular and spatial similarity is maximized.

#### 1.3.2 Model description

To quantify the global distance between reference and spatial dataset in the shared transcriptome space, we follow `moscot.time` and define a cost function *c*(***x****_i_,* ***y****_j_*) and associated cost matrix 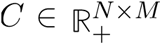. The matrix *C* quantifies expression distance in a shared latent space computed using PCA or scVI ^57^. Note that the shared latent space is constructed using only those genes which have been measured in both the dissociated reference and the spatial dataset.

##### Gromov-Wasserstein for structural correspondence

In their previous NovoSpaRc model, Nitzan et al. ^8^ showed how introducing a structural correspondence assumption between gene expression and spatial information greatly enhanced their ability to accurately solve the spatial mapping problem. In particular, they assume that cell pairs should be coupled such that there is a correspondence between distances in gene expression and distances in physical space. Following their suggestion, we encode the structural correspondence assumption in a *Gromov-Wasserstein* (GW)-type OT problem,

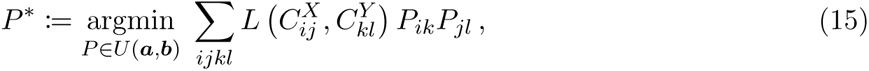

for spatial distance matrix 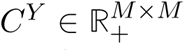, defined as above, and reference distance matrix 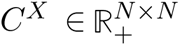, quantifying molecular similarity among cells in the dissociated reference. To compute *C^X^*, we measure expression distance among reference cells in a latent space defined using PCA or scVI^57^. Correspondence between *C^X^* and *C^Y^* is quantified entry-wise using the cost function *L*, which is set to square euclidean by default. This cost is evaluated element-wise, i.e. 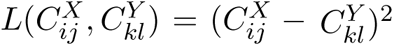.

Intuitively, the GW-type problem aims to find a coupling matrix to maximize the structural correspondence between gene expression and spatial information. Note that individual genes may still show sharp gradients in the spatial domain; the structural correspondence assumption applies to aggregated molecular profiles.

**The** `moscot.space.mapping` **model.** The `moscot.space.mapping` model is a combination of the W-term, quantifying expression distance between the reference and the spatial dataset, with the GW term, quantifying structural correspondence between the reference and the spatial dataset, in a *Fused Gromov-Wasserstein* (FGW)-type OT problem ^17, 58^,

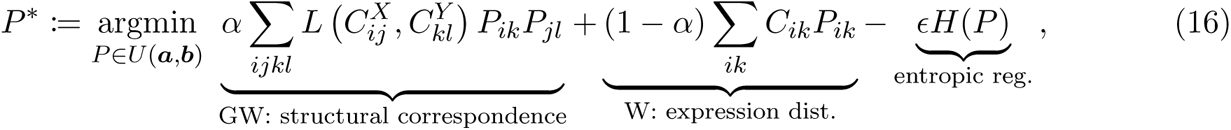

where we have added entropic regularization at strength *ɛ* and introduced the weight parameter *α* to control the relative contribution of W and GW-terms. The objective function contains the following cost matrices:

- 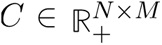 compares reference cells with spatial samples in terms of expression in shared genes.
- 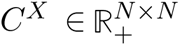 compares reference cells among each other in terms of gene expression.
- 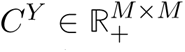 compared spatial samples among each other in terms of spatial distance.

We optimize the `moscot.space.mapping` objective function of Equation (16) using a mirror descent scheme ^17^ (Supplementary Note 1). To account for uneven cell type proportions between the reference and the spatial datasets, we optionally allow for unbalancedness in the FGW-type problem ^59^.

#### 1.3.3 Multi-modal data and scalability

The model presented here is an extension of the NovoSpaRc model^8^, which was restricted to a certain cost function, and only supported feature-centric interpretation. Further, NovoSpaRc is only applicable to uni-modal data and has cubic time- and quadratic memory complexity in the number of cells, largely preventig its application to atlas-scale spatial datsets and references containing multiple modalities. This section extends the `moscot.space.mapping` model to overcome both limitations.

##### Multimodal reference datasets

Multimodal data contains additional information about a cell’s molecular state, which can guide the mapping process. While previous methods could apply a mapping learned from gene expression data to other modalities collected for the same cells ^45^, `moscot.space.mapping` is the first method to make use of multimodal information in the actual mapping problem.

Consider a dissociated reference dataset with multi-modal data matrices, *X*^(R)^ and *X*^(O)^, where R refers to gene expression and O refers to another modality, e.g. chromatin accessibility ^26^ or surface marker expression ^19, 60^. We construct the across-space cost matrix *C* and the spatial cost matrix *C^Y^* as before but modify the construction of the reference cell cost matrix *C^X^*. Similar to `moscot.time`, we concatenate joint representations or use joint latent space learning techniques ^35–37, 61^ to obtain a single molecular representation and measure distances in this representation to define *C^X^*. Our multimodal approach allows learning a more faithful correspondence between molecular similarity in the dissociated reference and spatial proximity in the spatial dataset.

##### Atlas-scale spatial mapping

For squared euclidean loss function *L* and within-space cost functions *C^X^* and *C^Y^*, we implemented `moscot.space.mapping` to have quadratic time and memory consumption by exploiting low-rank properties of the euclidean distance ^62^ (Supplementary Note 2). Similarly to `moscot.time`, solving our FGW-type problem in the backend using OTT^25^ grants us GPU execution and jitting ^39^. While this leads to good performance on datasets of intermediate size (approx. 10k cells in reference and spatial datasets), the quadratic scaling becomes prohibitive for atlas-scale datasets.

To overcome the quadratic time and memory complexities, we make use of a recently proposed low-rank GW formulation ^62^ (Supplementary Note 2) which extends the original low-rank Sinkhorn formulation (Supplementary Note 3). This uniquely enables `moscot.space.mapping` to relate hundreds of thousands of dissociated reference cells to spatial locations.

#### 1.3.4 Downstream applications

`moscot.space.mapping` supports both sample and feature-centric downstream analysis techniques.

##### Annotation-centric perspective

In this perspective, we have cell type or -state labels available in the reference, which we use to map to the spatial dataset. Suppose we are given one-hot encoded reference labels through the matrix *F* ∈ {0, 1}*^N×S^* for *S* cell types or states. We obtain annotated cell types in the spatial domain via the matrix 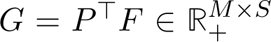. For each spatial sample *j*, the row *G_j,_*: contains the mapped cell type likelihood for each of the *S* cell types or -states. We can then assign discrete cell types to the spatial sample by either taking the argmax of the row or by integrating over probabilities per cell type and taking the maximum value.

##### Feature-centric perspective

In this perspective, we have more genes measured in the dissociated reference than the spatial dataset; we would like to use the mapping problem solution to impute spatial gene expression. This setting is relevant for spatial technologies that do not achieve full transcriptome coverage. Let 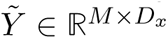 denote inferred expression in the spatial domain; it holds

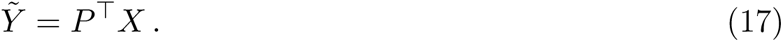

Analogous definitions hold for additional modalities collected in the dissociated reference; for example, we may use Equation (17) to map chromatin accessibility or surface marker expression into spatial coordinates.

To facilitate further downstream analysis of mapped spatial data, `moscot.space.mapping` interfaces with squidpy ^53^, a spatial analysis toolkit containing various visualization and testing capabilities. For example, squidpy can be used to test for spatial enrichment of mapped cell type annotations or to quantify spatial variability of imputed gene expression.

### 1.4 moscot.space.alignment for aligning spatial transcriptomics data

#### 1.4.1 Model rationale, inputs, and outputs

The rapidly increasing amount of spatial datasets poses massive data analysis challenges; in particular, faithful integration of spatial data across tissue slides, individuals, and labs is currently an open problem that limits our ability to study tissue architecture across scales^42, 63^. Different terms exist to refer to spatial integration problems; we prefer to speak of *spatial alignment*. Solving a spatial alignment problem can serve two principal objectives:

1. Joint analysis: aligning spatial datasets against a common coordinate framework ^63^ (CCF) enables us to gain statistical power by jointly considering multiple samples and enables new types of analysis, such as expression variability in space. Aligning data against CCFs will be a crucial step toward spatial atlas building.
2. 3D reconstruction: aligning sequential adjacent tissue sections allows us to build faithful 3D tissue models.

As previously suggested^10^, FGW-type OT^17, 58^ can be used to probabilistically align spatial datasets. However, the previous PASTE method ^10^ was targeted towards small-scale 10x Visium datasets; the authors considered a maximum of 4k spots per sample in their applications ^10, 64^. The scalability of PASTE is limited because it cannot run on GPUs and does not make use of entropic regularization, jitting, and recent low-rank formulations of FGW-type OT^62^. Further, PASTE is limited to adjacent Visium tissue slides from the same individual because it cannot handle varying cell type proportions. Moreover, the approach does not make use of multi-modal molecular readout.

With `moscot.space.alignment`, we present an approach that overcomes these limitations; in particular, `moscot.space.alignment` scales to large and diverse spatial datasets through GPU acceleration, entropic regularization ^1, 29^, jitting ^39^ and low-rank factorizations ^62^. Our approach can integrate samples from different individuals with varying cell type proportions through an unbalanced formulation ^59^ and applies to spatial technologies beyond 10x Visium, including in-situ sequencing (ISS) and in-situ hybridization (ISH)-based assays. Furthermore, our approach makes use of multi-model information where available.

Let thus *X* ∈ ℝ*^N×Dx^* and *Y* ∈ ℝ*^M×Dy^* represent a pair of state matrices for *N* and *M* spatial samples observed in two spatial datasets. We refer to *X* and *Y* as the *left* and *right* datasets, respectively. We assume that state matrices represent gene expression for varying gene numbers *D_x_* and *D_y_*. Optionally, we allow additional multimodal readout at both left and right datasets. In addition, let 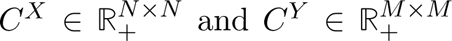 encode spatial similarity among the *N* samples in *X* and the *M* samples in *Y*, defined through e.g. euclidean distance in space or similarities in spatial graphs. Optionally, as in previous moscot models, the user may provide marginal distributions ***a*** ∈ *Δ_N_* and ***b*** ∈ *Δ_M_* over spatial samples in left and right datasets. In the context of `moscot.space.alignment`, these may represent sample-level uncertainties or estimated cell numbers per spot for barcoding-based spatial technologies^54–56^.

`moscot.space.alignment`’s key output is a coupling matrix *P* ∈ *U* (***a***, ***b***) linking spatial samples across the two datasets. In particular, the *i*-th row *P_i,_*: represents the amount of probability mass transported from spatial sample *i* in the left dataset towards any spatial sample *j* in the right dataset.

These definitions allow us to formalize the aim of `moscot.space.alignment`: we seek to find a coupling matrix *P* ∈ *U* (***a***, ***b***) which relates spatial samples across left and right datasets such that their distance in the shared transcriptome space is minimized while the correspondence between spatial arrangements is maximized.

#### 1.4.2 Model description

To quantify the global distance between left and right datasets in the shared transcriptome space, we follow previous moscot models and define a cost function *c*(***x****_i_,* ***y****_j_*) and associated cost matrix 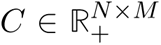 The matrix *C* quantifies expression distance in a shared latent space computed using PCA or scVI^57^. Using the transcriptome-cost matrix *C* in the W-term and the spatial cost matrices *C^X^* and *C^Y^* in the GW-term, we define a FGW-type OT problem ^17, 58^ as for `moscot.space.mapping` (Equation (16)) and solve it using the mirror descent scheme (Supplementary Note 1). For samples with varying cell type proportions, we optionally allow for unbalancedness ^59^.

#### 1.4.3 Multi-modal data and scalability

We include additional multi-modal data collected at left- and right datasets in the W-term; in particular, we follow `moscot.time` and use concatenated representations or joint latent space learning techniques ^35–37, 61^.

We use the same scalability improvements as for `moscot.space.mapping`; in particular, we achieve fast runtimes on datasets of intermediate size through GPU acceleration and jitting ^25, 39^. For atlas-scale left and right datasets, we employ low-rank factorizations to achieve linear time and memory-complexity ^62^ (Supplementary Note 2).

#### 1.4.4 Downstream applications

`moscot.space.alignment` supports both joint analysis of several spatial datasets in a CCF and 3D reconstruction of adjacent tissue sections through different alignment *policies*.

For joint analysis of several spatial datasets, we rely on a pre-defined CCF^63^. To define such a CCF, one may either use a dedicated computational method^65^ or manually designate one spatial sample to serve as the CCF. Given a CCF *X* ∈ R*^N×Dx^* and *R* query datasets *Y* ^(^*^r^*^)^ ∈ ℝ*^Mr ×Dr^* ∀*r* ∈ {1*, …, R*}, `moscot.space.alignment` solves a *star-policy* alignment problem where each query *Y* ^(^*^r^*^)^ is aligned against the central CCF *X*. To enable joint analysis of all query datasets *Y* ^(^*^r^*^)^ in terms of CCF-spatial coordinates, we compute the projection

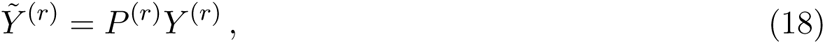

for projected gene expression 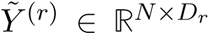 and corresponding coupling matrix 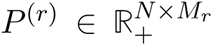. Solving the star-policy alignment problem with `moscot.space.alignment` and projecting into CCF coordinates allows joint analysis of all spatial query samples {*Y* ^(1)^*, …, Y* ^(^*^R^*^)^}.

For 3D reconstruction of adjacent tissue sections, let *X*^(^*^r^*^)^ ∈ ℝ*^Nr ×Dr^* represent gene expression of slide *r* for *N_r_* spatial samples and *D_r_* genes. Let further *Z*^(^*^r^*^)^ ∈ ℝ*^Nr ×^*^2^ represent the corresponding spatial coordinates. We consider *R* sequential slides, *r* ∈ {1*, …, R*}. To align their coordinate systems, `moscot.space.alignment` solves a *sequential policy* alignment problem where each dataset *X*^(^*^r^*^)^ is aligned against the next dataset *X*^(^*^r^*^+1)^ in the sequence. Given the corresponding coupling matrix 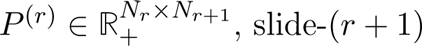, coordinates are transformed into slide-*r* coordinates using

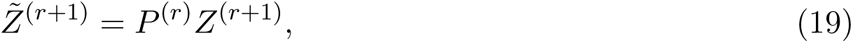

for 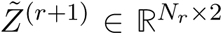. We refer to this as the *warping* transformation because it non-linearly warps *Z*^(^*^r^*^+1)^ coordinates onto *Z*^(^*^r^*^)^ coordinates. Alternatively, `moscot.space.alignment` implements the previously suggested affine-linear transformation^10^. We recommend the warping transformation whenever non-linear effects between adjacent slides are expected. By designating any reference slide *r^∗^*, all other coordinate systems can be transformed into *Z*^(r∗)^ coordinates via sequential application of either the warping or the affine transformation.

In either cases of the alignment problem, it’s possible to further refine the alignment by solving an additional W-type problem on the spatial coordinates.

We interface with squidpy ^53^ for further joint analysis of several spatial datasets in a CCF. For example, squidpy may be used to study expression heterogeneity at a defined spatial location in the CCF across several spatial datasets.

### 1.5 moscot.spatiotemporal to decipher spatiotemporal variation

#### 1.5.1 Model rationale, inputs, and outputs

Cellular state-change processes, including development, regeneration, and reprogramming, do not unfold isolated in single cells but in constant communication with the surrounding tissue ^42^. Recent experimental advances allow for spatially-resolved gene expression measurements at near single-cell resolution across developmental processes; in particular, the StereoSeq ^66^ technology has been applied to various developmental settings ^66–70^. These experiments yield time series of gene expression measurements (as in `moscot.time`), with additional spatial readout at each time point. With `moscot.spatiotemporal` we present the first method to map cells across time points while preserving spatial organization, allowing us to decipher spatiotemporal variation during complex cell-state changes.

Let thus *X* ∈ ℝ*^N×D^* and *Y* ∈ ℝ*^M×D^* represent pairs of state matrices for *N* and *M* spatial samples observed at early (*t*_1_) and late (*t*_2_) time points, respectively. In addition, as stated for `moscot.space.alignment`, let 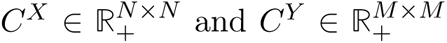 encode spatial similarity among the *N* samples in *X* and the *M* samples in *Y* . Optionally, as in previous moscot models, the user may provide marginal distributions ***a*** ∈ *Δ_N_* and ***b*** ∈ *Δ_M_* over cells at *t*_1_ and *t*_2_. In the context of `moscot.spatiotemporal` these usually correspond to cellular growth-and death rates.

`moscot.spatiotemporal`’s key output is a coupling matrix *P* ∈ *U* (***a***, ***b***) linking samples across the two time points. In particular, the *i*-th row *P_i,_*: represents the amount of probability mass transported from *t*_1_-sample *i* to any *t*_2_-sample *j*.

These definitions allow us to formalize the aim of `moscot.space.mapping`: we seek to find a coupling matrix *P* ∈ *U* (***a***, ***b***) which relates *t*_1_ and *t*_2_-samples such that their distance in the shared transcriptome space is minimized. At the same time, the correspondence between spatial arrangements is maximized.

#### 1.5.2 Model description

We use identical definitions to the `moscot.space.alignment` model, where *t*_1_ samples play the role of the left dataset, and *t*_2_ samples play the role of the right dataset. We adjust the marginals to accommodate cellular growth- and death rates as in the `moscot.time` model, and we optionally allow for unbalancedness ^59^ to handle noisy estimates.

#### 1.5.3 Multi-modal data and scalability

We use the same tricks as in `moscot.space.alignment` to include additional multi-modal readout at *t*_1_ and *t*_2_, and we employ the same strategy to scale our model towards atlas-scale datasets (Supplementary Note 2).

#### 1.5.4 Downstream applications

We extend our model towards more than two time points using the same recipe as in `moscot.time`, and we support all downstream analysis functions introduced for `moscot.time`. We extend the computation of ancestor and descendant probabilities towards spatial regions, i.e., the cell state P of interest in Equation (13) may now represent a spatial region. Thus, `moscot.space.mapping` allows for spatial regionalization to be studied throughout cell-state changes.

We interface with squidpy ^53^ for further downstream analysis of spatiotemporal variation. For example, squidpy may be used to study the spatial enrichment of a mapped cell state of interest across the temporal axis.

## 2 Datasets

### 2.1 Temporal analysis

If not stated otherwise, computations were done using SCANPY^23^ using default parameters. In the temporal setting, we distinguish between driver features and marker features computed with `moscot`. Here, features can be a feature in any modality, for example a gene or a peak. To obtain driver features for a subset of cells, e.g. for a certain cell type, we correlate (Pearson or Spearman) the pull-back distribution of the considered cell type with the corresponding feature, e.g. (processed) gene expression. In practical applications, additionally incorporating the indicator distribution of the pulled back cells (e.g. cell type) of the later time point proved useful. Hence, we obtain marker features by computing the correlation across cells in both the source and the target distribution. If we are interested in marker features for a specific transition, e.g. from a certain cell type to another cell type, we subset the set of cells accordingly.

#### 2.1.1 Application: moscot.time on a mouse embryogenesis atlas

The mouse embryogenesis atlas contains data generated by Mohammed et al. ^71^, Cheng et al. ^72^, Pijuan-Sala et al. ^73^, Qiu et al. ^74^, and Cao et al. ^75^ . These datasets were preprocessed and annotated by Qiu et al. ^74^, and we downloaded them as Seurat objects from http://tome.gs.washington.edu/.

The original authors showed how their embedding computation successfully handled batch effects; thus, we followed their pipeline and reproduced these representations by selecting genes using Seurat v3’s FindVariableFeatures and batch-correcting the data using FindIntegrationAnchors^76^. For further analysis using `moscot.time` in Python, the Seurat objects where transformed into AnnData objects using SeuratData^77^. For the E8.0/8.25 pair of time points, we computed a UMAP ^78^ using the 30-dimensional Seurat PCA latent space and a k-nearest neighbor (k-NN) graph with *k* = 15.

##### Memory and runtime benchmark: moscot.time versus Waddington OT

To investigate method scalability, we ran a memory and runtime benchmark. We selected cells from the E11.5/12.5 time point pair which had the largest number of cells out of all time point pairs: 455,124 cells at E11.5 and 292,726 cells at E12.5. To investigate the dependency of memory and runtime on the number of cells, we generated 11 subsets of increasing size, each containing the same number of cells at E11.5 and 12.5. We increased the cell number in steps of 25,000, up to a maximum of 275,00 cells in either time point.

We compared the performance of three different approaches: Waddington OT^2^ (WOT), `moscot.time` and low-rank `moscot.time`^40^ (Subsubsection 1.2.3). For `moscot.time`, we evaluated the cost function on the fly (online evaluation, Subsubsection 1.2.3) to achieve linear memory complexity. We tested low-rank `moscot.time` for ranks *r* ∈ {50, 200, 500, 2000}. For the memory benchmark, we run all algorithms on CPU, as GPU memory benchmarking is extremely difficult, and memory consumption is likely to be very similar on CPU. For the runtime benchmark, we run the `moscot.time` variants on GPU, but had to run WOT on CPU as it cannot make use of GPU acceleration. For `moscot.time`, we used *ɛ* = 0.005 but had to use a larger value (*ɛ* = 0.05) for WOT as it run into numerical overflow otherwise. Choosing a larger value for *ɛ* makes the problem computationally easier to solve and grants an advantage to WOT.

##### Accuracy benchmark moscot.time versus TOME

We compared the accuracy of the cell transitions inferred by `moscot.time` and Trajectories Of Mammalian Embryogenesis (TOME) ^74^. TOME is a k-NN based algorithm that Qiu et al. ^74^ developed specifically for this dataset. For each *t*_2_-cell *j*, TOME finds the *k* = 5 nearest neighbors at *t*_1_ and treats these as putative ancestors. By aggregating over cell states at both time points, TOME computes weighted ancestor/descendant relationships on the cell-state level. To improve robustness, TOME median-aggregates the inferred edges over 500 randomly subsampled cell sets, each containing 80% of all cells.

Of note, TOME computes neighborhood relationships in a three-dimensional UMAP space, despite the known pitfalls ^79–81^ of low-dimensional non-linear representations. In particular, low-dimensional embeddings like UMAP ^78^ or t-SNE ^82^ do not preserve global data topology well ^83, 84^; trajectories inferred in such spaces are thus prone to suffer from projection artefacts. In addition, TOME is a deterministic approach with no notion of probability mass conservation - cells at *t*_1_ or *t*_2_ can remain without descendants or ancestors, respectively. In contrast, `moscot.time` computes cell-cell distances in a higher dimensional latent space (30-dimensional PCA in this application) - a crucial feature to faithfully describe the data topology of complex developmental state changes. Moreover, `moscot.time` is a probabilistic approach, equipped with a notion of mass conservation grounded on OT.

We applied both `moscot.time` and TOME to all time-point pairs. For `moscot.time` we used *ɛ* = 0.005, initialized marginals using estimated growth-rates ^2^ (Subsection 1.2), set the right unbalancedness parameter *τ_b_* = 1 (i.e., no unbalancedness), and chose the left unbalancedness parameter *τ_a_* such that the predicted amount of apoptotic cells adheres to biologically reasonable ranges ^85–89^.

Specifically, for pre-gastrualtion (E4.5-E6.5), the target apoptotic range was set to 10-15%, for gastrulation (E6.5-E8.5) it was set to 4-6% and for organogenesis (E8.5-E13.5) it was set to 2-4%. For `moscot.time`, we aggregated the cell-level couplings to cell-states transition rates via the pull-back operation of the corresponding cell-state (Subsubsection 1.2.4). These cell state transition rates correspond to the weighted cell state transition edges obtained by TOME, which allows a direct comparison of both approaches.

##### Metrics for the accuracy benchmark: germ-layer and cell-type scores

We developed two metrics to evaluate the accuracy of obtained cell state transitions, one germ-layer and one cell-type metric. First, our germ-layer metric aggregates cell states into germ layers and considers transitions within and across germ layers as correct and incorrect, respectively. This metric is motivated by the observation that cells typically do not cross germ layers ^90^. A prominent exception to this rule is the neural crest, which we excluded from our evaluation ^91^. We followed Qiu et al. ^74^ in classifying cell types into neuroectoderm, surface ectoderm, endoderm, and mesoderm. As in the original study, we excluded transitions between cell types that could not be assigned to a germ layer unambiguously, and transitions with edge weights below 0.05.

Second, our cell-type metric compares every predicted transition with a curated set of allowed transitions. To curate the set of allowed transitions during mouse embryogenesis, we conducted an extensive literature search for all 89 cell types present in the data, identifying previously reported ancestor and descendant states (Supplementary Table 2).

We compute accuracy scores for the germ-layer and cell-type metric by dividing the weighted sum over all transitions satisfying germ-layer boundaries and cell-type restrictions, respectively, by the weighted sum over all transitions included in the evaluation. We mean-aggregated accuracy scores for pre-gastrulation (E3.5-6.5), gastrulation (E6.5-8.5) and organogenesis (E8.5-13.5).

##### Growth rate comparison

Beyond comparisons on the germ-layer and cell-type levels, we wanted to evaluate how `moscot.time` and TOME compare on the single-cell level. However, the TOME method does not output single-cell transitions; it only reports aggregated cell-type transitions. Thus, to still have a baseline, we implemented a variant of the TOME approach in Python, which we call “cell-level TOME” (cl-TOME). Based on the TOME-computed k-NN graph, we consider the 5 nearest neighbors of every *t*_2_-cell *j* as its putative ancestors, and assign each a weight of 0.2. Following the original approach, we increase robustness by repeating the process over 500 randomly subsampled data sets, each containing 80% of the original cells. We sum over the corresponding ancestor weights for each cell, and normalize such that columns in the early-to-late cell transition matrix sum to one. In other words, each *t*_2_ cell *j* receives the same unit mass of incoming transition probability. This interpretation of the k-NN approach allows us to define cell-cell coupling matrices in cl-TOME. Analogously to `moscot.time`, we use pull- and push operations (Subsubsection 1.2.4) to compute ancestors and descendants.

We computed cell-level couplings across time points using `moscot.time` and cl-TOME, excluding extraembryonic tissues to avoid introducing additional variance from the experimental protocol. For `moscot.time` we did not initialize the growth rates using marker genes to enable a fair comparison with cl-TOME, which does not support such initialization. Instead, we ran `moscot.time` with uniform marginals and used unbalancedness to learn growth-rates de-novo. As before, we set *τ_b_* = 1 and chose *τ_a_* such that the resulting predicted fraction of apoptotic cells lies in a biologically reasonable range (Supplementary Table 3) ^87, 88^. For both methods, we calculated growth-rates via the left marginal (row sum) of the corresponding coupling matrix, Σ*_j_ P_ij_*. To avoid overcrowding our histograms of growth rates per cell-type, we only show the 5 cell types with most cells per time point.

To translate these growth rates onto a scale that is biologically interpretable, we adjusted them for the mean population growth from *t*_1_ to *t*_2_. Specifically, we computed the change in population size between two time points, *s* = |*e*_1_|*/*|*e*_2_| where |*e*_1_| and |*e*_2_| represent the estimated cell number at early and late embryo stages, respectively ^71–75^. Next, we scaled the mean growth rate of *t*_1_ cells to match the factor *s*. To obtain an estimate of the apoptosis rate, we calculated, for each cell, the difference between 1 and the scaled growth rate. By summing over these differences for all *t*_1_ cells for which the scaled growth rate was smaller than 1, we calculated the predicted number of dying cells at *t*_1_. We divided the sum by the total number of *t*_1_-cells in the dataset to obtain estimated apoptosis rates. We run the above calculations independently for all time points for `moscot.time` and cl-TOME.

##### Comparison in terms of driver gene correlations

To further assess the cell-level couplings predicted by either method, we reasoned that high correlations between ancestor probabilities and known driver genes for a cell state are indicative of method success. Thus, for the cell states described in the main text, we computed their `moscot.time`- and cl-TOME-predicted ancestor distributions (Subsubsection 1.2.4). To exclude the influence of driver genes involved in unrelated differentiation events, we restricted the correlation computation to known progenitor populations. For each pulled cell state, we curated a list of known driver genes (Supplementary Table 4), filtered the list to contain only highly-variable genes at the corresponding time point, and imputed their expression using scVI’s decoder output using `get_normalized_expression`. After filtering to highly-variable genes, we retained 36 genes for definitive endoderm, 15 genes for allantois cells, 24 genes for the first heart field, and 24 genes for the pancreatic epithelium. We Spearman-correlated these imputed expression values with predicted ancestor distributions using scipy.stats.spearmanr ^92^.

#### 2.1.2 Application: **moscot.time** on multi-model pancreas development

##### Dataset generation

Embryonic pancreata from NVF homozygous mice were collected and pooled together (8 pancreata from E14.5 and 11 pancreata from E15.5). 0.25% Trypsin were added to the samples for 5 min on ice and following incubation at 37 °C for 10 min. Next, the single-cell samples were centrifuged at 1700 rpm (290 x g) for 5 min at 4 °C. After removing the supernatant, cells were counted. 5*µl* rat IgG2a K isotype control (eBioscience, 12-4321-42) and anti-mouse CD326 (EpCAM) PE (eBioscience, 12-5791-81) were used for 1×106 cells (100*µl* total volume). Sample were stained for 30 min at 4 °C following staining with DAPI to detect dead cells. After washing twice and resuspended in FACS buffer (PBS, 1% BSA, 0.5 mM EDTA), the single-cell samples were loaded for FACS sorting. The following gating strategy was used: main population>single-cells>living cells (DAPI negative)>Ngn3+ (FITC+) and EpCAM+ (PE+) cells. Then, cells were counted and Trypan Blue staining was used to identify the number of dead cells. Sample with less than 20% of dead cells were processed for single-cell RNA seq.

##### Preprocessing

We preprocessed the samples independently for gene expression and chromatin accessibility. With respect to gene expression, all cells with mitochondrial gene fraction higher than 0.025 in E14.5 or higher than 0.02 in E15.5 were removed. Moreover, cells with fewer than 2,350 counts or more than 30,000 counts were removed in E14.5, while cells with fewer than 4,000 counts or more than 35,000 counts were removed in E15.5. Doublets were identified using DoubletFinder ^93^, resulting in 9.87% doublets in sample E14.5 and 8.45% doublets in E15.5. After concatenation of the two samples, genes which were detected in fewer than 20 cells were filtered, resulting in 18,768 genes.

Concerning the ATAC modality, all cells in E14.5 with nucleome signal lower than 0.55 or higher than 1.2 were removed. All cells with transcription start site (TSS) enrichment score lower than 4.8 or higher than 8 were filtered out. Additionally, cells were removed if their total open chromatin region count was below 7,000 or above 150,000.

For cells in E15.5 the lower nucleome signal threshold was set to 0.5, while the upper one was set to 1.1. The TSS enrichment scores thresholds were chosen to be 5 and 8.5, while the minimum number of total counts was set to 4,000, and the maximum number was chosen to be 90,000. Peak calling was performed using Signac ^94^. The CellRanger-computed peaks of both samples were concatenated and subsequently pruned. This resulted in 228,259 peaks.

16,978 cells passed quality control in both modalities; 9,852 belonging to E14.5 and 7,126 belonging to E15.5. More cells were filtered in the course of cell type annotation, which is described in the next paragraph, resulting in 9,811 cells for E14.5 and 7,107 cells for E15.5 in the final preprocessed and annotated version of the dataset.

##### Cell type annotation

To construct a weighted nearest neighbor graph, an embedding of both modalities is needed. Therefore, before performing a Principal Component Analysis (PCA, 50 dimensions) on the log1p-tranformed gene expression data, the count data was normalized using SCTransform ^95^ and cell cycle genes and ambient genes were discarded. Ambient genes were identified using DropletUtils ^96^. The ATAC data was processed by term frequency-inverse document frequency (tf-idf) normalisation followed by singular-value decomposition using Signac, computing the first 50 singular components. Due to high correlation with the sequencing depth, the first and the fifth components were removed. Having computed the embeddings for GEX and ATAC, respectively, we constructed a weighted nearest neighbor graph in MUON, and used it for multi-modal, unsupervised clustering. If not stated otherwise, this is also the graph based on which we computed UMAPs ^78^.

Annotation was performed based on the expression of marker genes as reported in previous studies ^97–102^ (Supplementary Table 9) and cell cycle scores for the proliferating populations computed with `scanpy.tl.score_genes_cell_cycle`. It is important to mention that we identified a cluster branching off the Ngn3 high population, which we found to express similar genes as a cluster called Fev+ epsilon in Bastidas-Ponce et al. ^98^. In fact, neither the cluster reported in Bastidas-Ponce et al. ^98^ nor the cluster found in our new dataset has a substantially high expression of *Fev* (Supplementary Fig. 21). Hence, we labeled this cluster as *epsilon progenitors*.

To arrive at the finer resolution of cell types as shown in Figure 3e, subclustering was performed on the same neighborhood graph (incorporating both modalities).

##### moscot.time model

We computed the cost matrix defining the OT problem between E14.5 and E15.5 using geodesic distance along the kNN graph (Supplementary Note 5). The solution was computed using the `HeatFilter` provided by PyGSP^103^. The embedding based on which the underlying graph (connectivities as obtained from `scanpy.pp.neighbors`,^23^) is calculated was obtained by MultiVI (^36^) with a Poisson likelihood as introduced in PoissonATAC ^104^. PoissonATAC was run with default parameters. This graph was constructed on a different embedding than the one we used for unsupervised clustering to reduce the bias towards one embedding.

`Two moscot` models were run based on the weighted nearest neighbor graph whose construction is described above. First, a model was run on the full dataset. The `moscot.time` model was run with default parameters, i.e. in a balanced manner with uniform marginals. In detail, the regularization parameter *ɛ* was set to 10*^−^*^3^ and the cost matrix was scaled by its mean. Uniform marginals were chosen as the large abundance of highly proliferating Ductal cells would have marginalized the influence of less abundant cell types. It is also important to note that the dataset is FACS-sorted and hence proportions of initially sequenced cells are highly biased and do not reflect the true cell type distribution. This also causes the final model to predict descendants (ancestors) across one day (wall clock time). In effect, the directionality of the developmental process is kept, while its magnitude does not reflect ground truth biological progress.

For the analysis of the endocrine branch the optimal transport solution was computed on a reduced dataset, only containing endocrine cells (alpha, beta, delta, epsilon) and their progenitors (cell types labelled as Fev+ alpha, Fev+ beta, Fev+ delta, epsilon progenitors, Fev+, Ngn3 high, Ngn3 high cycling, Ngn3 low). Again, uniform marginals were chosen as the proliferation and apoptosis scores obtained from *TemporalProblem.score_genes_for_marginals* are almost constant. The optimal transport solution was computed with standard parameters, i.e. the same parameters as for the first model.

##### Marker feature and driver feature analysis with moscot.time

We compute marker features and driver features as described in 2.1. Moreover, when analyzing the transition from epsilon to alpha cells, we excluded the Fev+ alpha population (the main progenitor cell type) from the set of considered cell types. This helped to find genes which are particularly activated in the epsilon cells.

##### Marker regions of chromatin accessibility

To identify marker regions of chromatin accessibility, a Wilcoxon test was run by calling `FindMarkers` provided by Seurat. The test was performed with default settings and the considered cell type was run with respect to all remaining cell types (subset to endocrine cells and endocrine progenitors, i.e. Ngn3 low, Ngn3 high, epsilon progenitors, Fev+, Fev+ delta, Fev+ alpha, Fev+ beta, alpha, beta, delta, epsilon).

##### Motif analysis

Motif data was downloaded from cisBP ^105^. Position weight matrices (PWMs) and corresponding visualizations as well as metadata was downloaded as a *Bulk download* after filtering by species (*mus musculus*) on 01/03/2023. cisBP contains data from both experimentally measured binding activities and inferred ones (e.g. from other species). Transcription factors with DNA-binding domain amino acid similarity above a certain threshold (defined for each DNA-binding domain class separately and provided by cisBP) are hence also considered as binding candidates.

We define a transcription factor to have an association with a motif if it is either directly measured or is inferred and has a sufficiently high DNA-binding domain amino acid similarity, i.e. is reported as such by cisBP. This way, one motif can have an association with multiple transcription factors and one transcription factor can have an association with multiple motifs.

To obtain motif scores on a single-cell level, chromVAR was run using the API provided by Signac. In effect, `AddMotifs` was called, followed by `RunChromVAR`. To obtain marker motifs with `moscot`, we consider the temporal order of gene expression and activity of a motif. In fact, moscot comes with a list of transcription factors for different species (human, mouse, drosophila), obtained from the SCENIC+ database^106^. Thus, we compute marker transcription factors using moscot’s capability to compute marker features. Moreover, we perform a differential motif activity test (Wilcoxon test using scanpy’s `rank_genes_groups`) based on the ChromVAR scores. Subsequently, we find our marker motifs by combining these two sources of information. Therefore, we only keep marker TFs for which we have an associated transcription factor.

### 2.2 Spatial analysis

#### 2.2.1 Benchmark: moscot.space.mapping across a range of spatial datasets

We benchmarked moscot’s *MappingProblem* against two state of the arts methods: Tangram^45^ and gimVI^107^, as implemented in scVI tools ^18^. We employed the datasets collected by Li et al. ^44^. We chose all datasets from Li et al. that we were able to reprocess, result in 14 ones considered for the benchmark. Furthermore, in contrast to the original benchmark, we do not employ the single-cell dataset as reference, since we are not confident that such data represents a faithful ground truth for method comparison. Therefore, we split the spatial dataset in 50% of the data points treated as single-cell reference and 50% treated as spatial data. We also explicitly maintain the data type at input consistent with model requirements, therefore we normalize and scale counts for both moscot and Tangram and we keep raw unnormalized counts for gimVI. We randomly hold out 100 genes if the total number of genes in a dataset is > 2000, otherwise we hold out 10 genes. We train models on the remaining genes and evaluate performance based on Spearman correlation. We report the mean Spearman correlations across 3 random seeds (including random seeds both for dataset split and initialization/training routines). For some datasets, Tangram or gimVI could not be run either due to time complexity (we set a maximum budget of 5 gpu/hours for each method to run) or errors of the models (e.g. inability to match gene ids between train and impute data). The budget for hyperparameters was the same (6 configurations) for each model. Specifically, we ran the sweep on these parameters:

- moscot: *epsilon* (entropy regularization parameter) and *alpha* (interpolation parameter between W-term and GW-term).
- Tangram: *learning rate* and *number of epochs*.
- gimVI: *number of epochs* and *number of latent dimensions*.

We also report memory and time complexity for each algorithm across datasets and seeds. All experiments were run on GPU on the Helmholtz Cluster (mix of V100 an A100 GPUs). Benchmarks were run using SEML^108^.

##### Spatial correspondence

Spatial correspondence is computed as follows: First, we compute *n* increasing spatial distance (Euclidean) thresholds between all data points in the dataset. Then, at each threshold level, we compute the gene expression similarity (Euclidean distance) between all genes in all the spots whose (Euclidean) distance is below the selected threshold. Spatial correspondence is then calculated as the Pearson correlation between gene expression similarity and the spatial distance thresholds. The method is available in `moscot.problems.space.MappingProblem.spatial_correspondence`.

#### 2.2.2 Application: moscot.space.mapping on the liver

We applied `moscot`.space’s *MappingProblem* to the mouse liver dataset from Vizgen MERSCOPE downloaded from https://vizgen.com/data-release-program/. We processed the dataset following standard scanpy ^23^ and squidpy ^53^ processing. For the single-cell reference, we downloaded the CITE-seq dataset from here: https://www.livercellatlas.org/. The dataset was first reported in Guilliams et al. ^109^. We used the *MappingProblem* in the follwing way: we used the set of 336 common genes for the linear term while in the quadratic term we used the PCA of gene expression for the single-cell reference dataset and the PCA gene expression concatenated to the spatial coordinates for the spatial dataset. We then performed the gene expression and protein imputation by computing the barycentric projection of protein expression to the spatial dataset (Subsection 1.3). The same barycentric projection approach was also used to transfer annotations of cell types from the single-cell reference to the spatial dataset.

#### 2.2.3 Benchmark: moscot.space.alignment on simulated data

We benchmarked moscot’s *AlignmentProblem* against two other state-of-the-art alignment methods: PASTE^10^ and GPSA ^65^. We chose the same computational budgets across all methods, that is 12 unique sets of hyperparameters:

- Moscot: *epsilon* (entropy regularization parameter) and *alpha* (interpolation parameter between W-term and GW-term).
- PASTE: *alpha* (interpolation parameter between W-term and GW-term) and *norm* (scaling of the cost matrix). Please refer to PASTE^10^.
- GPSA: *kernel* (kernel for the Gaussian Process), *n_epochs* (number of epochs) and *lr* (learning rate). Please refer to GPSA ^65^.

Due to the inability to run GPSA on GPU, we ran all methods on CPU. We generated four synthetic datasets based on the data generation described in^65^. In short, samples from random normal distribution are generated to build a synthetic gene expression file arranged in a grid. Points are then randomly subsampled by a fraction of 0.7, 0.8, 0.9 of the original datasets, so that the total number of points do not match in the source and target dataset. This is a similar benchmark settings as the one proposed by Jones et al. ^65^ and Zeira et al. ^10^, yet to make all three methods comparable we decided to use the barycentric projection 17 of spatial coordinates with respect to the coupling for both PASTE^10^ and `moscot`. Because of the low sample size of the experiments, we ran moscot in full rank mode (as opposed to low rank mode). Larger dataset, such as the one analyzed in the main text, would be prohibitively large for both PASTE and GPSA.

#### 2.2.4 Application: moscot.space.alignment on mouse brain coronal sections

We applied moscot’s *AlignmentProblem* to a large scale MERFISH dataset from Vizgen Merscope https://vizgen.com/data-release-program/, specifically, to two sections of the mouse coronal brain. We aligned three samples from three different mice for each section. We performed the first alignment with the moscot’s *AlignmentProblem*, in the “affine” mode (Subsection 1.4 and Equation (19)). Thus, we obtained 2 of the 3 slices aligned to the remaining one, chosen as reference. Furthermore, we performed a second alignment on FGW-aligned coordinates with a W-type problem to obtain an improved warped alignment. This turned out to prove useful in low rank settings. We performed the same operations for both triplets of coronal sections.

##### Gene consistency analysis of aligned slices

We assessed the quality of the alignment based on gene expression only, as we do not have cell type annotations for the brain sections of interest. To this end, we computed the neighbor graph in the aligned space with squidpy ^53^ (*knn mode* with at least 30 neighbors for each observation). Then, for each gene, we filtered cells with no expression, and retrieved neighbors of the reference section (0) from the two other sections (1 and 2). We then assessed the gene expression histogram across all cells in the query sections that are neighbors in the reference section, and reported the expression of the gene of interest. We performed this analysis across all genes and reported the L1 Wasserstein distance between gene expression histograms. A low L1 Wasserstein distance between the gene expression density of the query section and the gene expression density of the reference section means that the set of cells in the reference is similar to the matches cells of the aligned section. On the opposite, if the L1 Wasserstein distance is high, it means that neighboring cells in query and reference slides are not similar in gene expression distribution, highlighting a potential mismatch in the alignment. It should be noted that a source of such a mismatch could also be the intrinsic biological variability between tissue sections. Nevertheless, because we don’t have access to tissue sections annotations we decided to use the gene expression similarity metric described above to quantitatively evaluate the alignment. We further evaluated whether the distribution of L1 Wasserstein distance between query and reference sections showed a correlation with the mean expression of the gene. We did not observe a strong association highlighting the fact that this analysis is robust to gene expression variability. All results are reported in Supplementary Figure 10.

### 2.3 Spatiotemporal analysis

#### 2.3.1 Application: moscot.spatiotemporal on mouse embryogenesis StereoSeq data

##### Preprocessing

We used the mouse embryogenesis StereoSeq data generated by^66^. The data was preprocessed and annotated by Chen et al. ^66^ and available to be downloaded as AnnData objects from https://db.cngb.org/stomics/mosta/. In the reported analysis, for full embryo mapping, we use “Mouse_embryo_all_stage.h5ad”, a file containing a single slide for each time point. This file was also used to extract brain cells from early time points. For the latest time point, E16.5, we used the detailed brain annotation slide given in “16.5_E1S3_cell_bin_whole_brain.h5ad”. For each section, we used the “count” layer and performed standard preprocessing with scanpy. We filtered cells (min_genes = 200) and genes (min_cells = 3), normalized cell counts and log-transformed the data.

To perform analysis over brain cells, and transfer the cell type annotation from E16.5 to earlier time points (E13.5-E15.5), we extracted cells annotated as “Brain” from the full embryo AnnData object and merged with the E16.5 annotated brain AnnData object.

##### Mapping accuracy

We used `moscot.spatiotemporal` on each time pair of the data and calculated cell state transition rates. We compared the accuracy to `moscot.time` using the germ-layer and cell type transition accuracy (see 2.1). In both cases we fixed epsilon (*ɛ* = 1*e* − 3), used the low-rank approach (rank= 500, *γ* = 10) and evaluated using biologically informed priors based on growth-and death rate modeling computed by `moscot`. For `moscot.spatiotemporal`we also performed a grid search for the interpolation parameter, *α* ∈ {0, 0.2, 0.4, 0.6, 0.8, 0.9, 0.99}.

##### Mapping annotations across time points

We used the detailed cell type annotation provided for E16.5 brain to infer annotations of earlier time points. We mapped cells across time points using the optimal *α* identified in the full embryo analysis and a higher rank (rank=10000), now possible as we are considering a sub-population of the cells. To obtain the cell annotation we start from the last couple (E15.5, E16.5) and used the `moscot.spatiotemporal` cell-transition matrix aggregated over cell types. We assigned each cell at E15.5 with the most probable cell type. Once we had the annotations for E15.5 we repeated this procedure towards earlier time points. To evaluate the accuracy of the annotations we used Scanpy’s^23^ `rank_genes_groups` with respect to the inferred annotations. For each cell type we queried whether the marker genes reported by Chen et al. ^66^ are within the top 50 ranked genes. We reported the percentage of cells types for which this condition holds.

##### CellRank analysis

We used CellRank ^110^, a computational fate mapping tool, to infer marker genes associated with terminal cell states. To define the CellRank kernel, *K*, a matrix containing cells from all time points which is used to obtain the transition probabilities between cells we followed these steps:

1. Obtain a sparse representation of the `moscot.spatiotemporal` cell transition maps. These cell transition matrices occupy the superdiagonal of *K* as they transport cells from early to late time points.
2. Compute the transition matrices within each time point based on gene expression similarity. These values occupy the diagonal of *K*.
3. Combine the above with weights 0.9 and 0.1, respectively to obtain *K*.
4. Row normalize *K*.

We used CellRank’s GPCCA estimator ^111, 112^ to compute terminal states, independently, for the full embryo and brain cells. We defined each terminal state by assigning the most likely 30 cells to it. We computed absorption probabilities on the Markov chain towards these combined cell sets per terminal state group, and interpreted these as fate probabilities. We correlated each gene’s expression with the computed fate probabilities across all cells. We identified the top 20 most strongly correlated genes and transcription factors per terminal cell group. The list of mice transcription factors was downloaded from AnimalTFDB.

## References

1. Buenrostro, J. D. et al. Single-cell chromatin accessibility reveals principles of regulatory variation. Nature 523, 486–490 (2015).

2. Chen, K. H., Boettiger, A. N., Moffitt, J. R., Wang, S. & Zhuang, X. RNA imaging. Spatially resolved, highly multiplexed RNA profiling in single cells. Science 348, aaa6090 (2015).

3. Chen, A. et al. Spatiotemporal transcriptomic atlas of mouse organogenesis using DNA nanoball-patterned arrays. Cell 185, 1777–1792.e21 (2022).

4. Schiebinger, G. et al. Optimal-Transport Analysis of Single-Cell Gene Expression Identifies Developmental Trajectories in Reprogramming. Cell 176, 928–943.e22 (2019).

5. Mittnenzweig, M. et al. A single-embryo, single-cell time-resolved model for mouse gastrulation. Cell 184, 2825–2842.e22 (2021).

6. Rao, A., Barkley, D., França, G. S. & Yanai, I. Exploring tissue architecture using spatial transcriptomics. Nature 596, 211–220 (2021).

7. Palla, G., Fischer, D. S., Regev, A. & Theis, F. J. Spatial components of molecular tissue biology. Nat. Biotechnol. (2022) doi:10.1038/s41587-021-01182-1.

8. Peyré, G. & Cuturi, M. Computational Optimal Transport. Preprint at https://doi.org/10.1561/9781680835519 (2019).

9. Villani, C. Optimal Transport. (Springer Berlin Heidelberg).

10. Tong, A., Huang, J., Wolf, G., Van Dijk, D. & Krishnaswamy, S. Trajectorynet: A dynamic optimal transport network for modeling cellular dynamics. in International Conference on Machine Learning 9526–9536 (PMLR, 2020).

11. Yang, K. D. et al. Predicting cell lineages using autoencoders and optimal transport. PLoS Comput. Biol. 16, e1007828 (2020).

12. Nitzan, M., Karaiskos, N., Friedman, N. & Rajewsky, N. Gene expression cartography. Nature 576, 132–137 (2019).

13. Zeira, R., Land, M., Strzalkowski, A. & Raphael, B. J. Alignment and integration of spatial transcriptomics data. Nat. Methods 19, 567–575 (2022).

14. Cuturi, M. Sinkhorn Distances: Lightspeed Computation of Optimal Transportation Distances. arXiv [stat.ML] (2013).

15. Peyré, G., Cuturi, M. & Solomon, J. Gromov-Wasserstein Averaging of Kernel and Distance Matrices. in Proceedings of The 33rd International Conference on Machine Learning (eds. Balcan, M. F. & Weinberger, K. Q.) vol. 48 2664–2672 (PMLR, 2016).

16. Qiu, C. et al. Systematic reconstruction of cellular trajectories across mouse embryogenesis. Nat. Genet. 54, 328–341 (2022).

17. Demetci, P., Santorella, R., Sandstede, B., Noble, W. S. & Singh, R. SCOT: Single-Cell Multi-Omics Alignment with Optimal Transport. J. Comput. Biol. 29, 3–18 (2022).

18. Gayoso, A., et al. scvi-tools: a library for deep probabilistic analysis of single-cell omics data. bioRxiv 2021.04.28.441833 (2021) doi:10.1101/2021.04.28.441833.

19. Lopez, R. et al. DestVI identifies continuums of cell types in spatial transcriptomics data. Nat. Biotechnol. 40, 1360–1369 (2022).

20. Gayoso, A. et al. Joint probabilistic modeling of single-cell multi-omic data with totalVI. Nat. Methods 18, 272–282 (2021).

21. Ashuach, T., Reidenbach, D. A., Gayoso, A. & Yosef, N. PeakVI: A deep generative model for single-cell chromatin accessibility analysis. Cell Rep Methods 2, 100182 (2022).

22. Cuturi, M., et al. Optimal Transport Tools (OTT): A JAX Toolbox for all things Wasserstein. arXiv [cs.LG] (2022).

23. Scetbon, M., Cuturi, M. & Peyré, G. Low-Rank Sinkhorn Factorization. arXiv [stat.ML] (2021).

24. Scetbon, M., Peyré, G. & Cuturi, M. Linear-Time Gromov Wasserstein Distances using Low Rank Couplings and Costs. arXiv [cs.LG] (2021).

25. Wolf, F. A., Angerer, P. & Theis, F. J. SCANPY: large-scale single-cell gene expression data analysis. Genome Biol. 19, 15 (2018).

26. Virshup, I. et al. The scverse project provides a computational ecosystem for single-cell omics data analysis. Nat. Biotechnol. (2023) doi:10.1038/s41587-023-01733-8.

27. Stoeckius, M. et al. Simultaneous epitope and transcriptome measurement in single cells. Nat. Methods 14, 865–868 (2017).

28. Vayer, T., Chapel, L., Flamary, R., Tavenard, R. & Courty, N. Fused Gromov-Wasserstein Distance for Structured Objects. Algorithms 13, 212 (2020).

29. Ashuach, T., Gabitto, M. I., Jordan, M. I. & Yosef, N. MultiVI: deep generative model for the integration of multi-modal data. *bioRxiv* 2021.08.20.457057 (2021) doi:10.1101/2021.08.20.457057.

30. Lotfollahi, M., Litinetskaya, A. & Theis, F. J. Multigrate: single-cell multi-omic data integration. bioRxiv 2022.03.16.484643 (2022) doi:10.1101/2022.03.16.484643.

31. Cao, Z.-J. & Gao, G. Multi-omics single-cell data integration and regulatory inference with graph-linked embedding. Nat. Biotechnol. 40, 1458–1466 (2022).

32. Tu, X., Cao, Z.-J., Xia, C.-R., Mostafavi, S. & Gao, G. Cross-Linked Unified Embedding for cross-modality representation learning. (2022).

33. Frostig, R., Johnson, M. & Leary, C. Compiling machine learning programs via high-level tracing. (2018).

34. Virshup, I., Rybakov, S., Theis, F. J., Angerer, P. & Alexander Wolf, F. anndata: Annotated data. bioRxiv 2021.12.16.473007 (2021) doi:10.1101/2021.12.16.473007.

35. Lange, M. et al. CellRank for directed single-cell fate mapping. Nat. Methods 19, 159–170 (2022).

36. Manova, K. et al. Apoptosis in mouse embryos: elevated levels in pregastrulae and in the distal anterior region of gastrulae of normal and mutant mice. Dev. Dyn. 213, 293–308 (1998).

37. Martínez-Lagunas, K. et al. In vivo detection of programmed cell death during mouse heart development. Cell Death Differ. 27, 1398–1414 (2020).

38. Lewis, S. L. & Tam, P. P. L. Definitive endoderm of the mouse embryo: formation, cell fates, and morphogenetic function. Dev. Dyn. 235, 2315–2329 (2006).

39. Biancalani, T. et al. Deep learning and alignment of spatially resolved single-cell transcriptomes with Tangram. Nat. Methods 18, 1352–1362 (2021).

40. Lopez, R., et al. A joint model of unpaired data from scRNA-seq and spatial transcriptomics for imputing missing gene expression measurements. arXiv *[cs.LG]* (2019).

41. Li, B. et al. Benchmarking spatial and single-cell transcriptomics integration methods for transcript distribution prediction and cell type deconvolution. Nat. Methods 1–9 (2022).

42. Guilliams, M. et al. Spatial proteogenomics reveals distinct and evolutionarily conserved hepatic macrophage niches. Cell 185, 379–396.e38 (2022).

43. MERSCOPE Spatial Transcriptomics. Vizgen https://vizgen.com/products/ (2021).

44. Cunningham, R. P. & Porat-Shliom, N. Liver Zonation - Revisiting Old Questions With New Technologies. Front. Physiol. 12, 732929 (2021).

45. Sun, T. et al. AXIN2+ Pericentral Hepatocytes Have Limited Contributions to Liver Homeostasis and Regeneration. Cell Stem Cell 26, 97–107.e6 (2020).

46. Horvath, B. et al. Measurement of von Willebrand factor as the marker of endothelial dysfunction in vascular diseases. Exp. Clin. Cardiol. 9, 31–34 (2004).

47. Rood, J. E. et al. Toward a Common Coordinate Framework for the Human Body. Cell 179, 1455–1467 (2019).

48. Jones, A., William Townes, F., Li, D. & Engelhardt, B. E. Alignment of spatial genomics and histology data using deep Gaussian processes. bioRxiv 2022.01.10.475692 (2022) doi:10.1101/2022.01.10.475692.

49. Bruneau, B. G. Signaling and transcriptional networks in heart development and regeneration. Cold Spring Harb. Perspect. Biol. 5, a008292 (2013).

50. Chen, Y. et al. The Role of Tbx20 in Cardiovascular Development and Function. Front Cell Dev Biol 9, 638542 (2021).

51. Honkoop, H. et al. Single-cell analysis uncovers that metabolic reprogramming by ErbB2 signaling is essential for cardiomyocyte proliferation in the regenerating heart. Elife 8, (2019).

52. Fukuda, R. et al. Metabolic modulation regulates cardiac wall morphogenesis in zebrafish. Elife 8, (2019).

53. Guo, Y. & Pu, W. T. Cardiomyocyte Maturation: New Phase in Development. Circ. Res. 126, 1086–1106 (2020).

54. Chodelkova, O., Masek, J., Korinek, V., Kozmik, Z. & Machon, O. Tcf7L2 is essential for neurogenesis in the developing mouse neocortex. Neural Dev. 13, 8 (2018).

55. Han, F., Liu, Y., Huang, J., Zhang, X. & Wei, C. Current Approaches and Molecular Mechanisms for Directly Reprogramming Fibroblasts Into Neurons and Dopamine Neurons. Front. Aging Neurosci. 13, 738529 (2021).

56. Treutlein, B. et al. Dissecting direct reprogramming from fibroblast to neuron using single-cell RNA-seq. Nature 534, 391–395 (2016).

57. Sagner, A. et al. A shared transcriptional code orchestrates temporal patterning of the central nervous system. PLoS Biol. 19, e3001450 (2021).

58. Aiken, J., Buscaglia, G., Bates, E. A. & Moore, J. K. The α-Tubulin gene TUBA1A in Brain Development: A Key Ingredient in the Neuronal Isotype Blend. J Dev Biol 5, (2017).

59. Tessarin, G. W. L. et al. A Putative Role of Teneurin-2 and Its Related Proteins in Astrocytes. Front. Neurosci. 13, 655 (2019).

60. Parra, A. S. & Johnston, C. A. Emerging Roles of RNA-Binding Proteins in Neurodevelopment. J Dev Biol 10, (2022).

61. Runge, K. et al. Disruption of NEUROD2 causes a neurodevelopmental syndrome with autistic features via cell-autonomous defects in forebrain glutamatergic neurons. Mol. Psychiatry 26, 6125–6148 (2021).

62. Han, J. et al. Concerted action of Msx1 and Msx2 in regulating cranial neural crest cell differentiation during frontal bone development. Mech. Dev. 124, 729–745 (2007).

63. Lee, K.-W. et al. PRRX1 is a master transcription factor of stromal fibroblasts for myofibroblastic lineage progression. Nat. Commun. 13, 2793 (2022).

64. La Manno, G. et al. Molecular architecture of the developing mouse brain. Nature 596, 92–96 (2021).

65. Bastidas-Ponce, A. et al. Comprehensive single cell mRNA profiling reveals a detailed roadmap for pancreatic endocrinogenesis. Development 146, (2019).

66. Scavuzzo, M. A. et al. Endocrine lineage biases arise in temporally distinct endocrine progenitors during pancreatic morphogenesis. Nat. Commun. 9, 3356 (2018).

67. Duvall, E. et al. Single-cell transcriptome and accessible chromatin dynamics during endocrine pancreas development. Proc. Natl. Acad. Sci. U. S. A. 119, e2201267119 (2022).

68. Yu, X.-X. et al. Sequential progenitor states mark the generation of pancreatic endocrine lineages in mice and humans. Cell Res. 31, 886–903 (2021).

69. Arnes, L., Hill, J. T., Gross, S., Magnuson, M. A. & Sussel, L. Ghrelin expression in the mouse pancreas defines a unique multipotent progenitor population. PLoS One 7, e52026 (2012).

70. Duong, T. E. et al. A single-cell regulatory map of postnatal lung alveologenesis in humans and mice. Cell Genom 2, (2022).

71. Ranzoni, A. M. et al. Integrative Single-Cell RNA-Seq and ATAC-Seq Analysis of Human Developmental Hematopoiesis. Cell Stem Cell 28, 472–487.e7 (2021).

72. Ma, S. et al. Chromatin Potential Identified by Shared Single-Cell Profiling of RNA and Chromatin. Cell 183, 1103–1116.e20 (2020).

73. Bastidas-Ponce, A., Scheibner, K., Lickert, H. & Bakhti, M. Cellular and molecular mechanisms coordinating pancreas development. Development 144, 2873–2888 (2017).

74. Bergen, V., Lange, M., Peidli, S., Wolf, F. A. & Theis, F. J. Generalizing RNA velocity to transient cell states through dynamical modeling. Nat. Biotechnol. 38, 1408–1414 (2020).

75. Shih, H. P., Wang, A. & Sander, M. Pancreas organogenesis: from lineage determination to morphogenesis. Annu. Rev. Cell Dev. Biol. 29, 81–105 (2013).

76. Xiafukaiti, G. et al. MafB Is Important for Pancreatic β-Cell Maintenance under a MafA-Deficient Condition. Mol. Cell. Biol. 39, (2019).

77. Veres, A. et al. Charting cellular identity during human in vitro β-cell differentiation. Nature 569, 368–373 (2019).

78. Cuesta-Gomez, N. et al. Characterization of stem-cell-derived islets during differentiation and after implantation. Cell Rep. 40, 111238 (2022).

79. Dutto, I., Tillhon, M., Cazzalini, O., Stivala, L. A. & Prosperi, E. Biology of the cell cycle inhibitor p21(CDKN1A): molecular mechanisms and relevance in chemical toxicology. Arch. Toxicol. 89, 155–178 (2015).

80. Zhang, J., McKenna, L. B., Bogue, C. W. & Kaestner, K. H. The diabetes gene Hhex maintains δ-cell differentiation and islet function. Genes Dev. 28, 829–834 (2014).

81. Yosef, N. & Regev, A. Impulse control: temporal dynamics in gene transcription. Cell 144, 886–896 (2011).

82. Liu, C. et al. Spatiotemporal mapping of gene expression landscapes and developmental trajectories during zebrafish embryogenesis. Dev. Cell 57, 1284–1298.e5 (2022).

83. Wang, M. et al. High-resolution 3D spatiotemporal transcriptomic maps of developing Drosophila embryos and larvae. Developmental Cell vol. 57 1271–1283.e4 Preprint at https://doi.org/10.1016/j.devcel.2022.04.006 (2022).

84. Qiu, X., et al. Spateo: multidimensional spatiotemporal modeling of single-cell spatial transcriptomics. bioRxiv 2022.12.07.519417 (2022) doi:10.1101/2022.12.07.519417.

85. Cao, K., Gong, Q., Hong, Y. & Wan, L. A unified computational framework for single-cell data integration with optimal transport. Nat. Commun. 13, 7419 (2022).

86. Chen, W. S. et al. Uncovering axes of variation among single-cell cancer specimens. Nat. Methods 17, 302–310 (2020).

87. Tong, A., et al. Diffusion Earth Mover’s Distance and Distribution Embeddings. ArXiv (2021).

88. Cang, Z. et al. Screening cell–cell communication in spatial transcriptomics via collective optimal transport. Nat. Methods 20, 218–228 (2023).

89. Cang, Z. & Nie, Q. Inferring spatial and signaling relationships between cells from single cell transcriptomic data. Nat. Commun. 11, 2084 (2020).

90. Makkuva, A. V., Taghvaei, A., Oh, S. & Lee, J. D. Optimal transport mapping via input convex neural networks. arXiv [cs.LG] (2019).

91. Bunne, C., Papaxanthos, L., Krause, A. & Cuturi, M. Proximal Optimal Transport Modeling of Population Dynamics. in Proceedings of The 25th International Conference on Artificial Intelligence and Statistics (eds. Camps-Valls, G., Ruiz, F. J. R. & Valera, I.) vol. 151 6511–6528 (PMLR, 28--30 Mar 2022).

92. Bunne, C., et al. Learning Single-Cell Perturbation Responses using Neural Optimal Transport. bioRxiv 2021.12.15.472775 (2021) doi:10.1101/2021.12.15.472775.

93. Bunne, C., Krause, A. & Cuturi, M. Supervised Training of Conditional Monge Maps. arXiv [cs.LG] (2022).

94. Uscidda, T. & Cuturi, M. The Monge Gap: A Regularizer to Learn All Transport Maps. arXiv [cs.LG] (2023).

95. Rood, J. E., Maartens, A., Hupalowska, A., Teichmann, S. A. & Regev, A. Impact of the Human Cell Atlas on medicine. Nat. Med. 28, 2486–2496 (2022).

96. Becht, E. et al. Dimensionality reduction for visualizing single-cell data using UMAP. Nat. Biotechnol. (2018) doi:10.1038/nbt.4314.

97. McInnes, L., Healy, J. & Melville, J. UMAP: Uniform Manifold Approximation and Projection for Dimension Reduction. arXiv [stat.ML] (2018).

98. England, J., Pang, K. L., Parnall, M., Haig, M. I. & Loughna, S. Cardiac troponin T is necessary for normal development in the embryonic chick heart. J. Anat. 229, 436–449 (2016).

99. Nelson, D. O. et al. Irx4 Marks a Multipotent, Ventricular-Specific Progenitor Cell. Stem Cells 34, 2875–2888 (2016).

100. la O Sean, D., et al. Single-Cell Multi-Omic Roadmap of Human Fetal Pancreatic Development. bioRxiv 2022.02.17.480942 (2022) doi:10.1101/2022.02.17.480942.

101. Szlachcic, W. J., Ziojla, N., Kizewska, D. K., Kempa, M. & Borowiak, M. Endocrine Pancreas Development and Dysfunction Through the Lens of Single-Cell RNA-Sequencing. Front Cell Dev Biol 9, 629212 (2021).

102. Lim, C. T., Kola, B., Grossman, A. & Korbonits, M. The expression of ghrelin O-acyltransferase (GOAT) in human tissues. Endocr. J. 58, 707–710 (2011).

103. Dominguez Gutierrez, G., et al. Gene Signature of Proliferating Human Pancreatic α Cells. Endocrinology 159, 3177–3186 (2018).

104. Atla, G. et al. Genetic regulation of RNA splicing in human pancreatic islets. Genome Biol. 23, 196 (2022).

105. Mastrolia, V. et al. Loss of α2δ-1 Calcium Channel Subunit Function Increases the Susceptibility for Diabetes. Diabetes 66, 897–907 (2017).

106. Hrovatin, K., et al. Delineating mouse β-cell identity during lifetime and in diabetes with a single cell atlas. bioRxiv 2022.12.22.521557 (2022) doi:10.1101/2022.12.22.521557.

107. Schreiber, V. et al. Extensive NEUROG3 occupancy in the human pancreatic endocrine gene regulatory network. Mol Metab 53, 101313 (2021).

108. Salinno, C. et al. β-Cell Maturation and Identity in Health and Disease. Int. J. Mol. Sci. 20, (2019).

109. Salinno, C. et al. CD81 marks immature and dedifferentiated pancreatic β-cells. Mol Metab 49, 101188 (2021).

110. Wang, X. et al. Point mutations in the PDX1 transactivation domain impair human β-cell development and function. Mol Metab 24, 80–97 (2019).

111. Ramond, C. et al. Understanding human fetal pancreas development using subpopulation sorting, RNA sequencing and single-cell profiling. Development 145, (2018).

## References

[1] Gabriel Peyré, et al. Computational optimal transport: With applications to data science. Foundations and Trends in Machine Learning, 11(5-6):355–607, 2019.

[2] Geoffrey Schiebinger, et al. Optimal-transport analysis of single-cell gene expression identifies developmental trajectories in reprogramming. Cell, 176(4):928–943, 2019.

[3] Aden Forrow and Geoffrey Schiebinger. Lineageot is a unified framework for lineage tracing and trajectory inference. Nature communications, 12(1):1–10, 2021.

[4] Alexander Tong, et al. Trajectorynet: A dynamic optimal transport network for modeling cellular dynamics. In International conference on machine learning, pages 9526–9536. PMLR, 2020.

[5] Karren Dai Yang, et al. Predicting cell lineages using autoencoders and optimal transport. PLoS computational biology, 16(4):e1007828, 2020.

[6] Neha Prasad, et al. Optimal transport using gans for lineage tracing. arXiv preprint arXiv:2007.12098, 2020.

[7] Stephen Zhang, et al. Optimal transport analysis reveals trajectories in steady-state systems. PLoS computational biology, 17(12):e1009466, 2021.

[8] Mor Nitzan, et al. Gene expression cartography. Nature, 576(7785):132–137, 2019.

[9] Zixuan Cang and Qing Nie. Inferring spatial and signaling relationships between cells from single cell transcriptomic data. Nature communications, 11(1):1–13, 2020.

[10] Ron Zeira, et al. Alignment and integration of spatial transcriptomics data. Nature Methods, 19(5): 567–575, 2022.

[11] Pinar Demetci, et al. Scot: Single-cell multi-omics alignment with optimal transport. Journal of Computational Biology, 29(1):3–18, 2022.

[12] Kai Cao, et al. uniport: a unified computational framework for single-cell data integration with optimal transport. bioRxiv, 2022.

[13] William S Chen, et al. Uncovering axes of variation among single-cell cancer specimens. Nature methods, 17(3):302–310, 2020.

[14] Alexander Y Tong, et al. Diffusion earth mover’s distance and distribution embeddings. In International Conference on Machine Learning, pages 10336–10346. PMLR, 2021.

[15] Charlotte Bunne, et al. Learning single-cell perturbation responses using neural optimal transport. bioRxiv, 2021.

[16] Charlotte Bunne, et al. Supervised training of conditional monge maps. *arXiv preprint arXiv:2206.14262*, 2022.

[17] Gabriel Peyré, et al. Gromov-wasserstein averaging of kernel and distance matrices. In International Conference on Machine Learning, pages 2664–2672. PMLR, 2016.

[18] Adam Gayoso, et al. A python library for probabilistic analysis of single-cell omics data. Nature Biotechnology, 40(2):163–166, 2022.

[19] Marlon Stoeckius, et al. Simultaneous epitope and transcriptome measurement in single cells. Nature methods, 14(9):865–868, 2017.

[20] Sai Ma, et al. Chromatin potential identified by shared single-cell profiling of rna and chromatin. Cell, 183(4):1103–1116, 2020.

[21] Chenxu Zhu, et al. An ultra high-throughput method for single-cell joint analysis of open chromatin and transcriptome. Nature structural & molecular biology, 26(11):1063–1070, 2019.

[22] Song Chen, et al. High-throughput sequencing of the transcriptome and chromatin accessibility in the same cell. Nature biotechnology, 37(12):1452–1457, 2019.

[23] F Alexander Wolf, et al. Scanpy: large-scale single-cell gene expression data analysis. Genome biology, 19(1):1–5, 2018.

[24] Isaac Virshup, et al. The scverse project provides a computational ecosystem for single-cell omics data analysis. Nature Biotechnology, pages 1–3, 2023.

[25] Marco Cuturi, et al. Optimal transport tools (ott): A jax toolbox for all things wasserstein. arXiv preprint arXiv:2201.12324, 2022.

[26] Jason D Buenrostro, et al. Single-cell chromatin accessibility reveals principles of regulatory variation. Nature, 523(7561):486–490, 2015.

[27] Scott Deerwester, et al. Indexing by latent semantic analysis. Journal of the American society for information science, 41(6):391–407, 1990.

[28] Aude Genevay, et al. Sample complexity of sinkhorn divergences. In The 22nd international conference on artificial intelligence and statistics, pages 1574–1583. PMLR, 2019.

[29] Marco Cuturi. Sinkhorn distances: Lightspeed computation of optimal transport. Advances in neural information processing systems, 26, 2013.

[30] G Udny Yule. On the methods of measuring association between two attributes. Journal of the Royal Statistical Society, 75(6):579–652, 1912.

[31] Richard Sinkhorn. A relationship between arbitrary positive matrices and doubly stochastic matrices. The annals of mathematical statistics, 35(2):876–879, 1964.

[32] Matthias Liero, et al. Optimal entropy-transport problems and a new hellinger–kantorovich distance between positive measures. Inventiones mathematicae, 211(3):969–1117, 2018.

[33] Lenaic Chizat, et al. Scaling algorithms for unbalanced optimal transport problems. Mathematics of Computation, 87(314):2563–2609, 2018.

[34] Lenaic Chizat, et al. Unbalanced optimal transport: Dynamic and kantorovich formulations. Journal of Functional Analysis, 274(11):3090–3123, 2018.

[35] Adam Gayoso, et al. Joint probabilistic modeling of single-cell multi-omic data with totalvi. Nature methods, 18(3):272–282, 2021.

[36] Tal Ashuach, et al. Multivi: deep generative model for the integration of multi-modal data. bioRxiv, 2021.

[37] Mohammad Lotfollahi, et al. Multigrate: single-cell multi-omic data integration. bioRxiv, 2022.

[38] Junyue Cao, et al. Joint profiling of chromatin accessibility and gene expression in thousands of single cells. Science, 361(6409):1380–1385, 2018.

[39] Roy Frostig, et al. Compiling machine learning programs via high-level tracing. Systems for Machine Learning, 2018.

[40] Meyer Scetbon, et al. Low-rank sinkhorn factorization. In *International Conference on Machine Learning*, pages 9344–9354. PMLR, 2021.

[41] Aden Forrow, et al. Statistical optimal transport via factored couplings. In The 22nd International Conference on Artificial Intelligence and Statistics, pages 2454–2465. PMLR, 2019.

[42] Giovanni Palla, et al. Spatial components of molecular tissue biology. Nature Biotechnology, 40(3): 308–318, 2022.

[43] Cameron G Williams, et al. An introduction to spatial transcriptomics for biomedical research. Genome Medicine, 14(1):1–18, 2022.

[44] Bin Li, et al. Benchmarking spatial and single-cell transcriptomics integration methods for transcript distribution prediction and cell type deconvolution. Nature Methods, pages 1–9, 2022.

[45] Tommaso Biancalani, et al. Deep learning and alignment of spatially resolved single-cell transcriptomes with tangram. Nature methods, 18(11):1352–1362, 2021.

[46] Vitalii Kleshchevnikov, et al. Cell2location maps fine-grained cell types in spatial transcriptomics. Nature biotechnology, 40(5):661–671, 2022.

[47] Romain Lopez, et al. Destvi identifies continuums of cell types in spatial transcriptomics data. Nature biotechnology, pages 1–10, 2022.

[48] Alma Andersson, et al. Single-cell and spatial transcriptomics enables probabilistic inference of cell type topography. Communications biology, 3(1):1–8, 2020.

[49] Jeffrey R Moffitt, et al. Molecular, spatial, and functional single-cell profiling of the hypothalamic preoptic region. Science, 362(6416), November 2018.

[50] Chee-Huat Linus Eng, et al. Transcriptome-scale super-resolved imaging in tissues by RNA seqFISH. Nature, 568(7751):235–239, April 2019.

[51] Kok Hao Chen, et al. Spatially resolved, highly multiplexed rna profiling in single cells. Science, 348 (6233):aaa6090, 2015.

[52] Chee-Huat Linus Eng, et al. Transcriptome-scale super-resolved imaging in tissues by rna seqfish+. Nature, 568(7751):235–239, 2019.

[53] Giovanni Palla, et al. Squidpy: a scalable framework for spatial omics analysis. Nature methods, 19(2): 171–178, 2022.

[54] Samuel G Rodriques, et al. Slide-seq: A scalable technology for measuring genome-wide expression at high spatial resolution. Science, 363(6434):1463–1467, 2019.

[55] Robert R Stickels, et al. Highly sensitive spatial transcriptomics at near-cellular resolution with slide-seqv2. Nature biotechnology, 39(3):313–319, 2021.

[56] Yang Liu, et al. High-spatial-resolution multi-omics sequencing via deterministic barcoding in tissue. Cell, 183(6):1665–1681, 2020.

[57] Romain Lopez, et al. Deep generative modeling for single-cell transcriptomics. Nature methods, 15(12): 1053–1058, 2018.

[58] Titouan Vayer, et al. Fused gromov-wasserstein distance for structured objects. Algorithms, 13(9):212, 2020.

[59] Thibault Séjourné, et al. The unbalanced gromov wasserstein distance: Conic formulation and relaxation. Advances in Neural Information Processing Systems, 34:8766–8779, 2021.

[60] Vanessa M Peterson, et al. Multiplexed quantification of proteins and transcripts in single cells. Nature biotechnology, 35(10):936–939, 2017.

[61] Zhi-Jie Cao and Ge Gao. Multi-omics single-cell data integration and regulatory inference with graph-linked embedding. Nature Biotechnology, pages 1–9, 2022.

[62] Meyer Scetbon, et al. Linear-time gromov wasserstein distances using low rank couplings and costs. *arXiv preprint arXiv*:2106.01128, 2021.

[63] Jennifer E Rood, et al. Toward a common coordinate framework for the human body. Cell, 179(7): 1455–1467, 2019.

[64] Kristen R Maynard, et al. Transcriptome-scale spatial gene expression in the human dorsolateral prefrontal cortex. Nature neuroscience, 24(3):425–436, 2021.

[65] Andrew Jones, et al. Alignment of spatial genomics and histology data using deep gaussian processes. bioRxiv, 2022.

[66] Ao Chen, et al. Spatiotemporal transcriptomic atlas of mouse organogenesis using dna nanoball-patterned arrays. Cell, 185(10):1777–1792, 2022.

[67] Chang Liu, et al. Spatiotemporal mapping of gene expression landscapes and developmental trajectories during zebrafish embryogenesis. Developmental Cell, 57(10):1284–1298, 2022.

[68] Mingyue Wang, et al. High-resolution 3d spatiotemporal transcriptomic maps of developing drosophila embryos and larvae. Developmental Cell, 57(10):1271–1283, 2022.

[69] Keke Xia, et al. The single-cell stereo-seq reveals region-specific cell subtypes and transcriptome profiling in arabidopsis leaves. Developmental Cell, 57(10):1299–1310, 2022.

[70] Xiaoyu Wei, et al. Single-cell stereo-seq reveals induced progenitor cells involved in axolotl brain regeneration. Science, 377(6610):eabp9444, 2022.

[71] Hisham Mohammed, et al. Single-cell landscape of transcriptional heterogeneity and cell fate decisions during mouse early gastrulation. Cell reports, 20(5):1215–1228, 2017.

[72] Shangli Cheng, et al. Single-cell rna-seq reveals cellular heterogeneity of pluripotency transition and x chromosome dynamics during early mouse development. Cell reports, 26(10):2593–2607, 2019.

[73] Blanca Pijuan-Sala, et al. A single-cell molecular map of mouse gastrulation and early organogenesis. Nature, 566(7745):490–495, 2019.

[74] Chengxiang Qiu, et al. Systematic reconstruction of cellular trajectories across mouse embryogenesis. Nature genetics, 54(3):328–341, 2022.

[75] Junyue Cao, et al. The single-cell transcriptional landscape of mammalian organogenesis. Nature, 566 (7745):496–502, 2019.

[76] Tim Stuart, et al. Comprehensive integration of single-cell data. Cell, 177(7):1888–1902, 2019.

[77] Yuhan Hao, et al. Integrated analysis of multimodal single-cell data. Cell, 184(13):3573–3587, 2021.

[78] Leland McInnes, et al. Umap: Uniform manifold approximation and projection for dimension reduction. arXiv preprint arXiv:1802.03426, 2018.

[79] Haiyang Huang, et al. Towards a comprehensive evaluation of dimension reduction methods for transcriptomic data visualization. Communications biology, 5(1):719, 2022.

[80] Dmitry Kobak and George C Linderman. Initialization is critical for preserving global data structure in both t-sne and umap. Nature biotechnology, 39(2):156–157, 2021.

[81] Shamus M Cooley, et al. A novel metric reveals previously unrecognized distortion in dimensionality reduction of scrna-seq data. BioRxiv, page 689851, 2019.

[82] Laurens Van der Maaten and Geoffrey Hinton. Visualizing data using t-sne. Journal of machine learning research, 9(11), 2008.

[83] Tara Chari, et al. The specious art of single-cell genomics. BioRxiv, pages 2021–08, 2021.

[84] Cody N Heiser and Ken S Lau. A quantitative framework for evaluating single-cell data structure preservation by dimensionality reduction techniques. Cell reports, 31(5):107576, 2020.

[85] Daniel R Brison. Apoptosis in mammalian preimplantation embryos: regulation by survival factors. Human Fertility, 3(1):36–47, 2000.

[86] Daniel R Brison and Richard M Schultz. Apoptosis during mouse blastocyst formation: evidence for a role for survival factors including transforming growth factor *α*. Biology of reproduction, 56(5): 1088–1096, 1997.

[87] Katia Manova, et al. Apoptosis in mouse embryos: elevated levels in pregastrulae and in the distal anterior region of gastrulae of normal and mutant mice. Developmental dynamics: an official publication of the American Association of Anatomists, 213(3):293–308, 1998.

[88] Brian A Kilburn, et al. Rapid induction of apoptosis in gastrulating mouse embryos by ethanol and its prevention by hb-egf. Alcoholism: Clinical and Experimental Research, 30(1):127–134, 2006.

[89] Kristel Martínez-Lagunas, et al. In vivo detection of programmed cell death during mouse heart development. Cell Death & Differentiation, 27(4):1398–1414, 2020.

[90] Clemens Kiecker, et al. Molecular specification of germ layers in vertebrate embryos. Cellular and Molecular Life Sciences, 73:923–947, 2016.

[91] K Shyamala, et al. Neural crest: The fourth germ layer. Journal of oral and maxillofacial pathology: JOMFP, 19(2):221, 2015.

[92] Virtanen P, et al. Scipy 1.0: Fundamental algorithms for scientific computing in python. Nature Methods, 17(3):261–272, 2020.

[93] Christopher S McGinnis, et al. Doubletfinder: doublet detection in single-cell rna sequencing data using artificial nearest neighbors. Cell systems, 8(4):329–337, 2019.

[94] Tim Stuart, et al. Single-cell chromatin state analysis with signac. Nature methods, 18(11):1333–1341, 2021.

[95] Christoph Hafemeister and Rahul Satija. Normalization and variance stabilization of single-cell rna-seq data using regularized negative binomial regression. Genome biology, 20(1):1–15, 2019.

[96] Jonathan A Griffiths, et al. Detection and removal of barcode swapping in single-cell rna-seq data. Nature communications, 9(1):1–6, 2018.

[97] Laurence A Lemaire, et al. Bicaudal c1 promotes pancreatic neurog3+ endocrine progenitor differentiation and ductal morphogenesis. Development, 142(5):858–870, 2015.

[98] Aimée Bastidas-Ponce, et al. Comprehensive single cell mrna profiling reveals a detailed roadmap for pancreatic endocrinogenesis. Development, 146(12):dev173849, 2019.

[99] Itay Tirosh, et al. Dissecting the multicellular ecosystem of metastatic melanoma by single-cell rna-seq. Science, 352(6282):189–196, 2016.

[100] Jonas Ahnfelt-Rønne, et al. Preservation of proliferating pancreatic progenitor cells by delta-notch signaling in the embryonic chicken pancreas. BMC developmental biology, 7(1):1–13, 2007.

[101] Lauren E Byrnes, et al. Lineage dynamics of murine pancreatic development at single-cell resolution. Nature communications, 9(1):1–17, 2018.

[102] Jia Zhang, et al. The diabetes gene hhex maintains *δ*-cell differentiation and islet function. Genes & development, 28(8):829–834, 2014.

[103] Michaël Defferrard, et al. Pygsp: Graph signal processing in python. URL https://github.com/epfl-lts2/pygsp/.

[104] Laura D Martens, et al. Modeling fragment counts improves single-cell atac-seq analysis. bioRxiv, 2022.

[105] Matthew T Weirauch, et al. Determination and inference of eukaryotic transcription factor sequence specificity. Cell, 158(6):1431–1443, 2014.

[106] Carmen Bravo González-Blas, et al. Scenic+: single-cell multiomic inference of enhancers and gene regulatory networks. bioRxiv, pages 2022–08, 2022.

[107] Romain Lopez, et al. A joint model of unpaired data from scRNA-seq and spatial transcriptomics for imputing missing gene expression measurements. arXiv preprint arXiv:1905.02269, May 2019. eprint: 1905.02269.

[108] Daniel Zügner, et al. SEML: Slurm Experiment Management Library, 2022. URL https://github.com/TUM-DAML/seml.

[109] Martin Guilliams, et al. Spatial proteogenomics reveals distinct and evolutionarily conserved hepatic macrophage niches. Cell, 185(2):379–396.e38, January 2022.

[110] Marius Lange, et al. Cellrank for directed single-cell fate mapping. Nature methods, 19(2):159–170, 2022.

[111] Bernhard Reuter, et al. Generalized markov modeling of nonreversible molecular kinetics. The Journal of chemical physics, 150(17):174103, 2019.

[112] Bernhard Reuter, et al. Generalized markov state modeling method for nonequilibrium biomolecular dynamics: exemplified on amyloid *β* conformational dynamics driven by an oscillating electric field. Journal of Chemical Theory and Computation, 14(7):3579–3594, 2018.

