## Supplementary notes for "Mapping cells through time and space with moscot": moscot__supplementary_notes_.pdf

May 11, 2023

### Moscot Supplementary Notes

#### Contents

1 Fused Gromov-Wasserstein optimization 2

2 Fused Gromov-Wasserstein scalability 2

3 Low-rank Sinkhorn factorization yields linear time and memory complexity 3

4 Background information on pancreatic endocrinogenesis 5

5 Geodesic cost functions 5

6 Differential accessibility analysis 6

7 Motif activity analysis in pancreatic endocrinogenesis 6

### 1 Fused Gromov-Wasserstein optimization

Consider a generic FWG-type problem,

$$P^* \coloneqq \underset{P \in U(\boldsymbol{a},\boldsymbol{b})}{\operatorname{argmin}} \; \alpha \sum_{ijkl} L\left(C^X_{ij}, C^Y_{kl}\right) P_{ik} P_{jl} + (1-\alpha) \sum_{ik} C_{ik} P_{ik} - \epsilon H(P) \,, \tag{1}$$

for within-space cost matrices $C^X \in \mathbb{R}^{N \times N}$ and $C^Y \in \mathbb{R}^{M \times M}$, across-space cost matrix $C \in \mathbb{R}^{N \times M}$, distance metric $L$, weight parameter $\alpha \in [0, 1]$ and entropic regularization $H(P)$ at strength $\epsilon$. To write this optimization problem in shorter form, we define the 4-tensor[1]

$$\mathcal{T}(C^X, C^Y)_{ijkl} \coloneqq L\left(C^X_{ik}, C^Y_{jl}\right) \,, \tag{2}$$

which allows us to rewrite Equation (1) as,

$$P^* = \underset{P \in U(\boldsymbol{a},\boldsymbol{b})}{\operatorname{argmin}} \; \alpha \left\langle \mathcal{T}(C^X, C^Y) \otimes P, P \right\rangle + (1-\alpha) \left\langle C, P \right\rangle - \epsilon H(P) \,, \tag{3}$$

where tensor multiplication is defined as

$$(\mathcal{T} \otimes P)_{ij} \coloneqq \sum_{kl} \mathcal{T}_{ijkl} P_{kl} \,. \tag{4}$$

**Reduce to Sinkhorn iterations.** We next derive an algorithm in terms of Sinkhorn iterations; the basic idea is to use projected gradient descent with update rule[1]

$$P^{(l+1)} = \operatorname{Proj}^{\mathrm{KL}}_{U(\boldsymbol{a},\boldsymbol{b})}\left(P^{(l)} \odot e^{-\tau \nabla J(P)|_{P^{(l)}}}\right) , \tag{5}$$

where $\operatorname{Proj}^{\mathrm{KL}}_{U(\boldsymbol{a},\boldsymbol{b})}(\tilde{P}) = \operatorname{argmin}_{P \in U(\boldsymbol{a},\boldsymbol{b})} \sum_{ij} P_{ij} \log\left(P_{ij}/\tilde{P}_{ij}\right)$ is a KL projection operator, $\tau$ is a step size, $J$ is the FGW objective function defined in Equation (3) and $\odot$ denotes element-wise multiplication. We may rewrite the objective function gradient as

$$\nabla J = (1-\alpha) C + \alpha \mathcal{T}\left(C^X, C^Y\right) \otimes P \,. \tag{6}$$

Further, the KL projection can be solved via an OT problem[2],

$$\operatorname{Proj}^{\mathrm{KL}}_{U(\boldsymbol{a},\boldsymbol{b})}(\tilde{P}) = \underset{P \in U(\boldsymbol{a},\boldsymbol{b})}{\operatorname{argmin}} \left\langle -\epsilon \log \tilde{P}, P \right\rangle - \epsilon H(P) \,. \tag{7}$$

Using Equation (6), Equation (7) and setting $\tau = 1/\epsilon$, we can re-write the update rule of Equation (5) as

$$P^{(l+1)} = \underset{P \in U(\boldsymbol{a},\boldsymbol{b})}{\operatorname{argmin}} \left\langle (1-\alpha) C + \alpha \mathcal{T}\left(C^X, C^Y\right) \otimes P^{(l)}, P \right\rangle - \epsilon H(P) \,, \tag{8}$$

which is the entropically regularized W-type OT problem (Methods). We update the cost matrix at each outer iteration and solve the resulting W-type OT problem using the Sinkhorn algorithm[1,3,4].

### 2 Fused Gromov-Wasserstein scalability

Throughout this section, we assume equal cell/sample numbers across both datasets for simplicity, i.e., $N = M$. Further, we suppose entropic regularization is applied to the FGW-type OT problem[1,5] and the mirror descent algorithm (Section 1) is used for optimization.

**Cubic time complexity for separable loss function $L$.** The major computational bottleneck in FGW[1,5] optimization is the update of the tensor product of Equation (4) required at each update of Equation (8), which is quartic in cell number for general loss function $L$ (ref.[1] and Section 1). However, this can be improved if $L$ is given by a separable loss function. For simplicity, suppose $L$ is given by a squared $l_2$ loss function, this leads to

$$\mathcal{T}(C^X, C^Y) \otimes P^{(l)} = \underbrace{(C^X)^2 \boldsymbol{a}\, \mathbf{1}_M^\top + \mathbf{1}_N\, \boldsymbol{b}^\top (C^Y)^2}_{\text{constant}} - 2C^X P^{(l)} C^{Y^\top}, \tag{9}$$

which may be evaluated in time $\mathcal{O}(N^3)$ and memory $\mathcal{O}(N^2)$. This result holds beyond the squared $l_2$ norm for a class of separable loss functions, including the KL divergence[1].

**Quadratic time complexity for low-rank $C^X$ and $C^Y$.** For the second, non-constant term, further suppose both $C^X$ and $C^Y$ result from the application of a squared $l_2$ distance metric such that they admit low-rank factorizations,

$$C^X = A^X B^{X^\top} \text{ for } A^X, B^X \in \mathbb{R}^{N \times (D_x+2)}, \tag{10}$$

$$C^Y = A^Y B^{Y^\top} \text{ for } A^Y, B^Y \in \mathbb{R}^{M \times (D_y+2)}, \tag{11}$$

for within-space dimensions $D_x, D_y$ (ref.[6]). Such factorizations are obtained by specifying $A^X := [\boldsymbol{p}, \mathbf{1}_N, -2X]$ and $B^X := [\mathbf{1}_N, \boldsymbol{p}, X]$ for $\boldsymbol{p} := [||\boldsymbol{x}_1||_2^2, ..., ||\boldsymbol{x}_N||_2^2]^\top$, and analogously for $A^Y$ and $B^Y$ (Section 3 and ref.[6]). Thus, the non-constant part of Equation (9) may be written as

$$C^X P^{(l)} C^{Y^\top} = A^X B^{X^\top} P^{(l)} B^Y A^{Y^\top}. \tag{12}$$

This can be computed in time $\mathcal{O}(N^2(D_x + D_y))$, i.e., quadratic rather than cubic in the input size[7]. Note that all current `moscot` models use squared euclidean loss functions for all of $L$, $C^X$, and $C^Y$ and thus enjoy quadratic time complexity without any approximations. This represents a remarkable speedup compared to previous FGW-based[1,5] models in single-cell genomics[4,8] without any accuracy sacrifice.

For future `moscot` models that might require non-euclidean distance metrics, approximate algorithms exist to compute low-rank factorizations of $C^X$ and $C^Y$, which scale linearly in sample number[6,9,10]. Thus the overall algorithm time complexity remains quadratic.

**Linear time complexity for low-rank $P$.** The quadratic time and memory complexities become prohibitively expensive for atlas-scale datasets with hundreds of thousands of samples per dataset. Thus, we additionally assume low-rank structure in the coupling matrix $P$. Scetbon et al.[7] recently extended their low-rank Sinkhorn factorization (Section 3) to the FGW-setting[1,5], unlocking linear time and memory-complexity. Low-rank FGW solvers are implemented in OTT[11] and available to all FGW-based `moscot` models, including `moscot.space.mapping`, `moscot.space.alignment` and `moscot.spatiotemporal`.

### 3 Low-rank Sinkhorn factorization yields linear time and memory complexity

While the engineering improvements introduced in the Methods section allow the application to large datasets through GPU acceleration with linear or quadratic memory complexity for W-or GW-type problems, respectively, they still suffer from quadratic time complexity. Various authors have suggested approximations to the Sinkhorn iterations that yield linear time complexity to overcome this limitation. Altschuler et al.[12] suggest computing a low-rank approximation to the kernel matrix

$K$ using the Nystrom method[13]; their approach remains limited to squared euclidean cost functions $c$, is non-differentiable and only works for large regularization strength $\epsilon$ where inner iterations remain positive.

Forrow et al.[14] suggest a different route that imposes low-rank constraints on the feasible set of couplings $U(\boldsymbol{a}, \boldsymbol{b})$ rather than on the kernel matrix $K$. Their approach leads to an elegant solution via a barycenter problem; however, it remains limited to squared euclidean cost functions $c$. Scetbon et al.[6] generalize this approach to arbitrary cost functions $c$; their proposed solution is differentiable and applicable for a wide range of $\epsilon$ values, including no entropic regularization ($\epsilon = 0$). This approach is implemented in OTT and available through `moscot`; we refer to it as *low-rank Sinkhorn*. It has meanwhile been extended from W-type to GW-type problems[7] which is also implemented in OTT[11] and available to FGW-based[1,5] `moscot` models including `moscot.space.mapping`, `moscot.space.alignment`or `moscot.spatiotemporal` (Section 2).

For the low-rank Sinkhorn approach, following Scetbon et al.[6], define the nonnegative rank of a coupling matrix $P \in \mathbb{R}_+^{N \times M}$ to be

$$\mathrm{rk}_+(P) := \min \left\{ q \,\middle|\, P = \sum_{i=1}^{q} R_i \,, \mathrm{rk}(R_i) = 1,\ R_i \in \mathbb{R}_+^{N \times M} \right\} , \tag{13}$$

for rank rk. For $r \geq 1$, we make use of this to define the set of rank-$r$ couplings via

$$U(\boldsymbol{a}, \boldsymbol{b}, r) := \{ P \in U(\boldsymbol{a}, \boldsymbol{b}) \,|\, \mathrm{rk}_+(P) \leq r \} , \tag{14}$$

where $U(\boldsymbol{a}, \boldsymbol{b})$ is the set of feasible couplings. The rank-constrained feasible set $U(\boldsymbol{a}, \boldsymbol{b}, r)$ allows us to formulate the low-rank OT problem via

$$P^* := \operatorname*{argmin}_{P \in U(\boldsymbol{a}, \boldsymbol{b}, r)} \langle P, C \rangle - \epsilon H(P) \,. \tag{15}$$

An explicit characterisation of couplings $P$ in $U(\boldsymbol{a}, \boldsymbol{b}, r)$ is given by

$$P = Q \,\mathrm{diag}\,(1/\boldsymbol{g})\, R^\top \text{ for } \boldsymbol{g} \in \Delta_r^*,\ Q \in U(\boldsymbol{a}, \boldsymbol{g}),\ R \in U(\boldsymbol{b}, \boldsymbol{g}) , \tag{16}$$

where $\Delta_r^*$ denotes the $r$-simplex with strictly positive elements. Using this factorization, Scetbon et al.[7] derive a mirror descent optimization scheme for the low-rank OT problem of Equation (15); the time- and memory bottleneck in this algorithm is given by matrix-matrix multiplications of the form $CR$ and $C^\top Q$ for $Q \in \mathbb{R}^{N \times r}$ and $R \in \mathbb{R}^{M \times r}$. Thus, without any assumptions on the cost matrix $C$, the low-rank approach remains at memory complexity $\mathcal{O}(MN)$ and time complexity $\mathcal{O}(NMr)$.

To improve upon this complexity, assume that $C$ itself admits a low-rank factorization (Section 2), given by

$$C = AB^\top \text{ for } A \in \mathbb{R}^{N \times D},\ B \in \mathbb{R}^{M \times D} , \tag{17}$$

such that matrix-matrix multiplications $CR = A(B^\top R)$ and $C^\top Q = B(A^\top Q)$ can be evaluated in memory $\mathcal{O}\left((D + r)(M + N) + Dr\right)$ and time $\mathcal{O}\left(rD(N + M)\right)$, i.e. both linear in the total cell number $N + M$. In particular, such a factorization can be obtained if the cost results from applying a squared euclidean cost function, i.e. $C = c(X, Y) = ||X - Y||_2^2$. In such a case, $C$ may be written as

$$C = \boldsymbol{p}\mathbf{1}_M^\top + \mathbf{1}_N \boldsymbol{q}^\top - 2XY^\top , \tag{18}$$

for $\boldsymbol{p} := [||\boldsymbol{x}_1||_2^2, ..., ||\boldsymbol{x}_N||_2^2]^\top$ and $\boldsymbol{q} := [||\boldsymbol{y}_1||_2^2, ..., ||\boldsymbol{y}_M||_2^2]^\top$. The desired factorization is obtained by defining $A := [\boldsymbol{p}, \mathbf{1}_N, -2X] \in \mathbb{R}^{N \times (N_l + 2)}$, $B := [\mathbf{1}_M, \boldsymbol{q}, Y] \in \mathbb{R}^{M \times (N_l + 2)}$ for cells $\boldsymbol{x}_i$ and $\boldsymbol{y}_j$ embedded in some latent space of dimension $N_l$. In general, low-rank factorizations of cost matrices $C$ can be computed in linear time using randomized algorithms as long as the cost function $c$ is given by a proper distance metric[6,9,10].

### 4 Background information on pancreatic endocrinogenesis

The process of murine pancreatic endocrinogenesis can be divided into two stages. The primary stage is dominated by the creation of multipotent progenitor cells (MPCs), and mainly takes place between embryonic day (E) 9.5 and E12.5[15–17]. During the secondary stage the cells develop into three major lineages, namely Acinar cells, Ductal cells and the endocrine cells. During pancreatic lineage formation, first MPCs give rise to tip and trunk domains. The tip domain is differentiated into acinar cells and the trunk domain contains bipotent progenitors that give rise to ductal cells or endocrine progenitors. The later one is marked by the transient expression of the transcription factor Ngn3 and produce different hormone-producing cell including alpha cells (expressing glucagon), beta cells (insulin), epsilon cells (ghrelin), and delta cells (somatostatin) in a process so called endocrinogenesis.

### 5 Geodesic cost functions

The quality of a mapping obtained from an OT problem largely depends on the chosen cost function. To give an example, for time-series scRNA-seq data, we need to quantify the cost associated with transporting cells from $t_1$ to $t_2$. This amounts to defining a biologically meaningful distance metric among cells in gene expression space. Most OT applications to single-cell genomics use euclidean distances in latent spaces, such as PCA or scVI[18] latent space. However, euclidean distances might fail to capture the subtleties of cellular state changes during complex biological processes such as development, regeneration, or cancer. In related fields studying cellular manifolds and trajectories, researchers have successfully employed graph-based distance metrics to avoid the pitfalls of euclidean distances to capture phenotypic manifolds. For example, UMAP[19], t-SNE[20] and diffusion maps[21] use graph-based distance metrics to derive low-dimensional cellular representations, and DPT[22], Wanderlust[23] and Palantir[24] use graph-based distance metrics to estimate cellular trajectories[25].

Building on the success of these methods, Huguet et al.[26], Solomon et al.[27] suggested to use graph-based distance metrics to approximate geodesic distance in OT cost functions. Drawing on relationships between heat transfer on graphs and geodesic distance[28], they suggest to use a heat kernel $\mathcal{H}_t$ as drop in replacement for the usual Gibbs kernel (Methods), i.e.

$$K(x, y) = \mathcal{H}_t(x, y)\,, \tag{19}$$

for cells $x$ and $y$, and heat diffusion parameter $t$. Considering $N$ cells at $t_1$ and $M$ cells at $t_2$, they compute a joint single-cell k-nearest neighbor (kNN) graph $\mathcal{G}$ with associated graph laplacian matrix $L \in \mathbb{R}^{(M+N)\times(M+N)}$. The discrete heat operator $H_t$ on $\mathcal{G}$ may then be represented as $H_t = \exp(-tL)$[26]. This may be evaluated using the eigendecomposition of $L$, which exists and is real as $L$ is a real symmetric matrix. However, computing a full eigendecomposition of $L$ has cubic time complexity in terms of the total cell number $N + M$. Realizing that we do not actually require access to the full $H_t$ matrix, but only to matrix vector products within Sinkhorn iterations (Methods), Huguet et al.[26] suggest to approximate these matrix vector products using Chebyshev polynomials[27,29]. Taking advantage of the sparsity of kNN graphs, such an approach reduces the time complexity of Sinkhorn iterations to be linear in cell number $(N + M)$. The resulting *geodesic Sinkhorn* algorithm scales to large datasets and outperformed competing, euclidean-distance based OT-approaches, in a number of benchmarks, including interpolation of held-out time points in time-series scRNA-seq studies[26].

Geodesic distances currently have to be precomputed, but we plan to include more efficient computations into moscot in the future.

### 6 Differential accessibility analysis

To study the ATAC profile of the respective progenitor cells, we performed a differentially accessible peak analysis of epsilon progenitors to arrive at marker peaks using Signac (Methods). The most highly ranked peak overlaps with the coding regions of *Lrrtm3* (Fig. 5l, Supplementary Fig. 29a) and *Ctnna3* (Supplementary Fig. 29b), genes which the peak also shares a high peak-gene correlation with. The peak is accessible by the epsilon progenitor population, the Ngn3$^{\text{high-1}}$ population and the Fev+ delta populations, confirming the similarity of fate potentials of these cell states. We made similar observations for the most notable peaks in the Fev+ delta-0 and the Fev+ delta-1 population (Fig. 5m, Supplementary Fig. 29c-d), respectively. The marker peak of Fev+ delta-0 is most highly correlated with the expression of *Cxxc4*, a gene previously reported as a key transcription factor (TF) for the delta lineage[30]. The marker peak of Fev+ delta-1 is associated with *Cacnb2*, a gene responsible for insulin secretion in beta cells, and which was previously reported as a target gene of *Neurog3* in humans[31]. These observations support our lineage hypotheses and conjecture of high plasticity of Fev+ delta cells formulated above.

### 7 Motif activity analysis in pancreatic endocrinogenesis

#### 7.1 Motif activity in delta cells

For delta cells, the most remarkable marker TF identified with moscot is *Hhex* (Pearson correlation coefficient 0.44) but no motif associated with *Hhex* was highly expressed (we hypothesize this to be the case as there is no directly measured motif in the cisBP data base (Methods) and the inferred ones suffer from low quality). The second most remarkable is *Arg1* (0.25), followed by *Isl1* (0.22). The most remarkable motif according to the Wilcoxon test has cisBP identifier M09209_2.00, with a log-fold change of 4.08. It is directly measured, and associated with *Isl1*. Hence, we report this motif as the marker motif for delta cells.

#### 7.2 Motif activity in epsilon cells

For epsilon cells, motifs related to the *Foxa* family were particularly differentiably accessible, while *Arg1* was identified as a marker TF by moscot, yet, cisBP does not provide an associated motif for this TF. Thus, we identified *Tead1* (0.15) as key TF because the directly measured motif *M09438_2.00* is also substantially active. *Tead1*, a Hippo signalling effector, is known for its role in early fate decisions of endocrine progenitors, hence it is highly expressed in Ductal cells and Ngn3 low cells[32]. As the differential motif activity test was performed on the whole dataset, the motif score is negative for the cell types shown in the plots. Yet, the expression of *Tead1* decreases substantially along the Ngn3 high,0 / Fev+ lineage, while it remains high in the Ngn3 high,1 / epsilon progenitor lineage.

#### 7.3 Motif activity in alpha cells

Concerning alpha cells, motifs of the *Pax2*/*Pax6* family were particularly active but the expression of associated genes was not remarkable in the alpha/alpha progenitor population. Thus, we chose the motif *M03318_2.00*(2.68) as it is only associated with the TF *Pou3f4*, which is also among the marker TFs identified with moscot (0.26).

#### 7.4 Motif activity in beta cells

For beta cells, *Mafb* is the most highly correlated transcription factor identified with moscot (0.59), while the most active motif according to the Wilcoxon test is *M08835_2.00*, directly measured with *Mafg*, and moreover associated with [*Mafk*, *Mafa*, *Mafg*, *Maf*, *Mafb*]. As *Mafg* is also recovered as a marker feature with moscot (0.29), we identify *Mafg* and the aforementioned motif with cisBP identifier *M08835_2.00* as marker features for beta cells.

#### References

[1] Gabriel Peyré, et al. Gromov-wasserstein averaging of kernel and distance matrices. In *International Conference on Machine Learning*, pages 2664–2672. PMLR, 2016.

[2] Jean-David Benamou, et al. Iterative bregman projections for regularized transportation problems. *SIAM Journal on Scientific Computing*, 37(2):A1111–A1138, 2015.

[3] Marco Cuturi. Sinkhorn distances: Lightspeed computation of optimal transport. *Advances in neural information processing systems*, 26, 2013.

[4] Mor Nitzan, et al. Gene expression cartography. *Nature*, 576(7785):132–137, 2019.

[5] Titouan Vayer, et al. Fused gromov-wasserstein distance for structured objects. *Algorithms*, 13(9):212, 2020.

[6] Meyer Scetbon, et al. Low-rank sinkhorn factorization. In *International Conference on Machine Learning*, pages 9344–9354. PMLR, 2021.

[7] Meyer Scetbon, et al. Linear-time gromov wasserstein distances using low rank couplings and costs. *arXiv preprint arXiv:2106.01128*, 2021.

[8] Ron Zeira, et al. Alignment and integration of spatial transcriptomics data. *Nature Methods*, 19(5): 567–575, 2022.

[9] Ainesh Bakshi and David Woodruff. Sublinear time low-rank approximation of distance matrices. *Advances in Neural Information Processing Systems*, 31, 2018.

[10] Pitor Indyk, et al. Sample-optimal low-rank approximation of distance matrices. In *Conference on Learning Theory*, pages 1723–1751. PMLR, 2019.

[11] Marco Cuturi, et al. Optimal transport tools (ott): A jax toolbox for all things wasserstein. *arXiv preprint arXiv:2201.12324*, 2022.

[12] Jason Altschuler, et al. Massively scalable sinkhorn distances via the nyström method. *Advances in neural information processing systems*, 32, 2019.

[13] Christopher Williams and Matthias Seeger. Using the nyström method to speed up kernel machines. *Advances in neural information processing systems*, 13, 2000.

[14] Aden Forrow, et al. Statistical optimal transport via factored couplings. In *The 22nd International Conference on Artificial Intelligence and Statistics*, pages 2454–2465. PMLR, 2019.

[15] George K Gittes. Developmental biology of the pancreas: a comprehensive review. *Developmental biology*, 326(1):4–35, 2009.

[16] Eckhard Lammert, et al. Role of endothelial cells in early pancreas and liver development. *Mechanisms of development*, 120(1):59–64, 2003.

[17] Hjalte List Larsen and Anne Grapin-Botton. The molecular and morphogenetic basis of pancreas organogenesis. In *Seminars in cell & developmental biology*, volume 66, pages 51–68. Elsevier, 2017.

[18] Romain Lopez, et al. Deep generative modeling for single-cell transcriptomics. *Nature methods*, 15(12): 1053–1058, 2018.

[19] Leland McInnes, et al. Umap: Uniform manifold approximation and projection for dimension reduction. *arXiv preprint arXiv:1802.03426*, 2018.

[20] Laurens Van der Maaten and Geoffrey Hinton. Visualizing data using t-sne. *Journal of machine learning research*, 9(11), 2008.

[21] Laleh Haghverdi, et al. Diffusion maps for high-dimensional single-cell analysis of differentiation data. *Bioinformatics*, 31(18):2989–2998, 2015.

[22] Laleh Haghverdi, et al. Diffusion pseudotime robustly reconstructs lineage branching. *Nature methods*, 13(10):845–848, 2016.

[23] Sean C Bendall, et al. Single-cell trajectory detection uncovers progression and regulatory coordination in human b cell development. *Cell*, 157(3):714–725, 2014.

[24] Manu Setty, et al. Characterization of cell fate probabilities in single-cell data with palantir. *Nature biotechnology*, 37(4):451–460, 2019.

[25] Sophie Tritschler, et al. Concepts and limitations for learning developmental trajectories from single cell genomics. *Development*, 146(12):dev170506, 2019.

[26] Guillaume Huguet, et al. Geodesic sinkhorn: optimal transport for high-dimensional datasets. *arXiv preprint arXiv:2211.00805*, 2022.

[27] Justin Solomon, et al. Convolutional wasserstein distances: Efficient optimal transportation on geometric domains. *ACM Transactions on Graphics (ToG)*, 34(4):1–11, 2015.

[28] Sathamangalam R Srinivasa Varadhan. On the behavior of the fundamental solution of the heat equation with variable coefficients. *Communications on Pure and Applied Mathematics*, 20(2):431–455, 1967.

[29] Sibylle Marcotte, et al. Fast multiscale diffusion on graphs. In *ICASSP 2022-2022 IEEE International Conference on Acoustics, Speech and Signal Processing (ICASSP)*, pages 5627–5631. IEEE, 2022.

[30] Eliza Duvall, et al. Single-cell transcriptome and accessible chromatin dynamics during endocrine pancreas development. *Proceedings of the National Academy of Sciences*, 119(26):e2201267119, 2022.

[31] Valérie Schreiber, et al. Extensive neurog3 occupancy in the human pancreatic endocrine gene regulatory network. *Molecular metabolism*, 53:101313, 2021.

[32] Inês Cebola, et al. Tead and yap regulate the enhancer network of human embryonic pancreatic progenitors. *Nature cell biology*, 17(5):615–626, 2015.
