## Supplementary figures for "Mapping cells through time and space with moscot"

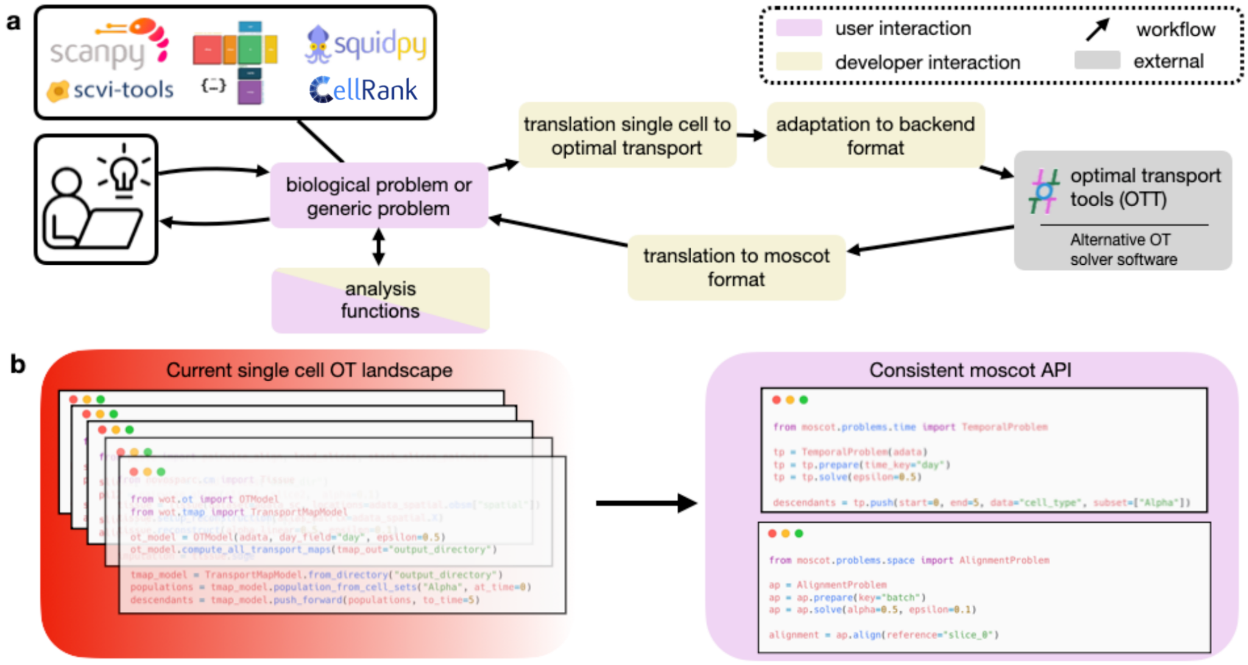


**Suppl. Fig. 1 | Moscot provides unified access to various problems in single-cell genomics**

**a.** The moscot workflow. The user interacts with a biological problem, which moscot translates into an OT problem and solves in the backend using Optimal Transport Tools (OTT[22]). Moscot then presents the solution to the user, who can further analyze it using downstream analysis functions. **b.** Comparison of code required to solve different biological problems in the current OT landscape and in moscot. Through its consistent API, moscot is user-friendly and easy to extend.

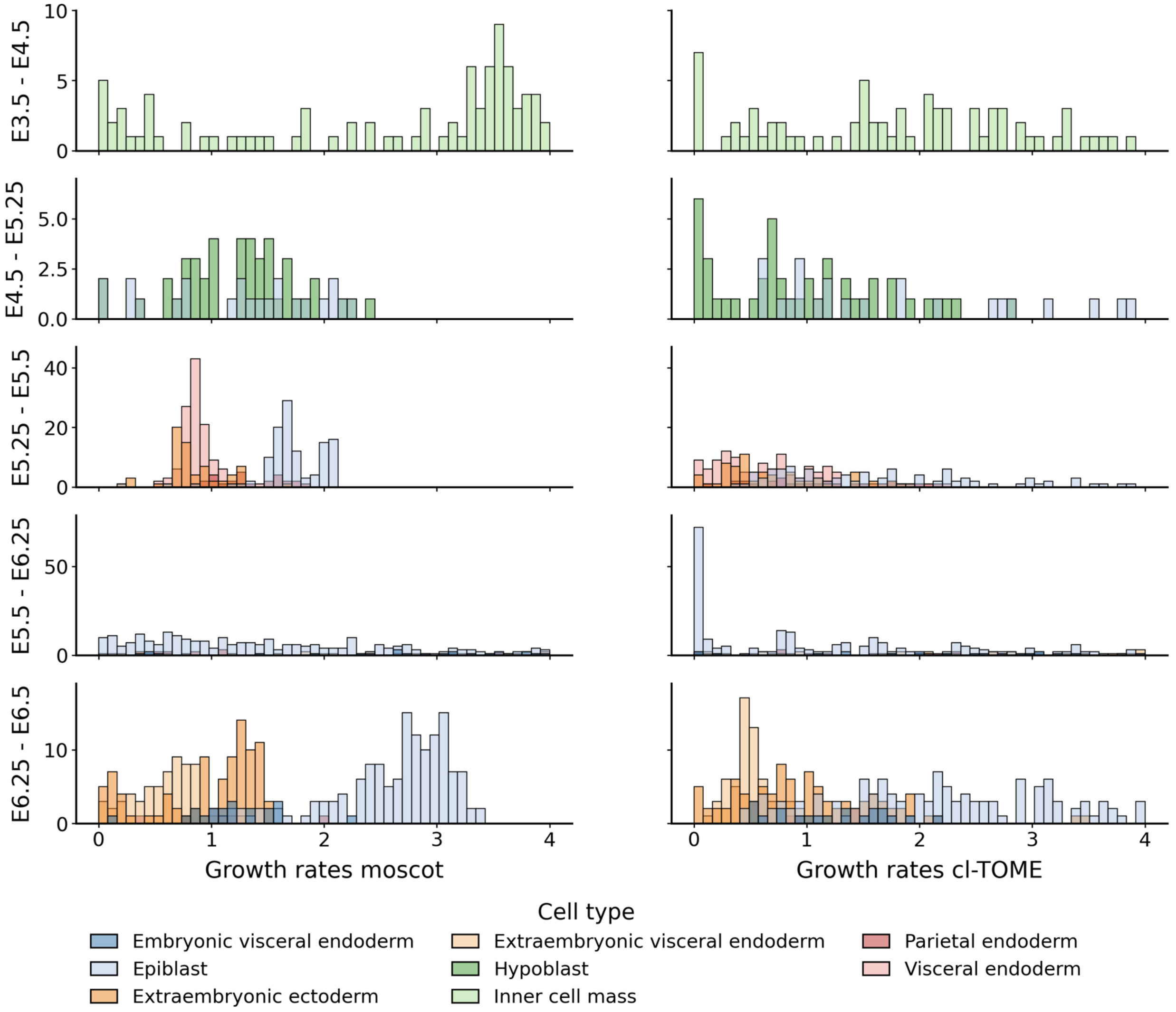


**Suppl. Fig. 2 | Comparison of predicted pre-gastrulation growth rates for moscot and cl-TOME**

Histograms over moscot and cl-TOME predicted growth rates, grouped by cell type (Methods). Growth rates correspond to the predicted amount of descendants for each cell for the transition from the earlier to the later time point. Growth rates bigger or smaller than one correspond to cell proliferation or growth, respectively.

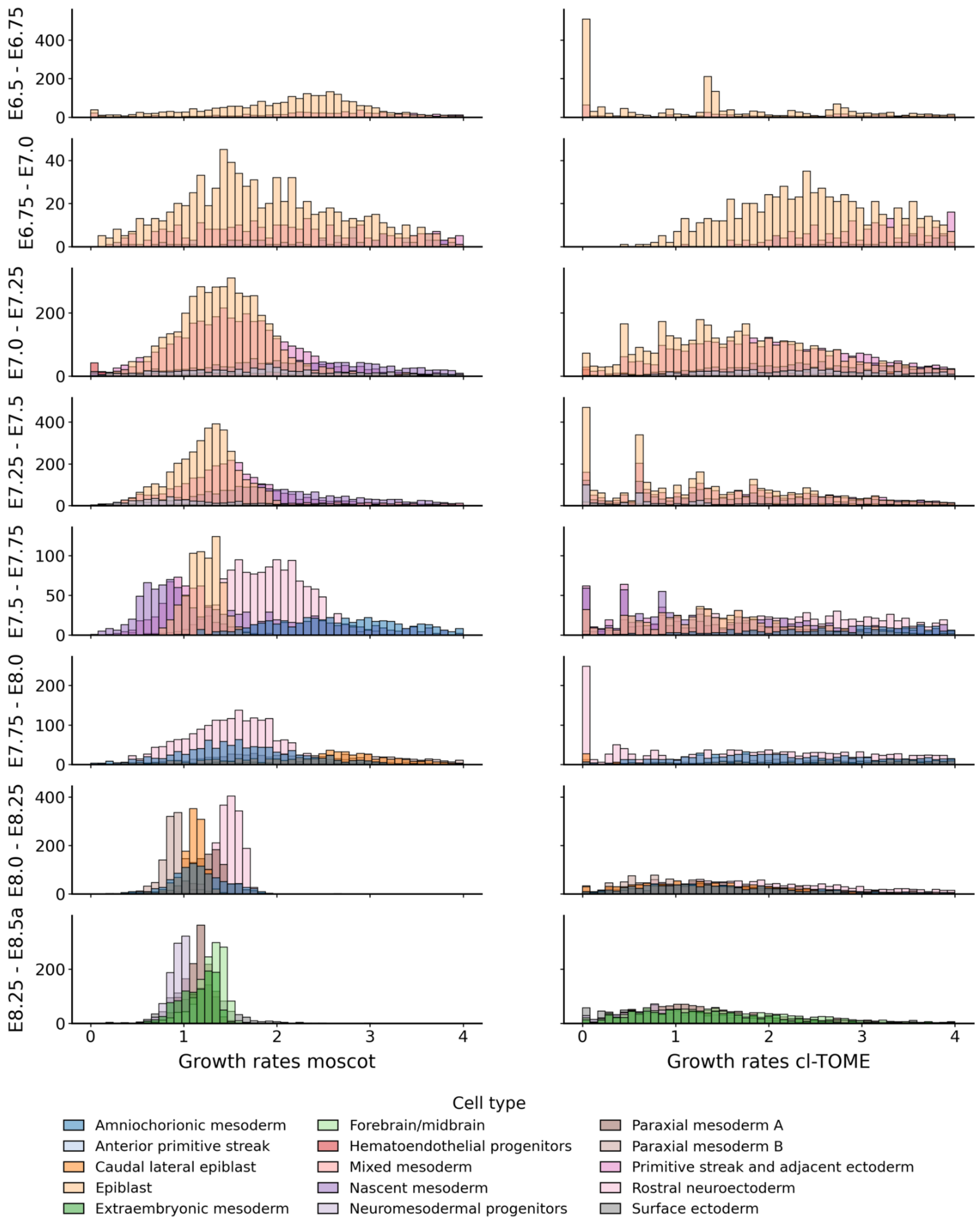


**Suppl. Fig. 3 | Comparison of predicted gastrulation growth rates moscot and cl-TOME**

See the description of Supplementary Fig. 2.

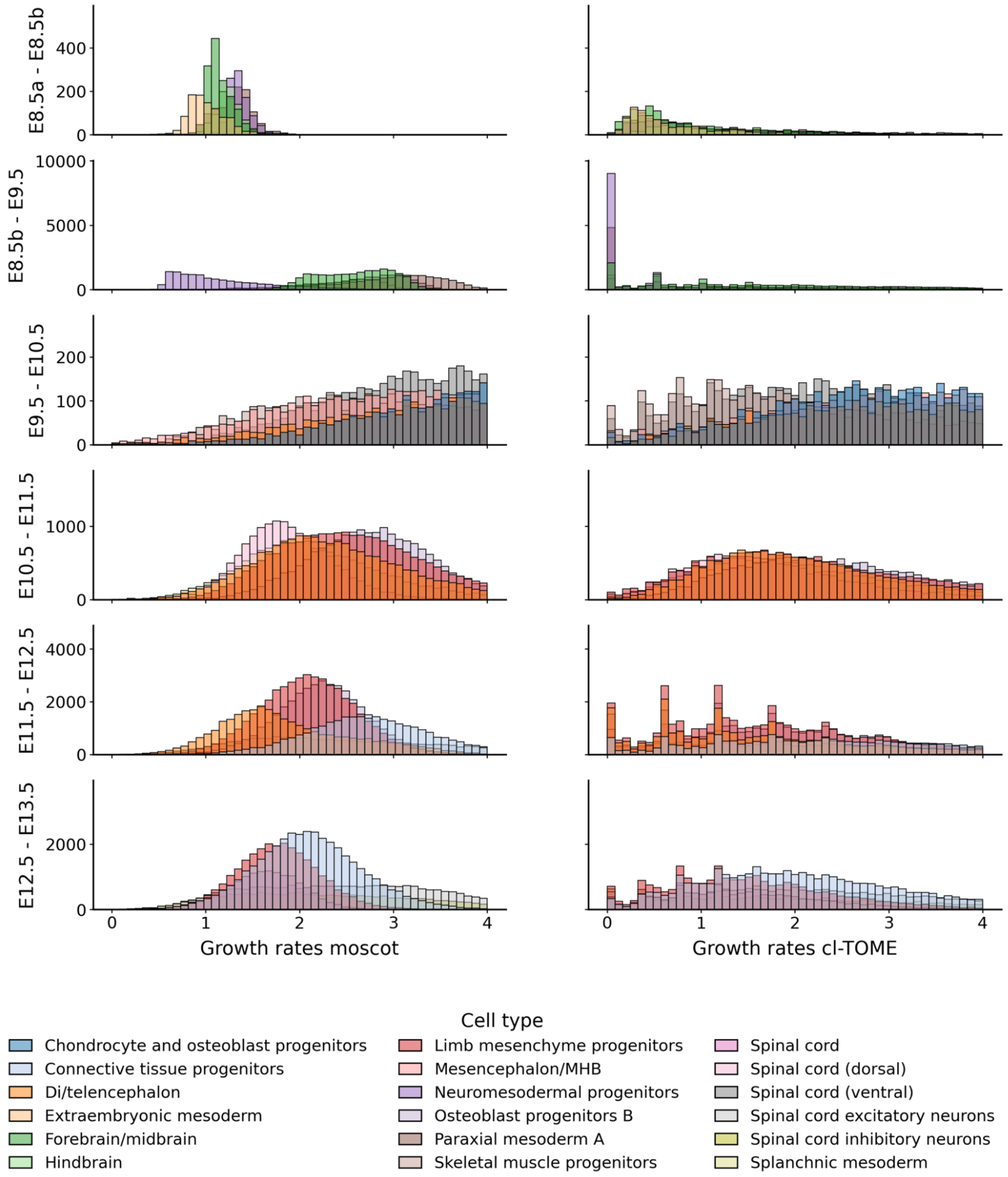


**Suppl. Fig. 4 | Comparison of predicted organogenesis growth rates moscot and cl-TOME**

See the description of Supplementary Fig. 2. Additionally, at E8.5a/b, no experimental time passes but the experimental protocol switches from 10x genomics to sci-RNA-seq3[16].

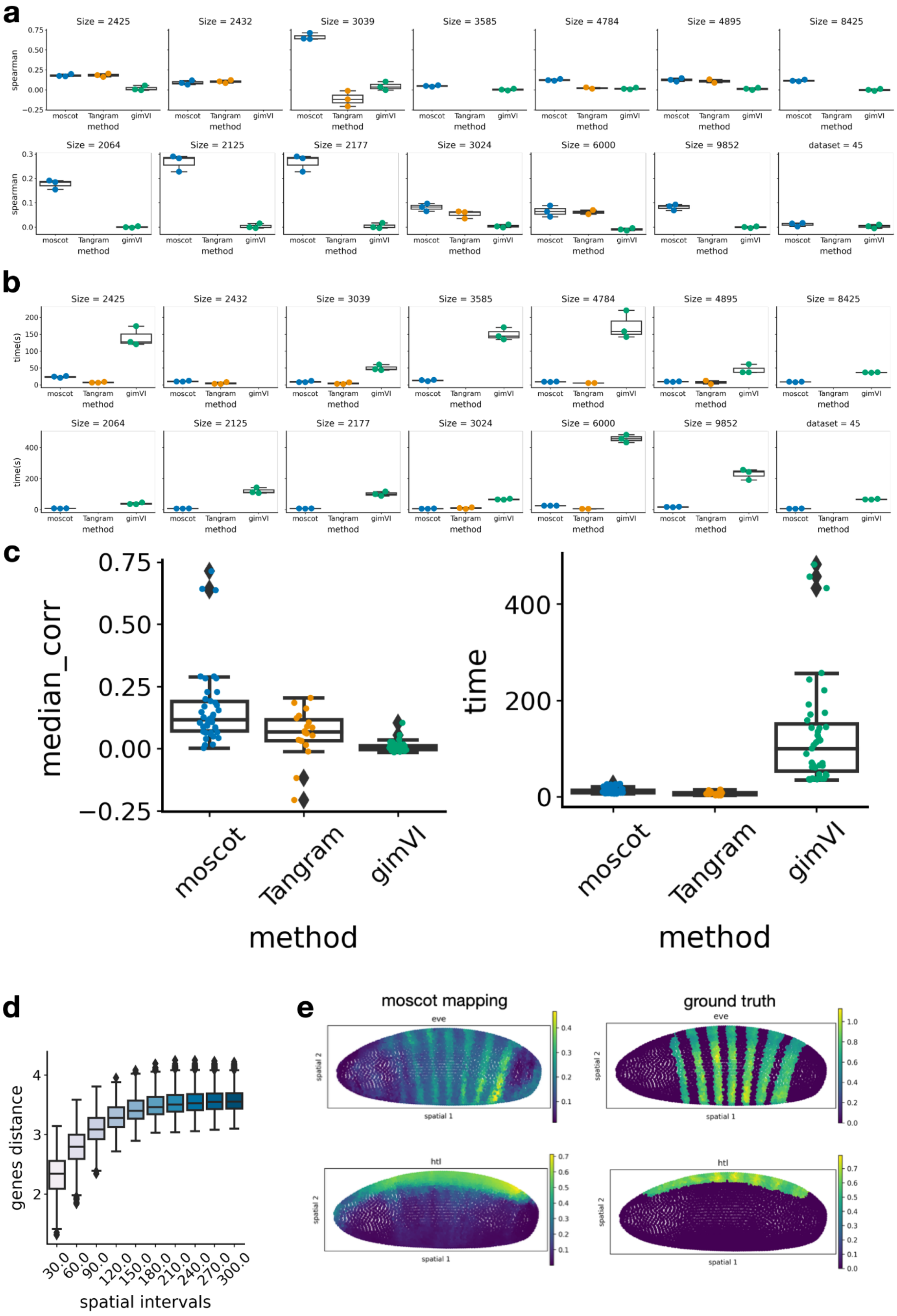

a
b
c
d
e
Size = 2425
Size = 2432
Size = 3039
Size = 3585
Size = 4784
Size = 4895
Size = 8425
Size = 2064
Size = 2125
Size = 2177
Size = 3024
Size = 6000
Size = 9852
dataset = 45
spearman
time(s)
moscot
Tangram
gimVI
method
median_corr
time
genes distance
spatial intervals
moscot mapping
ground truth
eve
htl
spatial 1
spatial 2

**Suppl. Fig. 5 | Multimodal mapping of CITE-seq data to spatial data**

**a.** Spearman correlation coefficient of predicted genes across three seeds for each method and dataset. **b.** Run times across three seeds for each method and dataset. **c.** Left: Spearman correlation of predicted gene expression aggregated across datasets, summarizing panel (a). Right: run time aggregated across dataset, summarizing panel (b). **d.** Spatial correspondence: The x-axis represents the spatial (Euclidean) distances and the y-axis represents the gene expression (Euclidean) distances across all genes for all cells within the corresponding spatial distance interval. The Spearman correlation is computed between the gene expression distance and the spatial distance. The spatial correspondence showcased here was computed from the drosophila embryo dataset from Li et al.[41] **e.** Example of moscot.space's mapping for the dataset with the highest correlation values (3rd from top left in (a)) with ground truth and predicted expression of two genes: *eve* and *htl*.

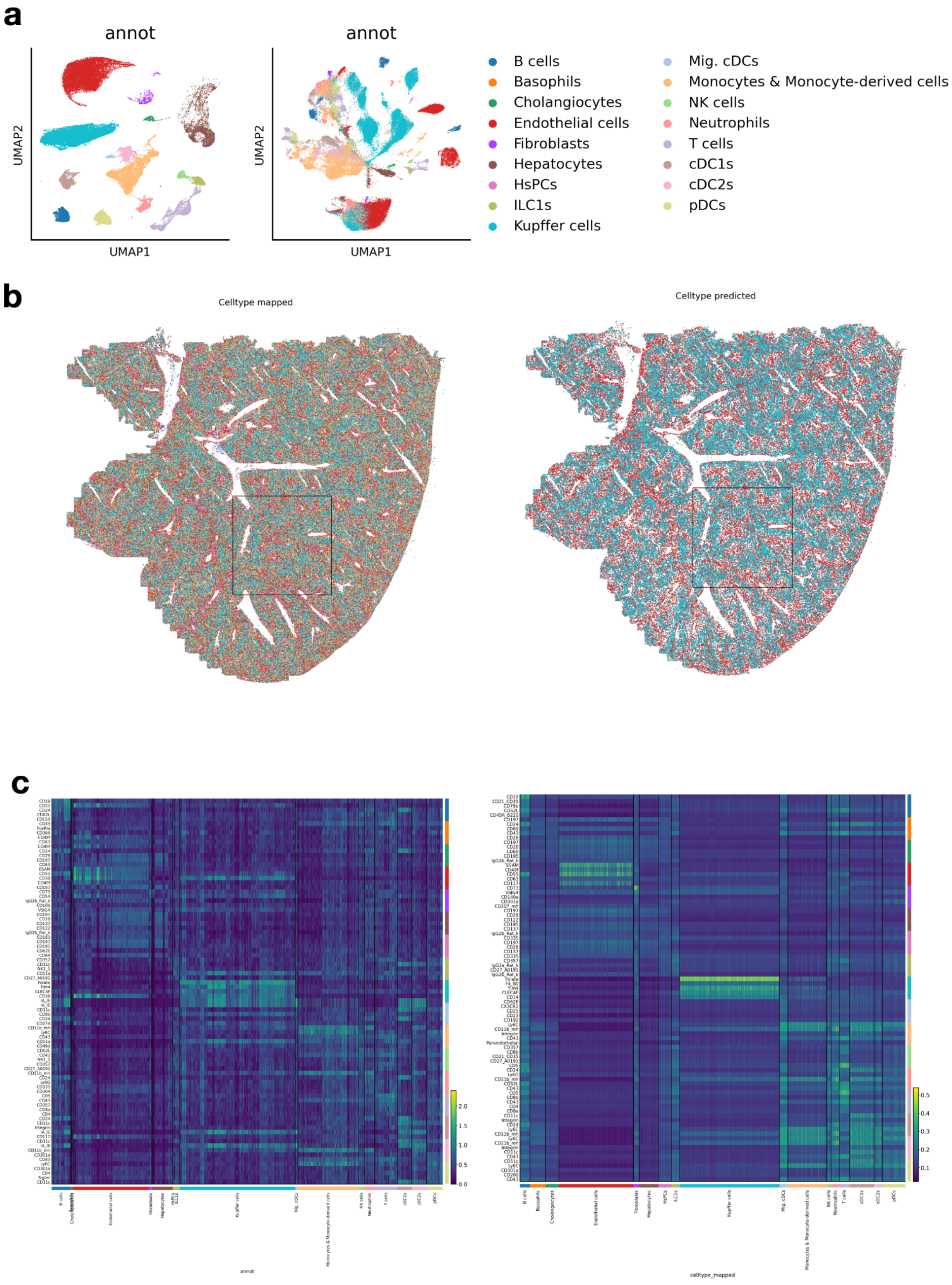

a
annot
annot
UMAP2
UMAP1
B cells
Basophils
Cholangiocytes
Endothelial cells
Fibroblasts
Hepatocytes
HsPCs
ILC1s
Kupffer cells
Mig. cDCs
Monocytes & Monocyte-derived cells
NK cells
Neutrophils
T cells
cDC1s
cDC2s
pDCs
b
Celltype mapped
Celltype predicted
c
annot
celltype_mapped

**Suppl. Fig. 6 | Overview of CITE-seq data and mapped annotations**

**a.** UMAP embedding of single-cell (left) and CITE-seq (right) dataset, respectively. Labels were provided in the original publication. **b.** Cell type annotation mapped in spatial coordinates. All cell types visualized in space (left), and spatial plot of only Kupffer cells (blue) and Endothelial cells (red,right). Boxes in solid lines correspond to insets in Fig. 3. **c.** Top five differentially expressed proteins (five genes/proteins per cluster in rows) in original CITE-seq dataset (left) and predicted cell types and protein expression in space (right).

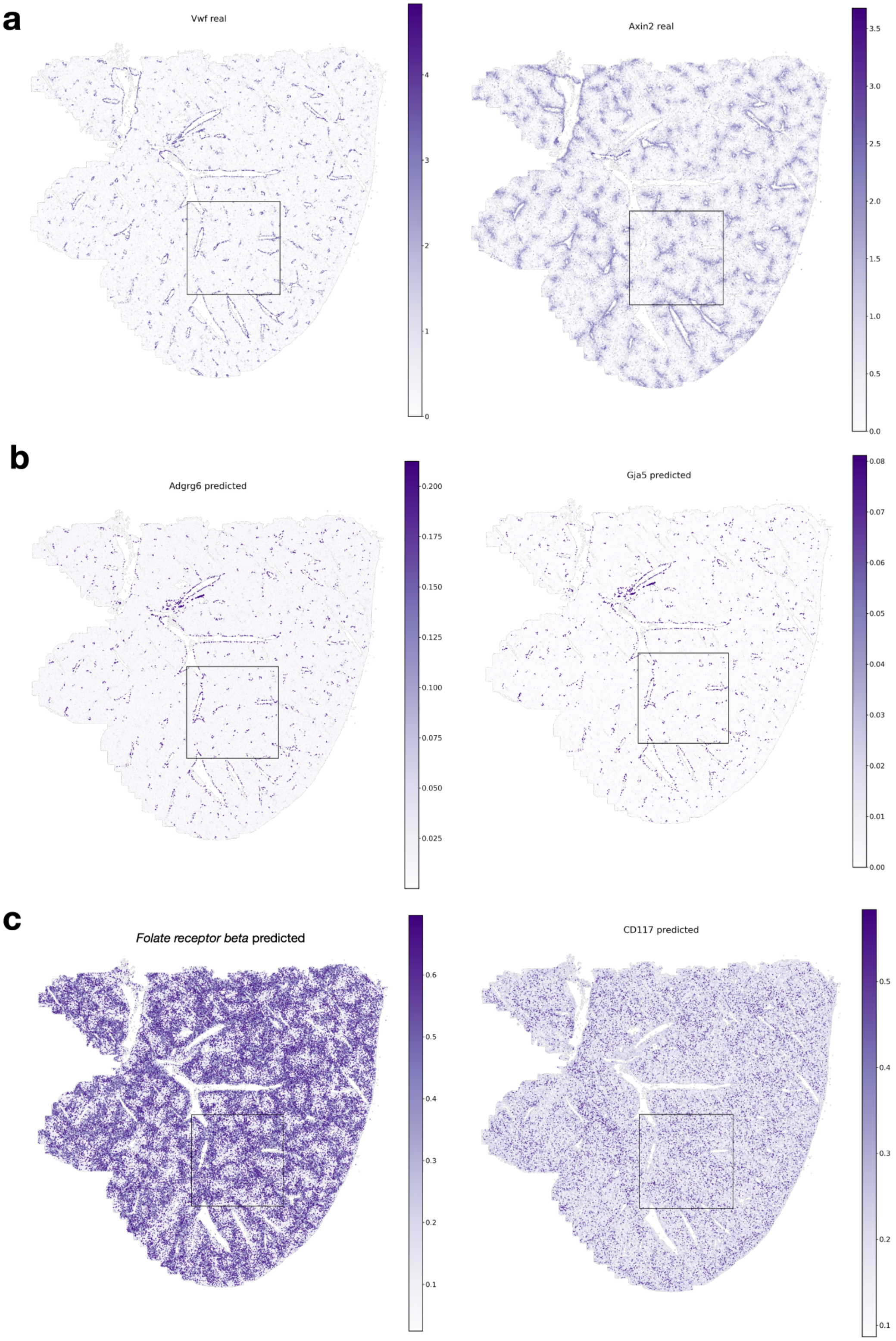
a
Vwf real
0
1
2
3
4
Axin2 real
0.0
0.5
1.0
1.5
2.0
2.5
3.0
3.5
b
Adgrg6 predicted
0.025
0.050
0.075
0.100
0.125
0.150
0.175
0.200
Gja5 predicted
0.00
0.01
0.02
0.03
0.04
0.05
0.06
0.07
0.08
c
Folate receptor beta predicted
0.1
0.2
0.3
0.4
0.5
0.6
CD117 predicted
0.1
0.2
0.3
0.4
0.5

**Suppl. Fig. 7 | Multimodal mapping of CITE-seq data to spatial data**

**a.** Spatial visualization of ground truth genes used to identify veins with *Vwf* (endothelial cells marker) and central veins with *Axin2* (hepatocytes, endothelial cells marker). **b.** Spatial visualization of predicted expression of genes *Adgrg6* and *Gja5*, markers of endothelial cells associated with portal veins. **c.** Spatial visualization of predicted expression of proteins *Folate receptor beta* (marker of Kupffer cells) and *CD117* (marker of endothelial cells associated with central veins). Boxes in solid lines correspond to insets in Fig. 3.

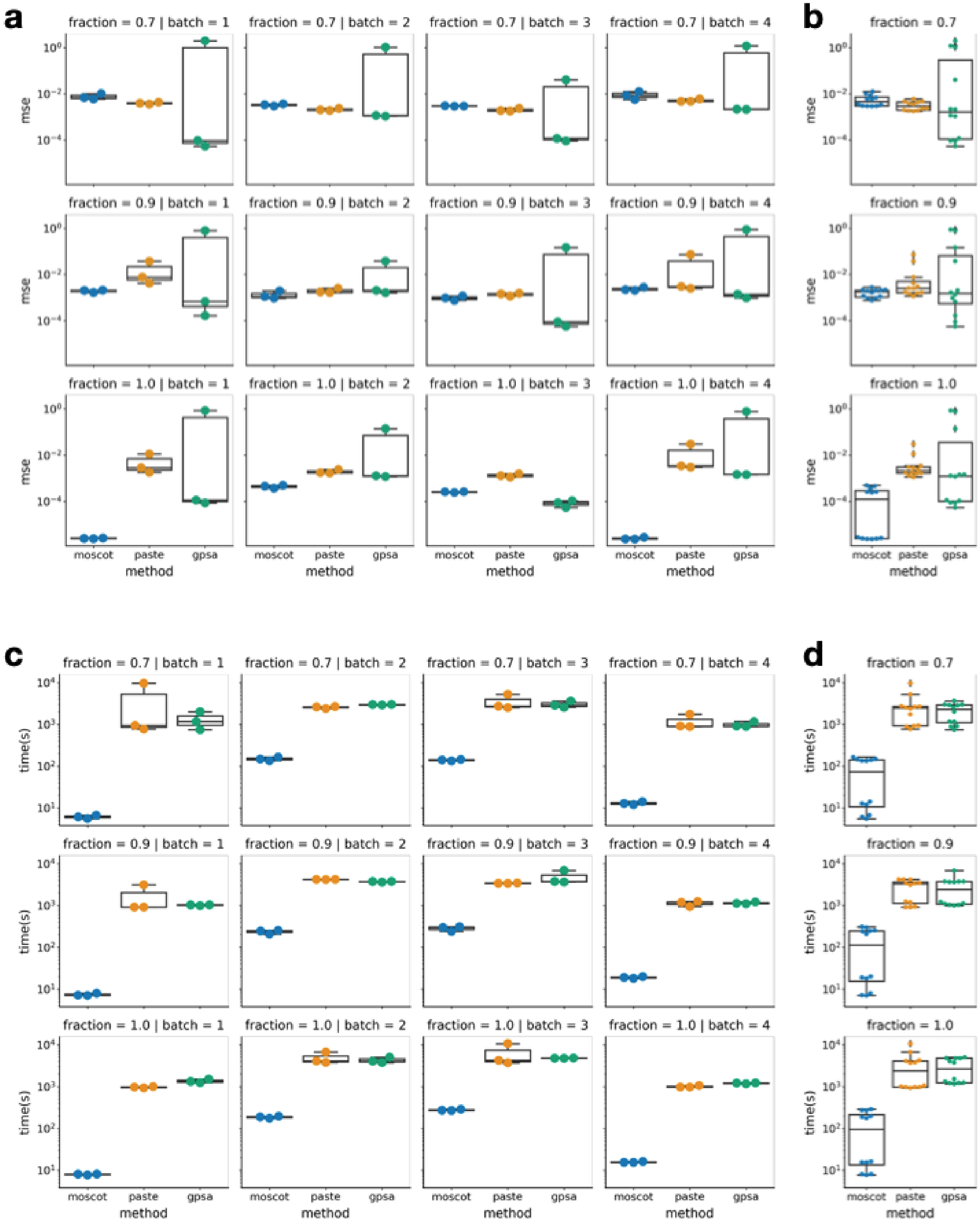
a
b
c
d
fraction = 0.7 | batch = 1
fraction = 0.7 | batch = 2
fraction = 0.7 | batch = 3
fraction = 0.7 | batch = 4
fraction = 0.9 | batch = 1
fraction = 0.9 | batch = 2
fraction = 0.9 | batch = 3
fraction = 0.9 | batch = 4
fraction = 1.0 | batch = 1
fraction = 1.0 | batch = 2
fraction = 1.0 | batch = 3
fraction = 1.0 | batch = 4
fraction = 0.7
fraction = 0.9
fraction = 1.0
mse
time(s)
moscot
paste
gpsa
method

**Suppl. Fig. 8 | Benchmark of spatial alignment datasets**

**a.** Mean squared error (mse) of aligned spots in spatial coordinates across three random seeds for PASTE[13], GPSA[48] and moscot. The columns of the grid correspond to four different simulated datasets, the rows correspond to different fractions of subsampling (1. means no subsampling and the number of points in source and target is exactly the same, 0.7 means that only 70% of the points in both source and target have been kept for the alignment. Subsampling was done to simulate a more realistic noisy scenario). **b.** Aggregated results across datasets. **c.-d.** Run time (c) and aggregated run time (d) of different methods for the experiments outlined before (datasets in columns, fraction of subsampling in rows, last column represents summary over rows).

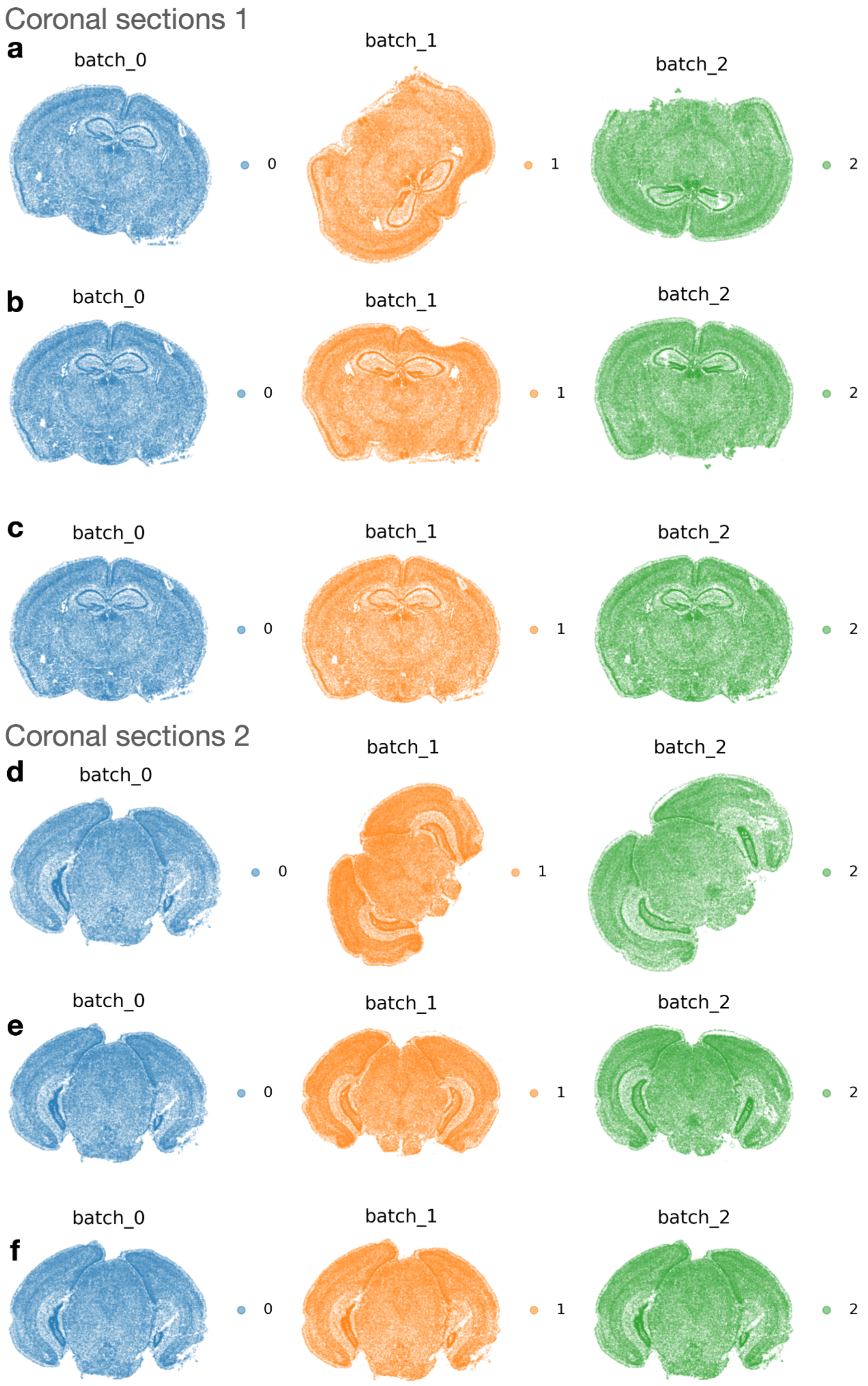

Coronal sections 1
a
batch_0
batch_1
batch_2
0
1
2
b
batch_0
batch_1
batch_2
0
1
2
c
batch_0
batch_1
batch_2
0
1
2
Coronal sections 2
d
batch_0
batch_1
batch_2
0
1
2
e
batch_0
batch_1
batch_2
0
1
2
f
batch_0
batch_1
batch_2
0
1
2

**Suppl. Fig. 9 | Alignment of spatial transcriptomics data of sections of the mouse brain**

**a.** Spatial visualization of the three coronal sections from three different mouse brains before the alignment. **b.** Spatial visualization of the three coronal sections after affine alignment. **c.** Spatial visualization of the three coronal sections after warping alignment. **d.-f.** Original, affine transform and warped transformed tissue slices from the second set of three coronal sections from three different mouse brains.

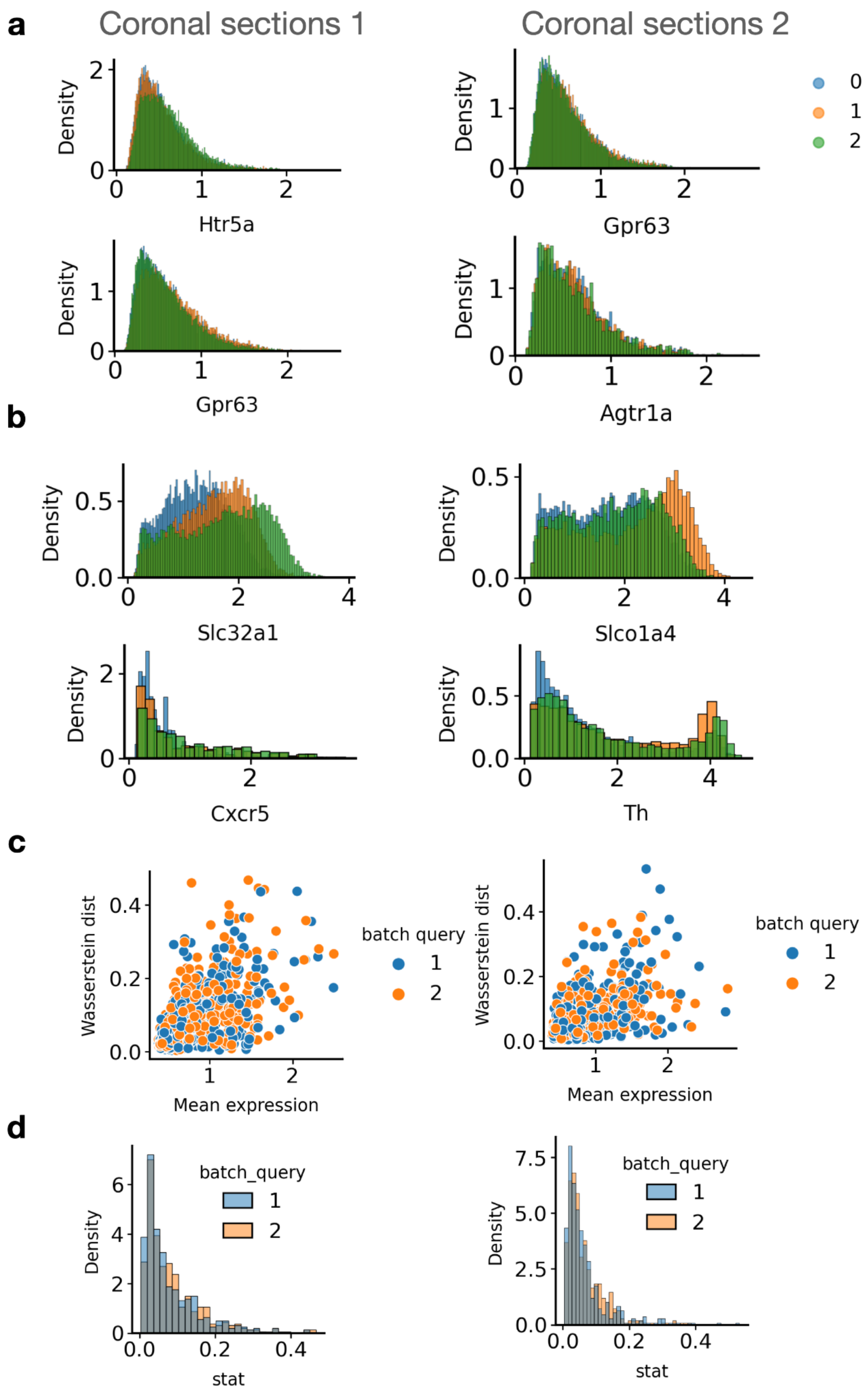

a
Coronal sections 1
Coronal sections 2
Density
Htr5a
Gpr63
Gpr63
Agtr1a
0
1
2
b
Slc32a1
Slco1a4
Cxcr5
Th
c
Wasserstein dist
Mean expression
batch query
1
2
d
batch_query
Density
stat

**Suppl. Fig. 10 | Gene expression consistency on cellular neighbors of aligned slices**

In every row the left panel refers to the first set of coronal sections and the right panel refers to the second set of coronal sections. **a.** Top two genes with lowest L1 Wasserstein distance between gene histograms of reference batch (0) and query batches (1 and 2) for cellular neighbors of aligned slices. **b.** Bottom two genes with highest L1 Wasserstein distance between reference and query batches. **c.** L1 Wasserstein distance across all genes vs. mean gene expression, for both query batches 1 and 2. Interestingly, there is no strong dependency of mean expression showing that the gene expression similarity between cellular neighborhoods of aligned slices is consistent. **d.** Distribution of L1 Wasserstein distances for all genes in query batches 1 and 2.

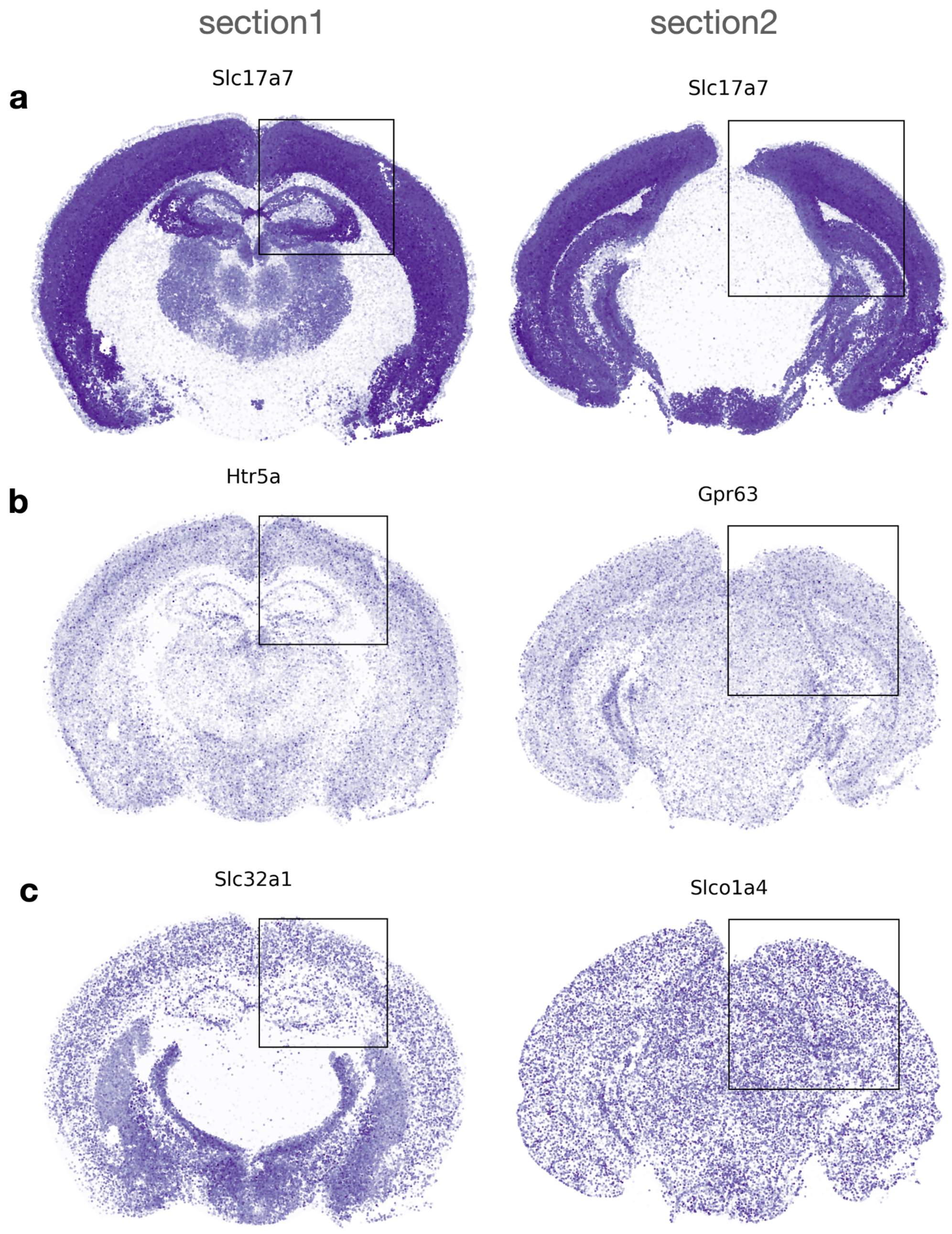


**Suppl. Fig. 11 | Gene expression consistency on cellular neighbors of aligned slices**

**a.** Spatial visualization of *Slc17a7.* **b.** Most consistent genes according to consistency analysis (Methods): *Htra5* for brain coronal sections 1 and *Gpr63* for brain coronal sections 2. **c.** Least consistent genes according to consistency analysis: *Slc32a1* for brain coronal sections 1 and *Slco1a4* for brain coronal sections 2. Boxes in solid lines correspond to insets in Fig. 3.

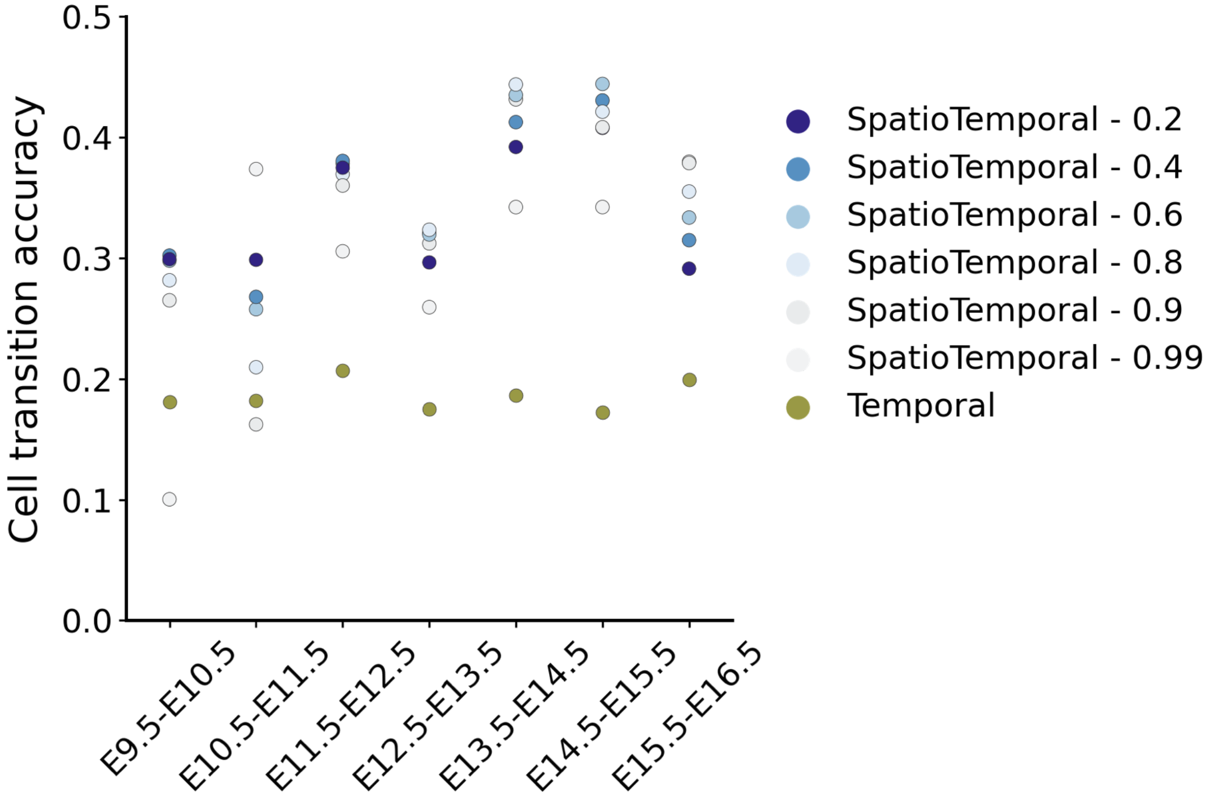


**Suppl. Fig. 12 | moscot.spatiotemporal is robust to hyperparameter choice**

Evaluation and comparison of moscot.spatiotemporal, assessed by the accuracy on curated transitions by developmental stages, presented for different values of the FGW interpolation parameter (Methods and Supplementary Table 5).

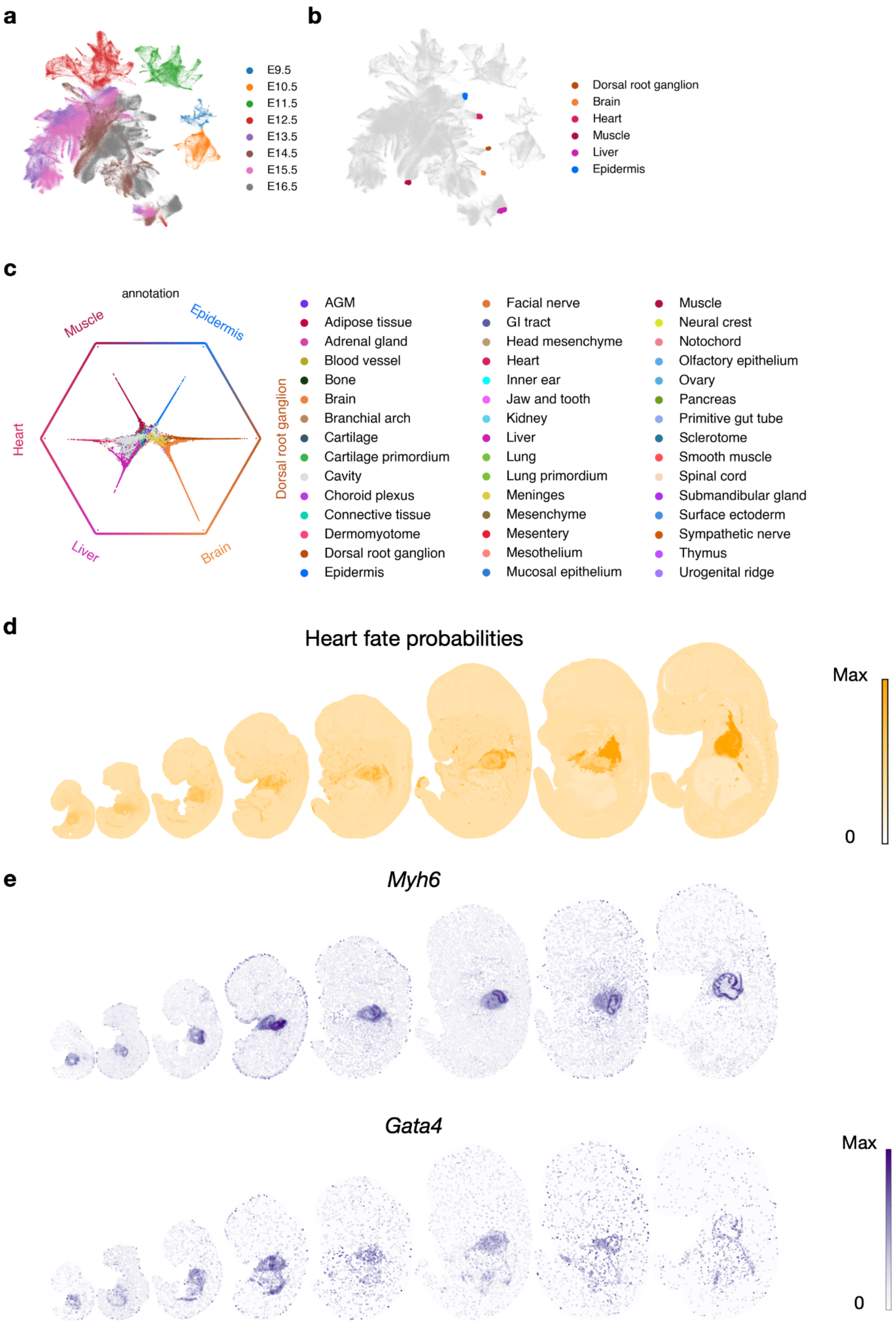
a
E9.5
E10.5
E11.5
E12.5
E13.5
E14.5
E15.5
E16.5
b
Dorsal root ganglion
Brain
Heart
Muscle
Liver
Epidermis
c
annotation
Muscle
Epidermis
Heart
Dorsal root ganglion
Liver
Brain
AGM
Adipose tissue
Adrenal gland
Blood vessel
Bone
Brain
Branchial arch
Cartilage
Cartilage primordium
Cavity
Choroid plexus
Connective tissue
Dermomyotome
Dorsal root ganglion
Epidermis
Facial nerve
GI tract
Head mesenchyme
Heart
Inner ear
Jaw and tooth
Kidney
Liver
Lung
Lung primordium
Meninges
Mesenchyme
Mesentery
Mesothelium
Mucosal epithelium
Muscle
Neural crest
Notochord
Olfactory epithelium
Ovary
Pancreas
Primitive gut tube
Sclerotome
Smooth muscle
Spinal cord
Submandibular gland
Surface ectoderm
Sympathetic nerve
Thymus
Urogenital ridge
d
Heart fate probabilities
Max
0
e
Myh6
Gata4
Max
0

**Suppl. Fig. 13 | Analysis of development lineages by interfacing moscot.spatiotemporal with CellRank**

**a.** UMAP representation of the spatiotemporal atlas of mouse embryogenesis (MOSTA) over eight time points, from E9.5 to E16.5[3] colored by time points. **b.** UMAP colored by macrostates identified by CellRank. **c.** Projection of cell's absorption probabilities towards identified macrostates. Cells colored by lineage annotation. **d.** CellRank fate probabilities for heart fate visualized in spatial coordinates **e.** Spatial visualization of driver genes identified for the heart development lineage, *Myh6* (top) and *Gata4* (bottom).

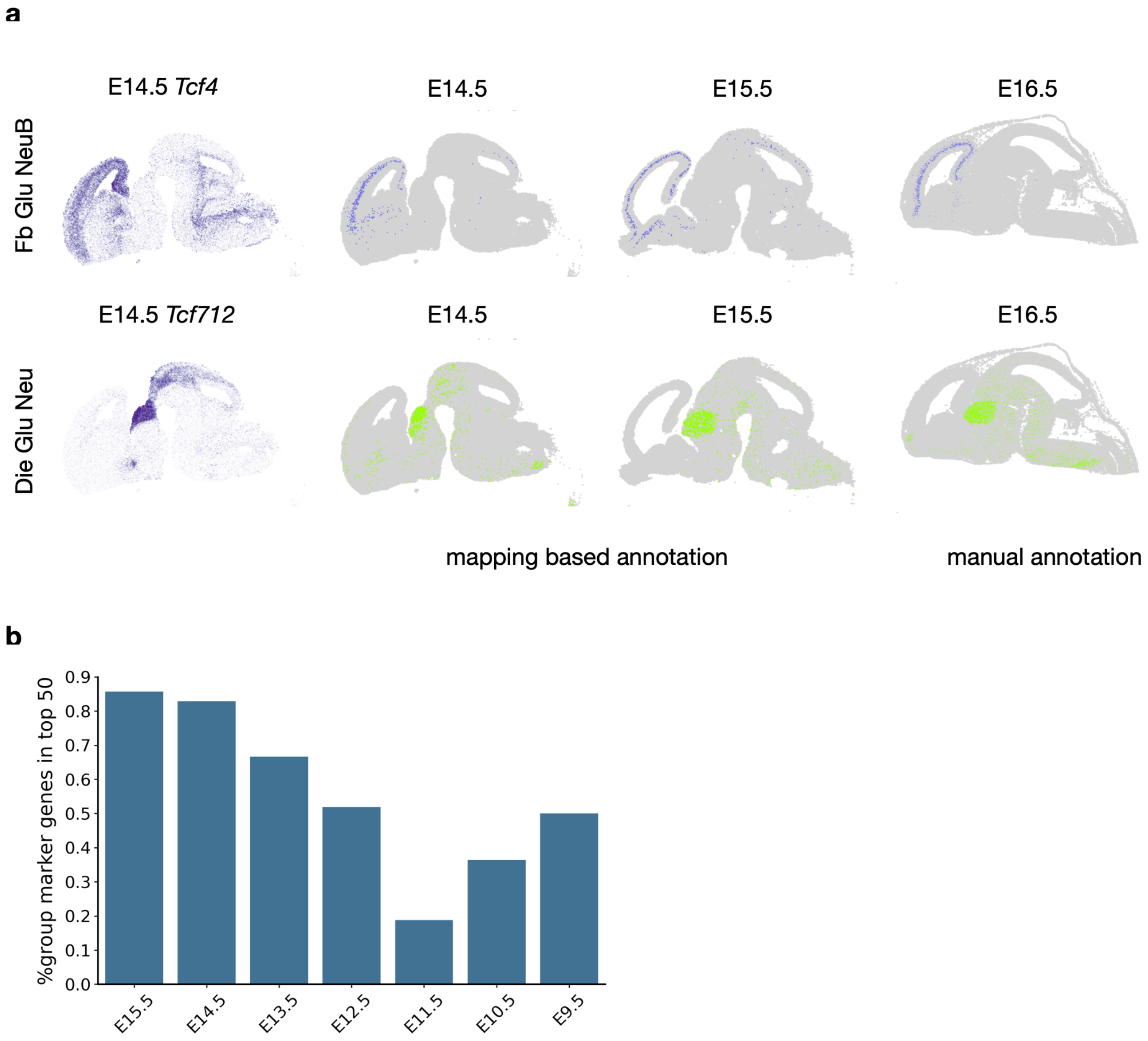


**Suppl. Fig. 14 | moscot.spatiotemporal allows for accurate inference of brain cell type annotations**

**a.** Spatial visualization of mapping based annotations of brain cells focusing on specific cell types, Fb Glu NeuB (forebrain glutamatergic neuroblast, top row) and Die Glu Neu (diencephalon glutamatergic neuron, bottom row). Columns, left to right, cell type reported marker gene, cells assigned to cell type at E14.5 and E15.5, manual reference annotation at E16.5. **b.** Bar plot visualizing the percentage of group marker genes found in the top 50 genes associated with the mapping based annotated group (Methods).

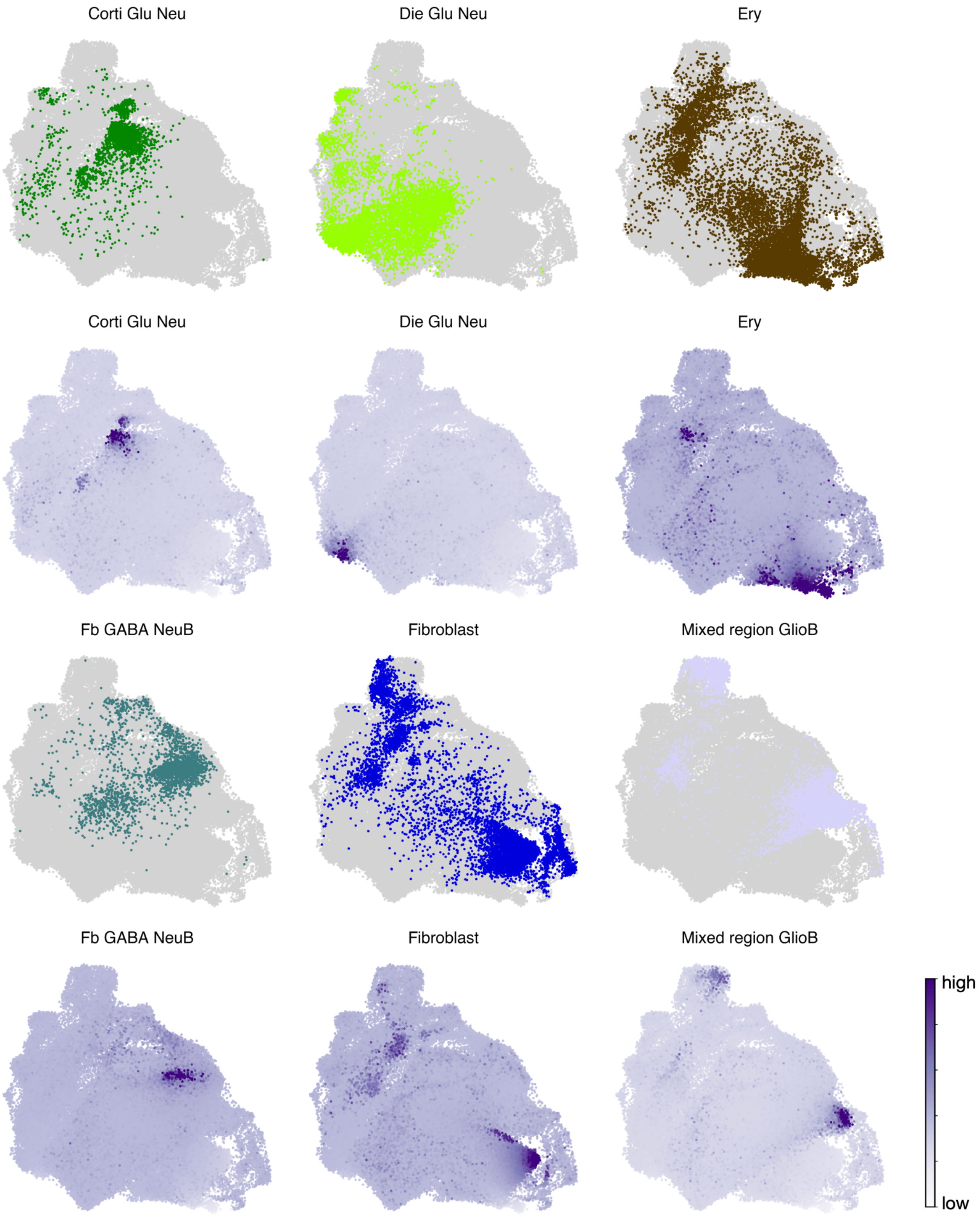


**Suppl. Fig. 15 | Terminal states of brain cells as inferred by interfacing moscot.spatiotemporal with CellRank**

Each subplot provides a visualization for a different terminal state. The UMAPs contain brain cells from E13.5-E16.5 and are colored according to mapping based annotation (first and third row) or CellRank fate probabilities (second and fourth row).

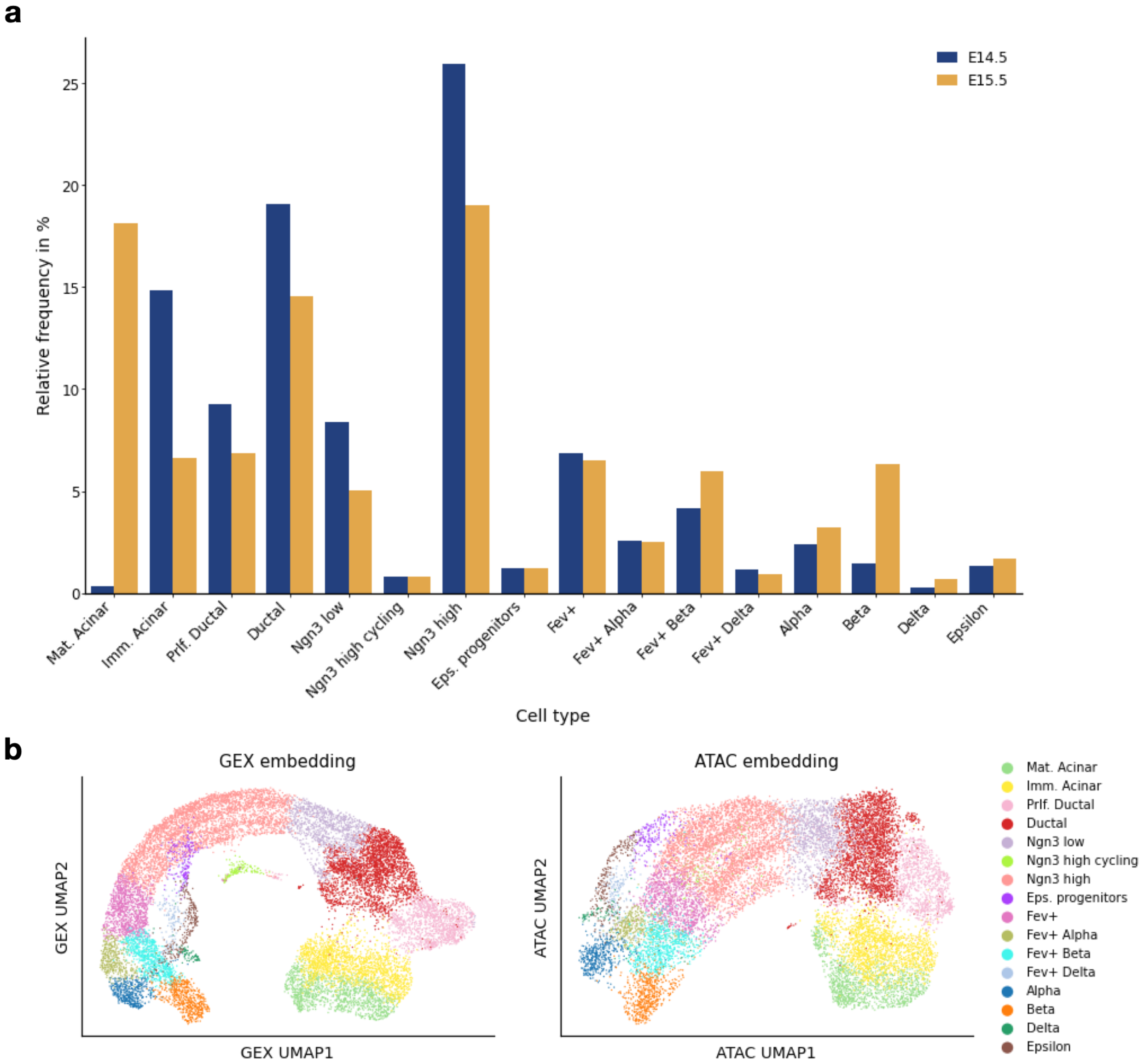


**Suppl. Fig. 16 | Summary statistics and visualization of the pancreatic endocrinogenesis dataset**

**a.** Distribution of cell types per time point. **b.** UMAP embeddings based on graphs constructed from gene expression and open chromatin accessibility, respectively.

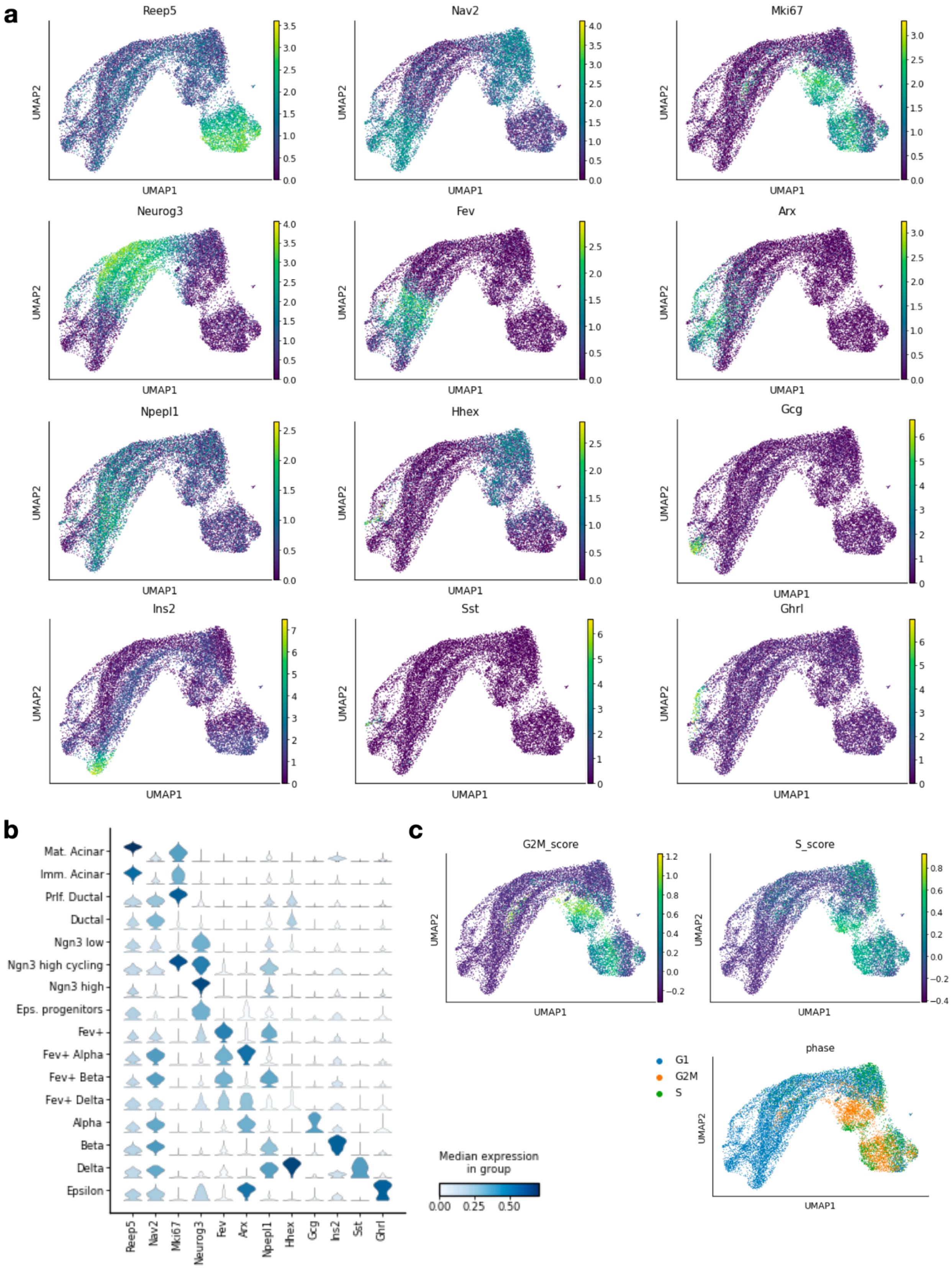

a
Reep5
Nav2
Mki67
Neurog3
Fev
Arx
Npepl1
Hhex
Gcg
Ins2
Sst
Ghrl
UMAP1
UMAP2
b
Mat. Acinar
Imm. Acinar
Prlf. Ductal
Ductal
Ngn3 low
Ngn3 high cycling
Ngn3 high
Eps. progenitors
Fev+
Fev+ Alpha
Fev+ Beta
Fev+ Delta
Alpha
Beta
Delta
Epsilon
Median expression in group
0.00 0.25 0.50
c
G2M_score
S_score
phase
G1
G2M
S

**Suppl. Fig. 17 | Marker gene expression in pancreatic endocrinogenesis**

**a.** Processed gene expression (processed with *scanpy.pp.normalize_total* and *scanpy.pp.log1p*) of selected marker genes. In particular, *Reep5* is a marker for acinar cells, *Nav2* for ductal cells, *Mki67* for proliferative ductal cells, *Neurog3* for Ngn3$^{low}$ and Ngn3$^{high}$ cells, *Fev* for Fev+ cells, *Arx* for Fev+ alpha cells, *Npepl1* for Fev+ beta cells, *Hhex* for Fev+ delta cells (and delta cells), *Gcg* for alpha cells, *Ins2* for beta cells, *Sst* for delta cells, and *Ghrl* for epsilon cells. **b.** Min-max normalized processed gene expression per cell type. **c.** G2M score, S score and proliferation phase as computed by scanpy's *score_genes_cell_cycle* function.

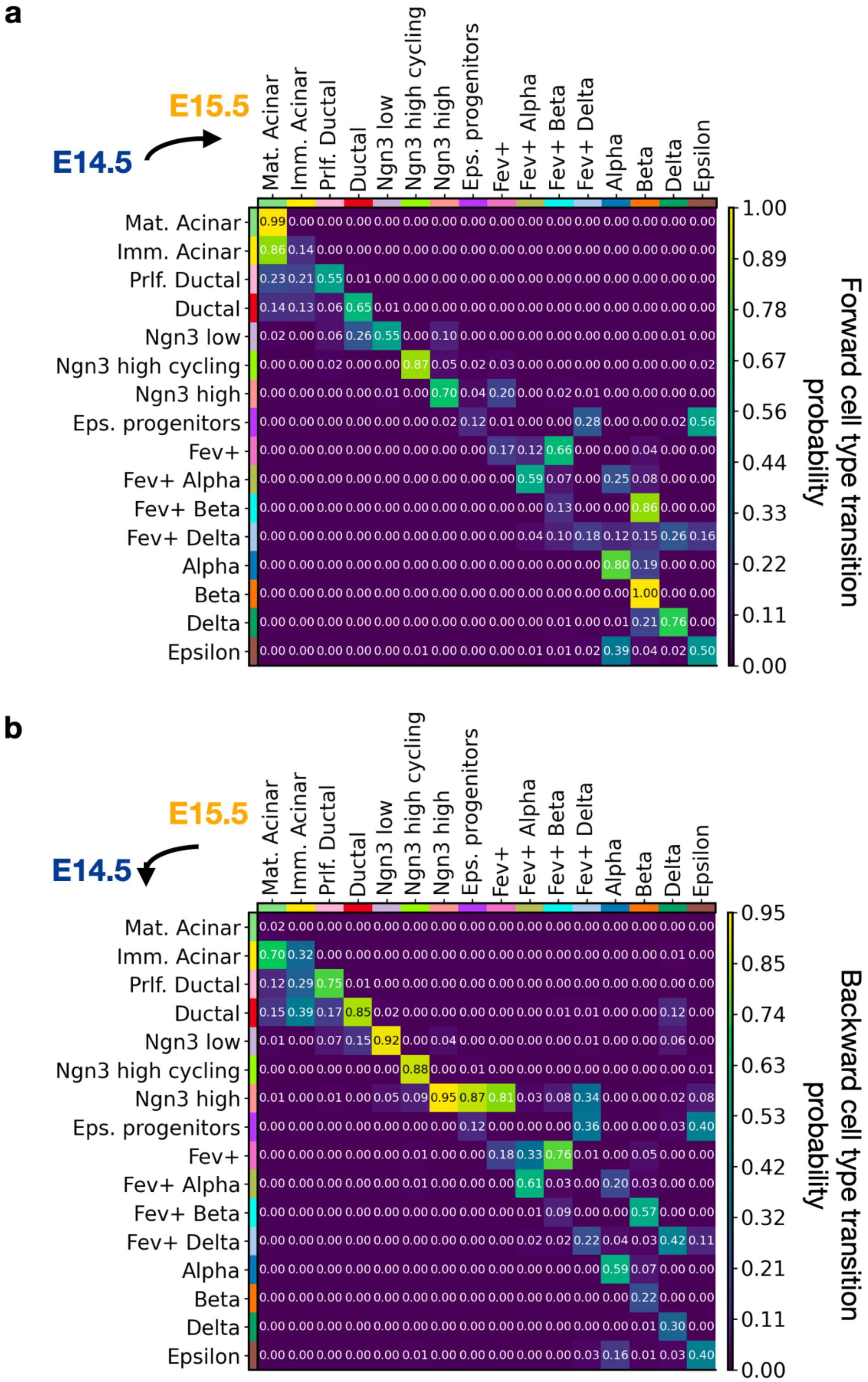
a
E15.5
E14.5
Mat. Acinar
Imm. Acinar
Prlf. Ductal
Ductal
Ngn3 low
Ngn3 high cycling
Ngn3 high
Eps. progenitors
Fev+
Fev+ Alpha
Fev+ Beta
Fev+ Delta
Alpha
Beta
Delta
Epsilon
1.00
0.89
0.78
0.67
0.56
0.44
0.33
0.22
0.11
0.00
Forward cell type transition probability
b
E15.5
E14.5
0.95
0.85
0.74
0.63
0.53
0.42
0.32
0.21
0.11
0.00
Backward cell type transition probability

**Suppl. Fig. 18 | Cell type transition probabilities of the full pancreatic endocrinogenesis dataset**

**a.** Transition probabilities obtained by moscot.time and aggregated per cell type. Each row sums up to 1, hence each entry (i,j) denotes the probability of cell type i in E14.5 to transition to cell type j in E15.5. **b.** Transition probabilities obtained by moscot.time, but this time each column adds up to 1, hence each entry (i,j) denotes the probability of cell type i in E14.5 to be an ancestor of cell type j in E15.5.

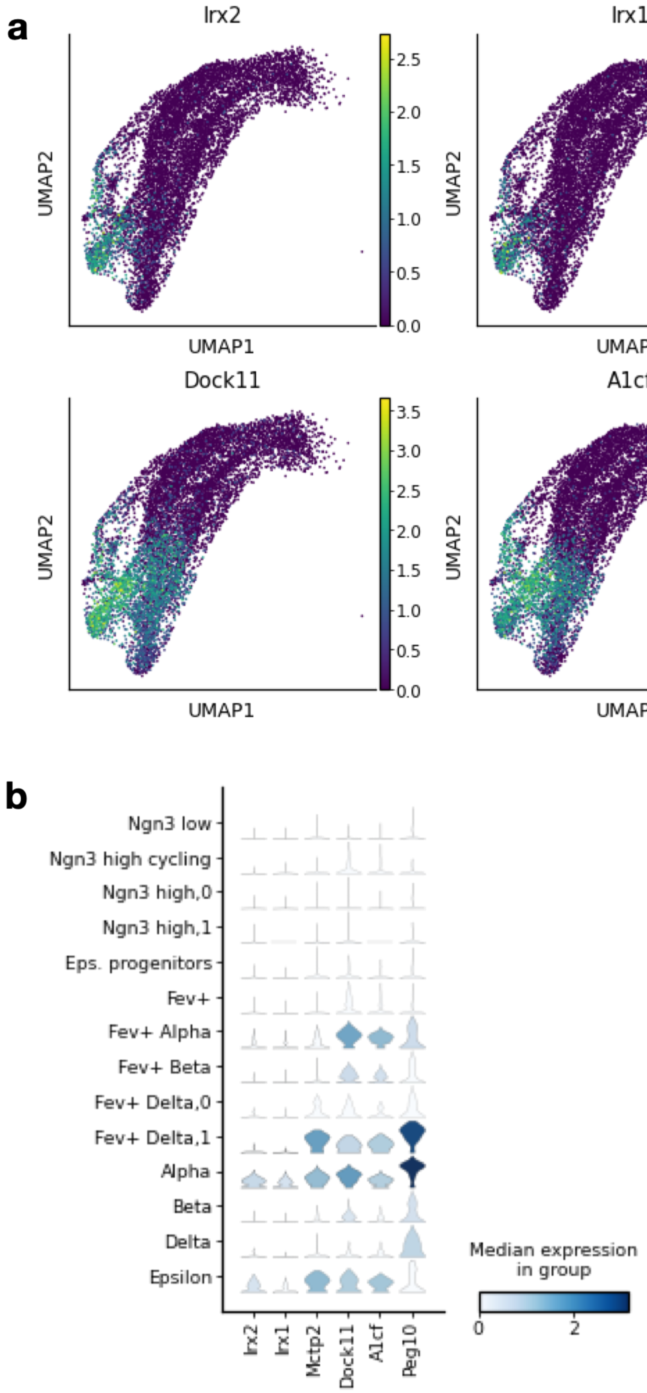


**Suppl. Fig. 19 | Marker genes for the transition from epsilon cells to alpha cells.**

**a.** By leveraging moscot's capability of identifying marker features (Methods) we identify genes which are highly expressed in epsilon cells that are likely to transition towards an alpha cell state. *Irx2* has been reported as a key TF for these cell states[68], while *Irx1* was reported for the analogue cell states in human pancreatic endocrinogenesis[100]. *Peg10* has been reported as a driver gene for alpha cells[101], while *Mctp2, Dock11,* and *A1cf* have not been reported in this context before. **b.** Processed gene expression of considered genes per cell type.

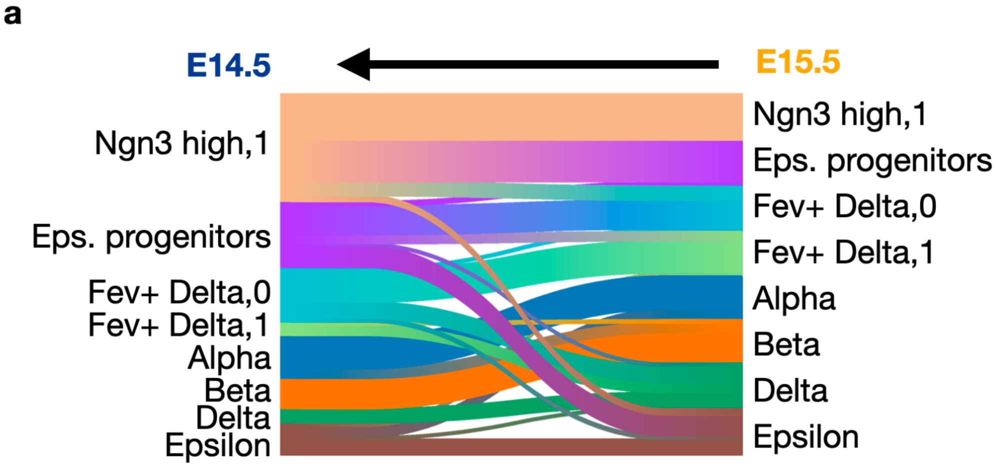


**Suppl. Fig. 20 | Ancestors of refined cell types obtained by moscot.time**

**a.** Subset of refined cell types at E15.5 and their ancestors in E14.5. Only ancestries with a probability of at least 0.05 are visualized.

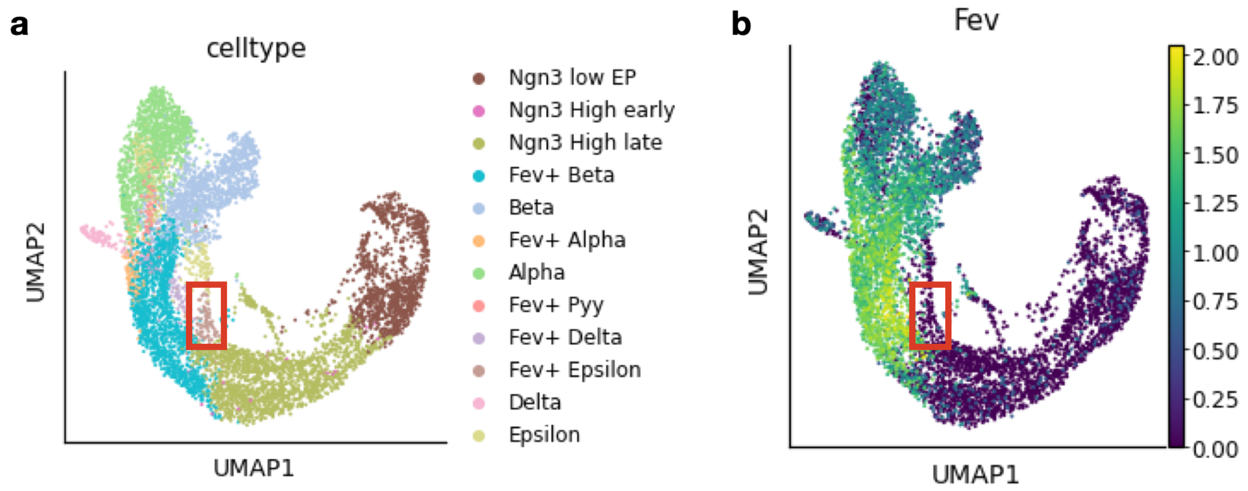


**Suppl. Fig. 21 | Fev expression in the pancreatic endocrinogenesis dataset published by Bastidas-Ponce et al.** [65]

**a.** UMAP embedding colored by cell type of the pancreatic endocrinogenesis dataset published by Bastidas-Ponce et al.[65]. The data was subset to time points E14.5 and E15.5 as well as to endocrine cells and their progenitors. Subsequently, the data was preprocessed by normalization, log1p-transformation and PCA computation before calculation of the neighborhood graph, based on which the UMAP was computed. **b.** Gene expression of *Fev* after preprocessing as described above. The red box highlights the cells annotated as Fev+ epsilon, showing there is no expression of *Fev* in the Fev+ epsilon population.

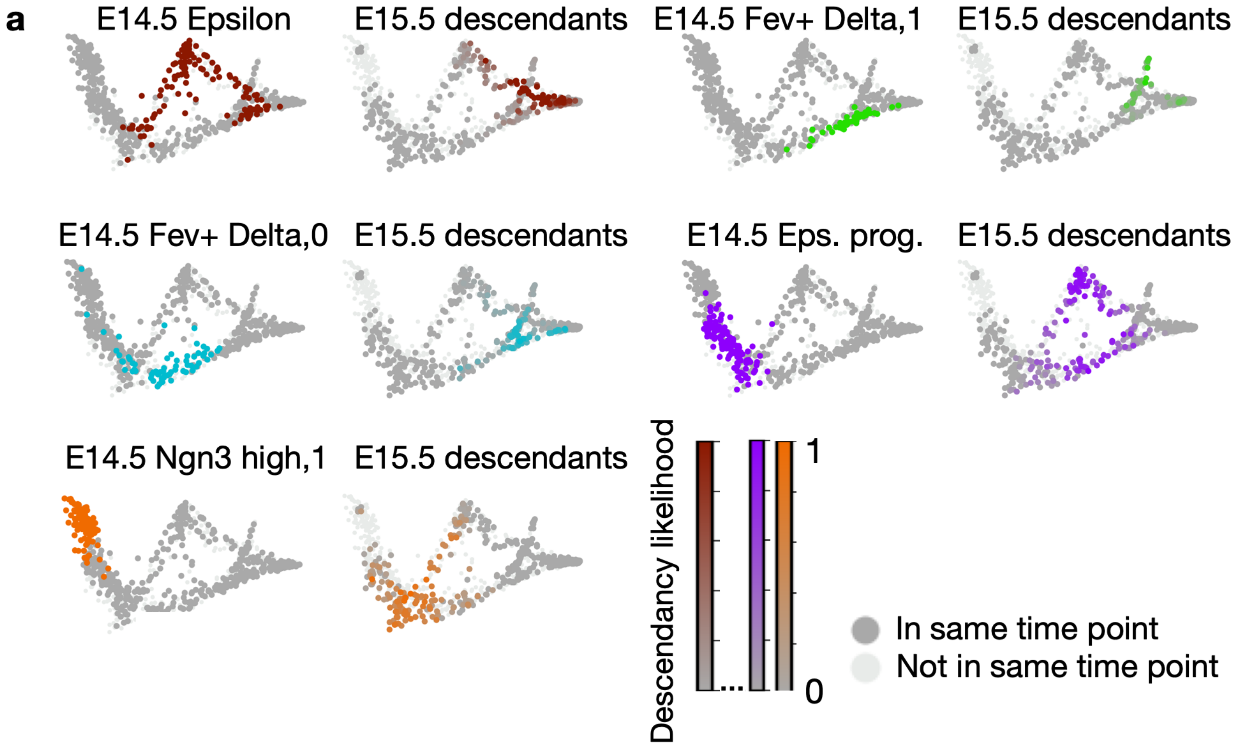


**Suppl. Fig. 22 | Descendants of endocrine and endocrine progenitors on a diffusion map**

**a.** Cell types at E14.5 and their respective descendants at E15.5 computed by moscot.time on a diffusion map.

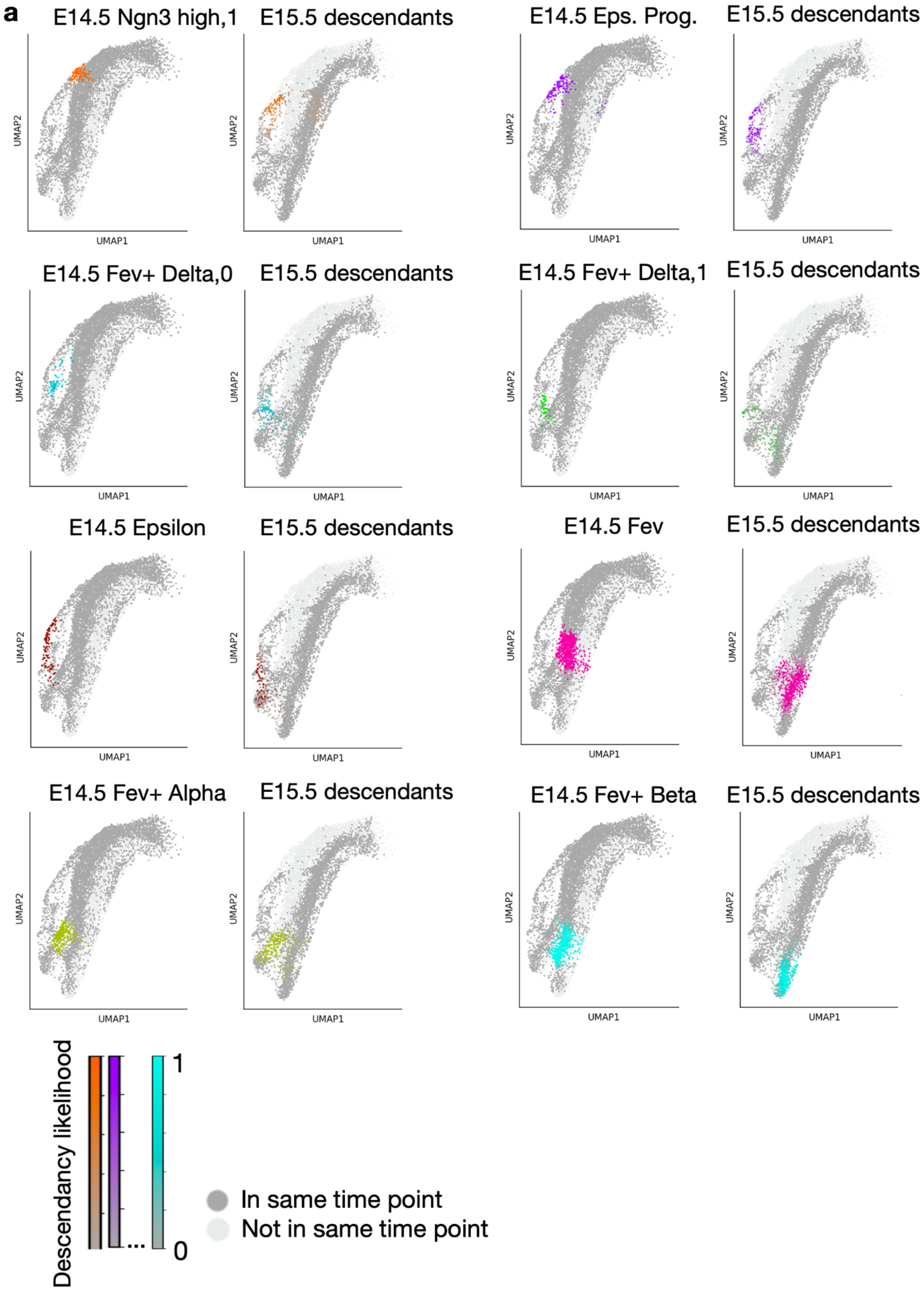
a
E14.5 Ngn3 high,1
E15.5 descendants
E14.5 Eps. Prog.
E15.5 descendants
E14.5 Fev+ Delta,0
E15.5 descendants
E14.5 Fev+ Delta,1
E15.5 descendants
E14.5 Epsilon
E15.5 descendants
E14.5 Fev
E15.5 descendants
E14.5 Fev+ Alpha
E15.5 descendants
E14.5 Fev+ Beta
E15.5 descendants
UMAP1
UMAP2
Descendancy likelihood
1
0
In same time point
Not in same time point

**Suppl. Fig. 23 | Descendancy of endocrine and endocrine progenitors on a UMAP**

**a.** Cell types at E14.5 and their respective descendants at E15.5 computed by moscot.time on a UMAP of the full endocrine branch.

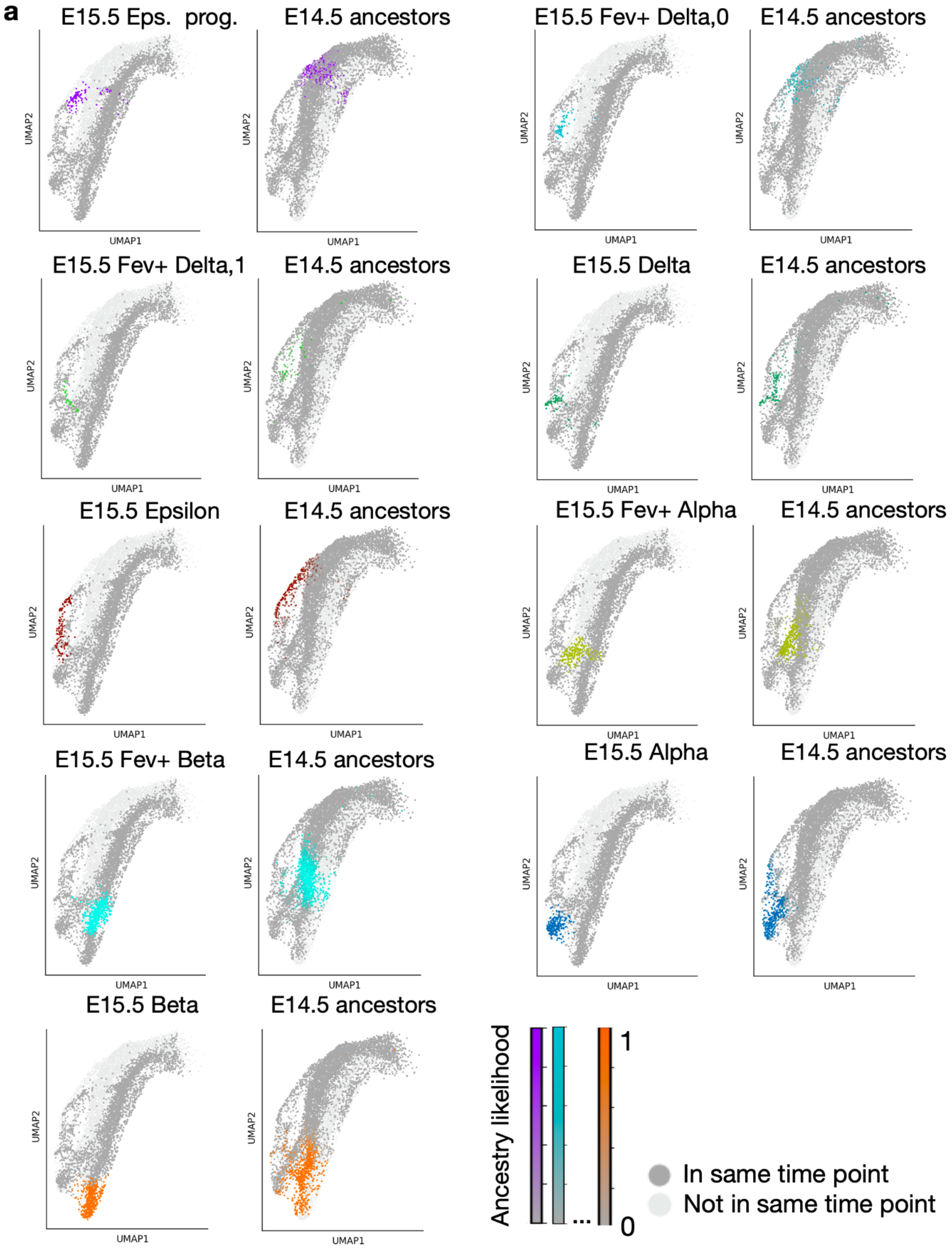
a
E15.5 Eps. prog.
E14.5 ancestors
E15.5 Fev+ Delta,0
E14.5 ancestors
E15.5 Fev+ Delta,1
E14.5 ancestors
E15.5 Delta
E14.5 ancestors
E15.5 Epsilon
E14.5 ancestors
E15.5 Fev+ Alpha
E14.5 ancestors
E15.5 Fev+ Beta
E14.5 ancestors
E15.5 Alpha
E14.5 ancestors
E15.5 Beta
E14.5 ancestors
UMAP1
UMAP2
Ancestry likelihood
1
0
...
In same time point
Not in same time point

**Suppl. Fig. 24 | Ancestry of endocrine and endocrine progenitors on a UMAP**

**a.** Cell types at E15.5 and their respective ancestors at E15.5 computed by moscot.time on a UMAP of the full endocrine branch.

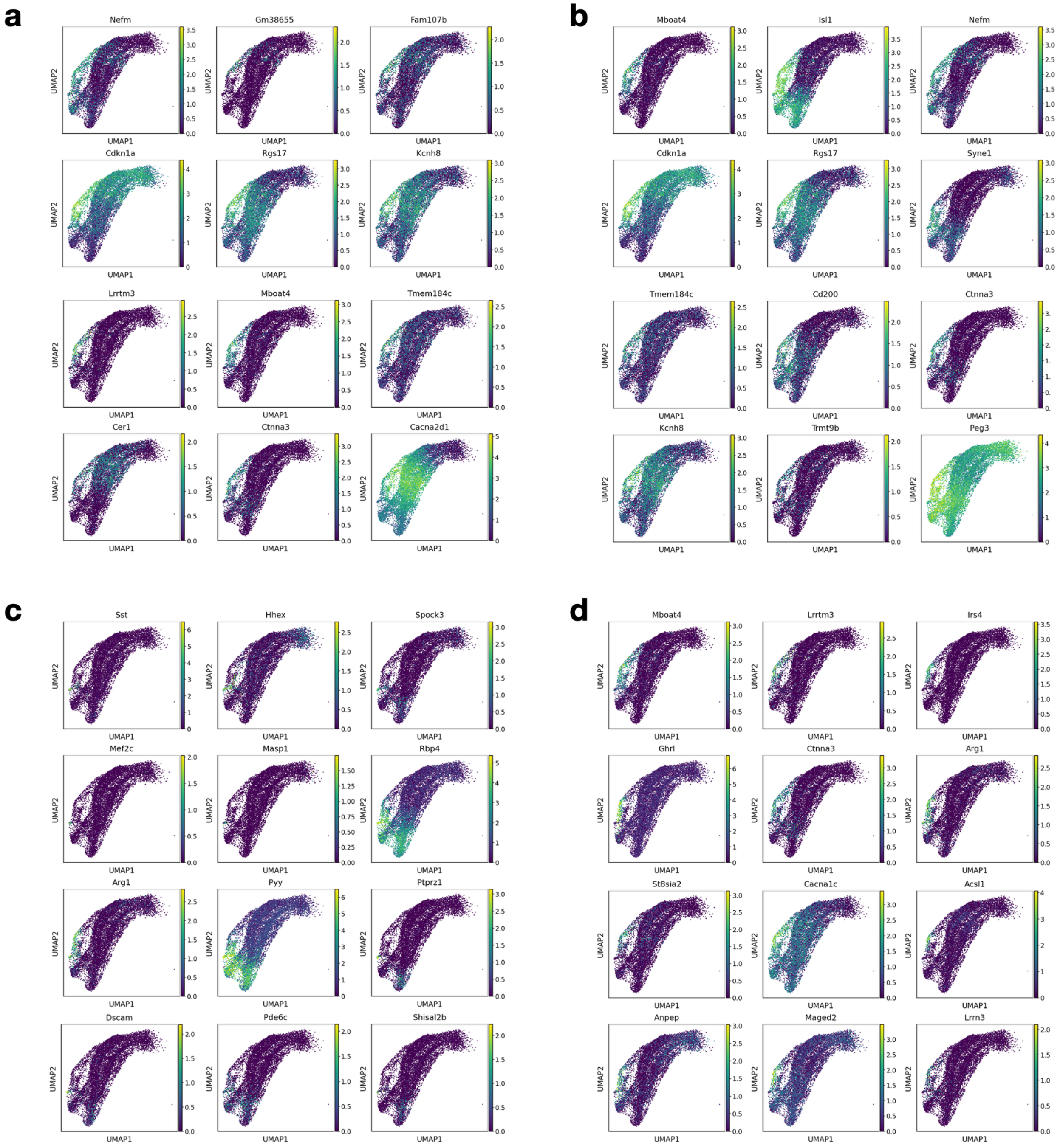


**Suppl. Fig. 25 | Marker genes computed by moscot.time for delta/epsilon lineage**

**a.** Marker genes for epsilon progenitors as computed by moscot.time (Methods). While a few markers have been reported (*Mboat*[102]), or were considered in different contexts (e.g. the cell cycle inhibitor *Cdkn1a*[103], *Rgs17*[104], *Cacna2d1*[105]), we report genes which have been less studied in the context of pancreatic endocrinogenesis (*Gm38655, Fam107b, Kcnh8, Lrrtm3* (Supplementary Fig. 29)*, Tmem184c, Cer1*) **b.** Marker genes for the Fev+ delta population (Fev+ delta-0 and Fev+ delta-1 combined due to their high similarity in gene expression and

ATAC profile). While we recover genes which we also report for epsilon progenitors in panel a, there are new marker genes, such as the well studied *Isl1* (Supplementary Fig. 31) and *Peg3* (transcription factor reported for beta cells[100]), and less studied, but significantly expressed genes (*Syne1, Cd200, Trmt9b)*. **c.** Marker genes for delta cells. While *Sst* and *Hhex* confirm the reliability of moscot's marker gene recovery method as the most well-known marker genes for delta cells (also *Spock3*[106], *Mef2c*[68], *Pyy*[65]), we also report new genes in this context: *Masp1, Rbp4, Ptprz1, Dscam, Pde6c, Shisal2b*. Moreover, *Arg1* is recovered due to the multipotency of Fev+ delta cells, and other genes shared with markers from Fev+ delta cells or epsilon progenitor cells are recovered. **d.** Marker genes for the epsilon population. Besides already considered genes we report the well-known markers *Irs4* [106] and *Ghrl*[69], as well as *Ctnna3* (Fig. 5l), *St8sia2, Cacna1c*[107]*, Acsl1, Anpep, Maged2,* and *Lrrn3.*

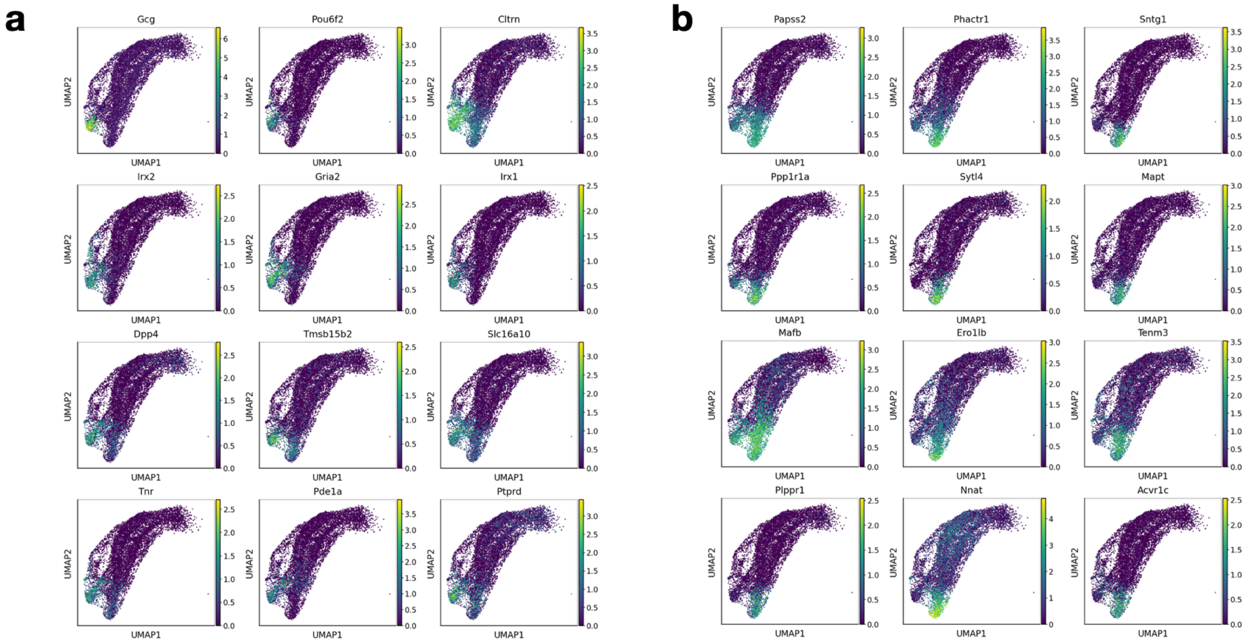


**Suppl. Fig. 26 | Marker genes computed by moscot.time for alpha and beta cells**

**a.** Marker genes of alpha cells computed with moscot.time. We recover well-known marker genes such as *Gcg, Pou6f2, Irx2*[68], *Irx1*[107], but also less reported ones like *Cltrn, Gria2, Dpp4, Tmsb15b2, Sl16a10, Tnr, Pde1a,* and *Ptprd*. **b.** Marker genes of beta cells. We recover known marker genes such as *Papss2*[68], *Ppp1r1a*[108], *Sytl4*[109], *Mafb*[76], *Ero1lb* [109], *Nnat* [110] and *Acvr1c* [111] but also less known ones like *Phactr1, Sntg1, Mapt, Tenm3,* and *Plppr1.*

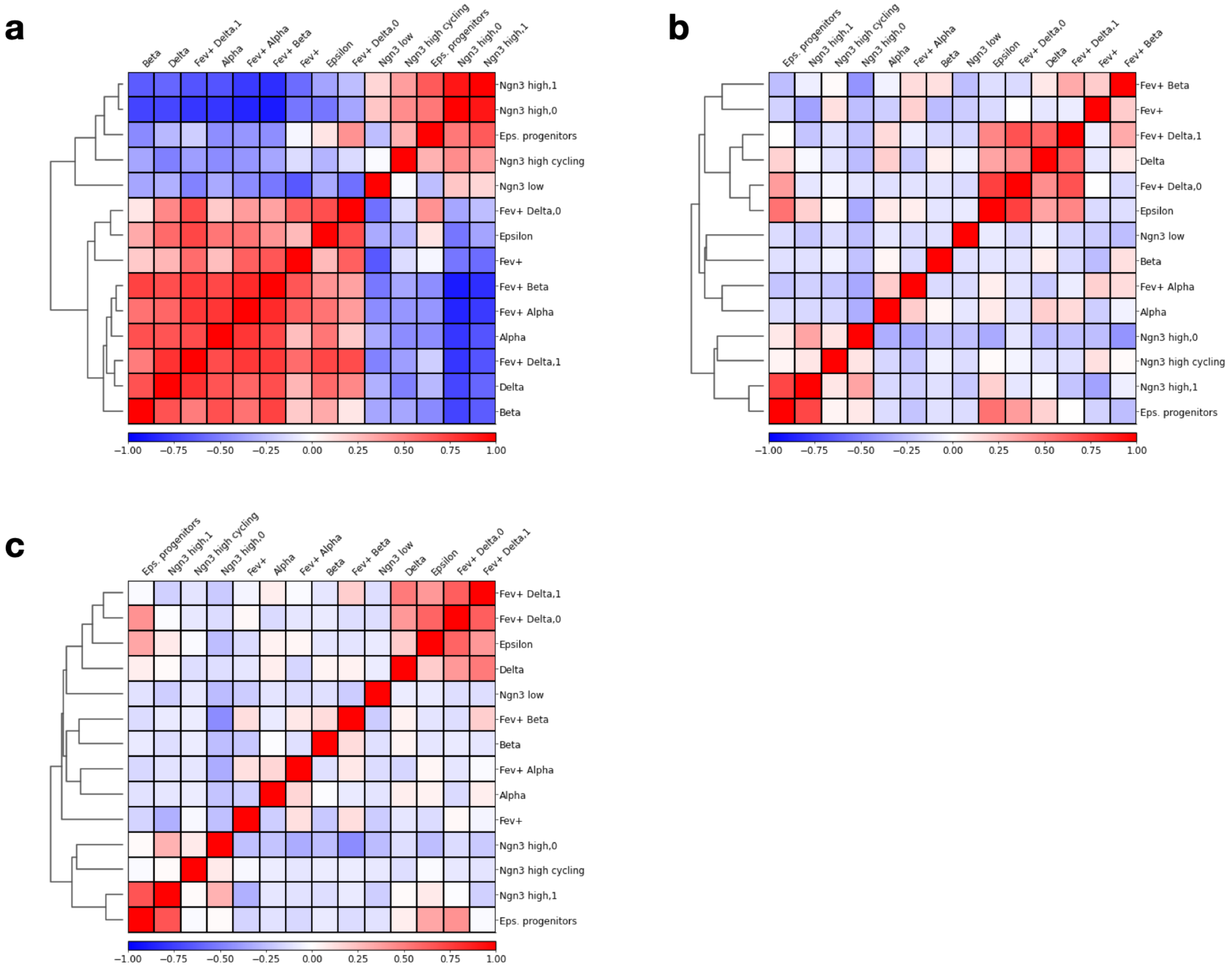


**Suppl. Fig. 27 | Similarity of cell types based on different modalities**

**a.** Aggregated correlation matrix of refined cell types based on processed gene expression, computed via scanpy's *dendrogram* function, followed by *scanpy.pl.correlation_matrix*. The gene expression data was preprocessed by normalization (sc.pp.*normalize_total*) and log1p-transformation, followed by 30-dimensional PCA computation. **b.** Aggregated correlation matrix of cell types based on processed ATAC peak counts, computed via *scanpy.tl.dendrogram* followed by *scanpy.pl.correlation_matrix*. The peak counts were preprocessed using tfidf-transformation (*muon.atac.pp.tfidf*), followed by normalization and log1p-transformation, before computing a singular value decomposition and removal of dimensions which are highly correlated with library size. **c.** Aggregated correlation matrix of cell types based on both gene expression and open chromatin accessibility. After scaling both modalities to unit variance, the processed gene expression was concatenated with the processed LSI embedding.

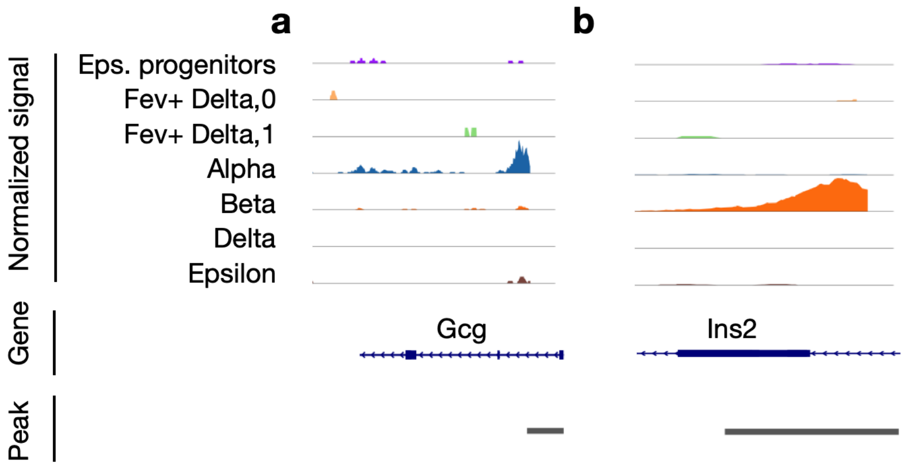


**Suppl. Fig. 28 | Chromatin accessibility at the promoter regions of *Gcg* and *Ins2***

**a.** Open chromatin around the promoter region of *Gcg* (peak 2-62474530-62483650). **b.** Chromatin accessibility around the promoter region of *Ins2* (peak 7-142678656-142679685).

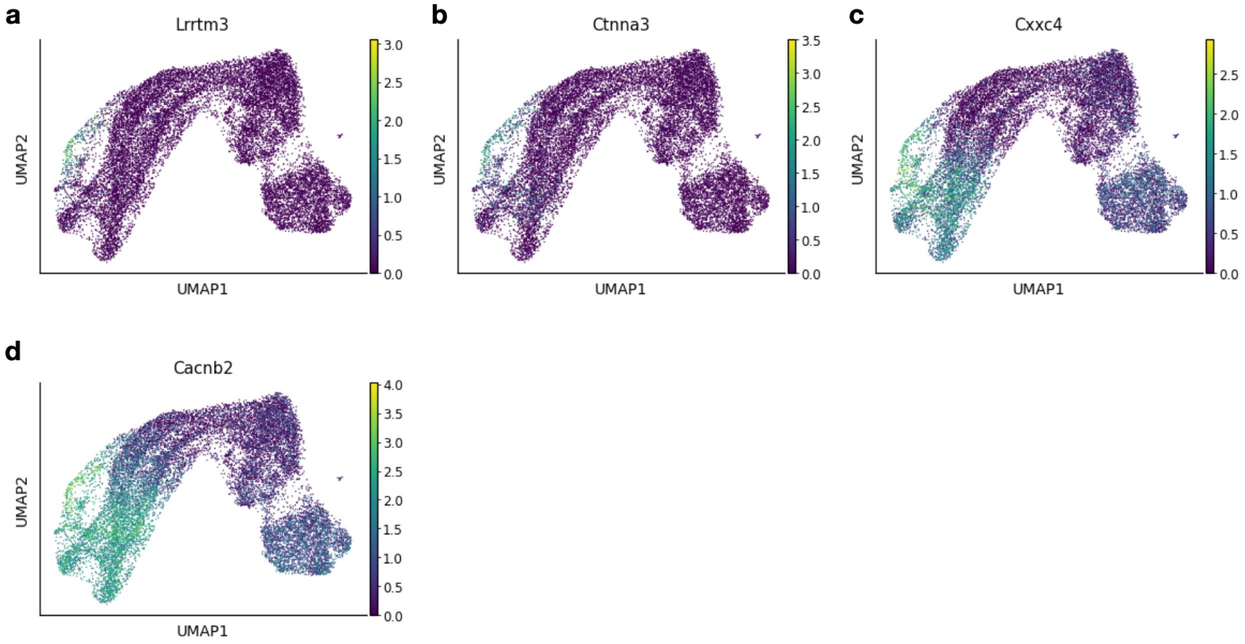


**Suppl. Fig. 29 | Expression of genes most associated with the marker peaks**

**a.** Processed (normalized and log1p-transformed) gene expression of *Lrrtm3*, the gene most highly correlated (Pearson correlation coefficient $0.16$) with the most significantly accessible peak (10-64082077-64082994, average log2 fold change of $3.1$) for the epsilon progenitor population (Fig. 5l). **b.** Processed gene expression of *Ctnna3*, the second highly correlated gene ($0.17$) with the marker peak of the epsilon progenitor population. **c.** Gene expression of *Cxxc4* (peak-gene correlation $0.16$), associated with the most significant peak (2-10757655-10758458, average log2 fold change of $2.8$) in the Fev+ delta-1 population (Fig. 5n). **d.** *Cacnb2* is the gene most highly correlated ($0.09$) with the most significantly accessible peak (3-137328983-137329833, $3.55$) in the Fev+ delta-0 population (Fig. 5m).

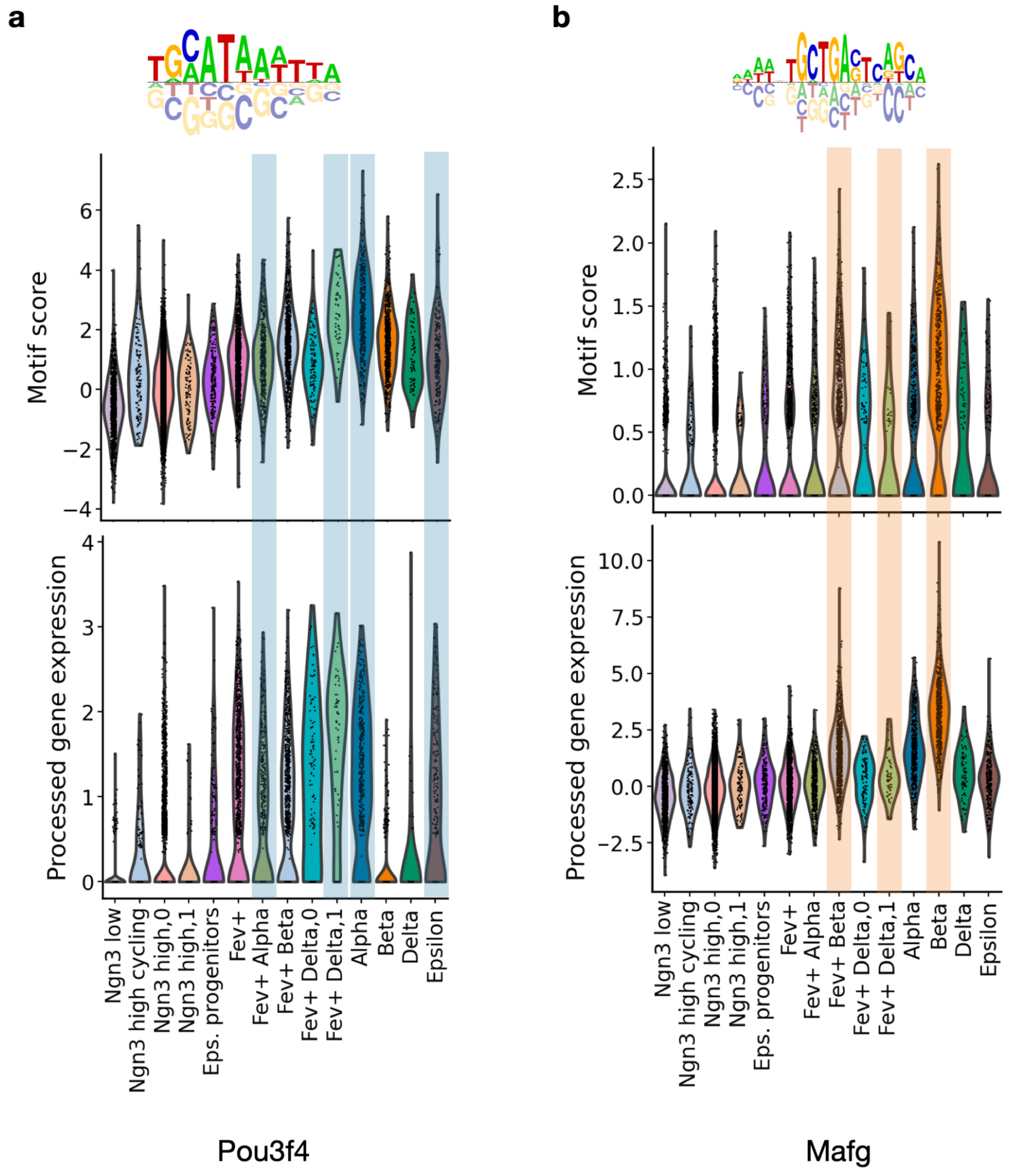


**Suppl. Fig. 30 | Alpha and beta motif activity calculated with moscot.time**

**a.** Motif with cisBP identifier M03318_2.00, which we identified as alpha cell marker motif using moscot.time and differential motif activity test. The upper plot shows the motif score computed by ChromVar, while the lower one shows the processed (normalized, log1p-transformed) gene expression of the associated gene *Pou3f4* (Methods). Alpha cells and their conjectured progenitors are underlaid in blue. For a motif to be active both the motif activity score and the gene expression should be high. **b.** Motif with cisBP identifier M08835_2.00, which we identified as a marker motif for the beta cell population. Again, direct beta cell progenitors are underlaid.

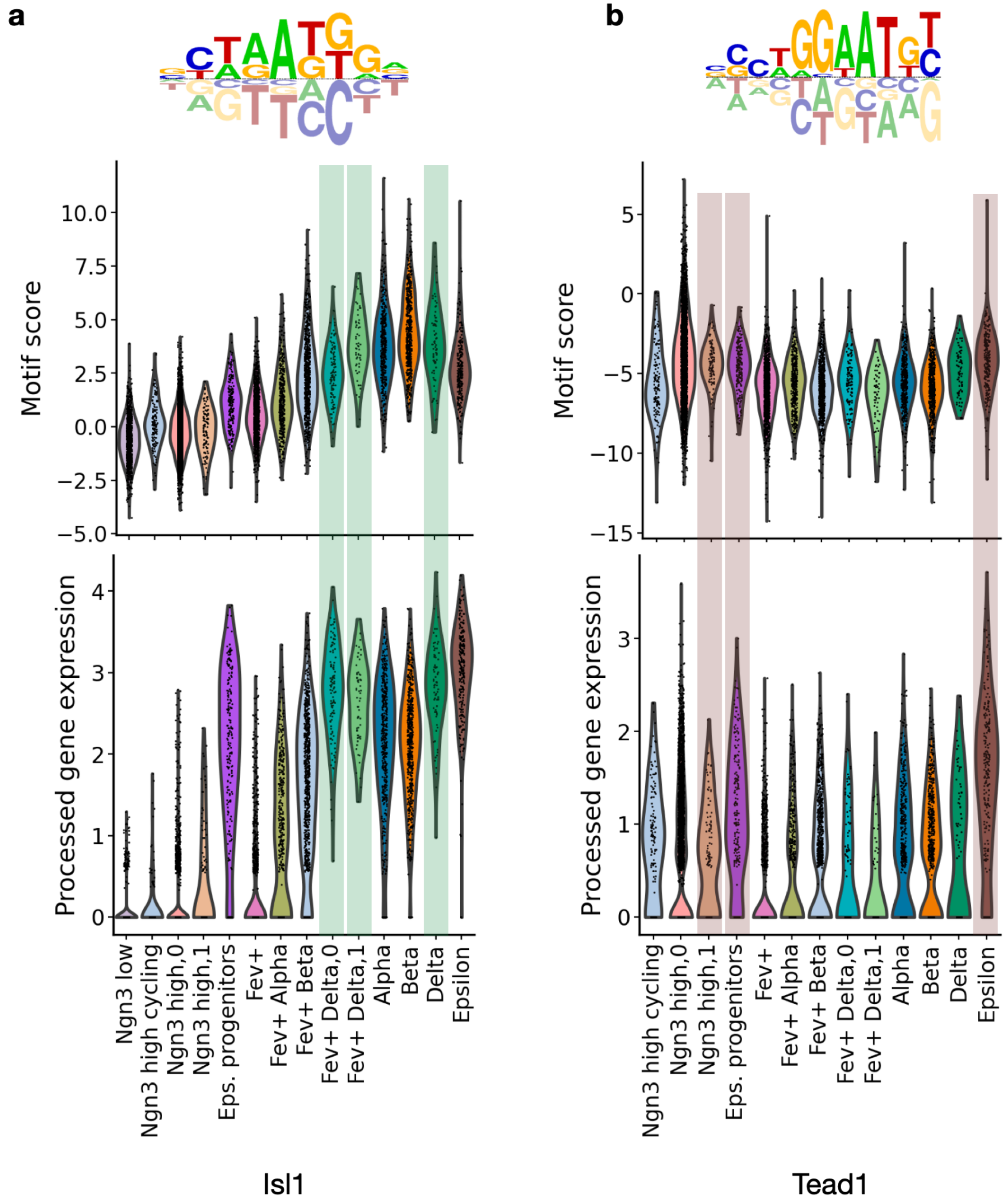


**Suppl. Fig. 31 | Delta and epsilon motif activity calculated with moscot.time**

**a.** Motif with cisBP identifier M09209_2.00, a marker motif identified for the delta population. For a motif to be active both the motif activity score (top) and the gene expression (associated transcription factor, here *Isl1*) should be high. Delta cells and their conjectured progenitors are underlaid in green. **b.** Motif with cisBP identifier M09438_2.00, which we identified to be a marker motif for the epsilon population. The motif is associated with the *Tead1* transcription factor (Methods).
